# Nature-Inspired Chemical Probes for in-cell Labeling of Lipidized RNA and Identification of New Regulatory Enzymes

**DOI:** 10.1101/2022.05.17.492276

**Authors:** Hongling Zhou, Yuanyuan Li, Jingwen Zhang, Li Wang, Ya Ying Zheng, Thomas J. Begley, Jia Sheng, Rui Wang

## Abstract

RNA modifications play pivotal roles in numerous cellular processes and human diseases. In addition to well-studied methyl-based modifications, hydrophobic prenyl-modifications have also been found in many RNA species. Here we report two chemical labeling strategies for tagging lipid-modified RNAs by taking advantage of a natural SelU-mediated tRNA geranylation process and the special reactivity of prenyl-groups. We synthesized a series of ‘clickable’ geranyl-pyrophosphate analogs and identified two candidates for indirect RNA labeling using a two-step process, *a*zidation-and-*c*lick-*t*agging of *flu*orescent dyes, namely ACT-Flu. We also developed a direct *m*etabolic *i*ncorporation and *b*iorthogonal tagging (MIBT-Tag) method based on the Ene-ligation of prenyl-groups. Both methods have been successfully applied to in-cell RNA labeling and the identification of new proteins associated with the geranylation process through proteomic and bioinformatic studies. These biochemical toolsets enable further *in vivo* applications to study prenylation pathways and monitor their status in both healthy and diseased cells.

## Introduction

Nucleic acids are fundamental building blocks in all life forms and play critical roles in cell proliferation, differentiation, migration, and apoptosis.^1-2^ RNAs are frequently decorated with more than 170 chemical modifications in both coding and non-coding regions to modulate their structure and function.^3-6^ RNA modifications, including methylation, acetylation, phosphorylation, glycosylation, palmitoylation and prenylation based changes, can not only diversify RNA stucture but also play critical functional roles in normal cell growth and development and regulating responses to environmental stress.^5^ Furthermore, chemical groups that modify the canonical nucleobases have shown to be involved in regulating the interactions of RNAs with other biomacromolecules including proteins, lipids, sugars and other forms of nucleic acids.^7-9^ Extending the networks of interaction, dysfunctional RNA modifications are linked to a wide range of human diseases including developmental disorders and cancers, as well as viral infections. Some of the most prevalent chemical modifications such as m^6^A and m^5^C have been extensively investigated and characterized on various RNAs species by many epitranscriptomic leaders and experts^10-21^ due to their high occurrence frequency and the crucial functions they play in biological processes.

In 2009, Liu and coworkers discovered a relatively large hydrophobic modification on the wobble position of some specific bacterial tRNAs.^22-23^ This C_10_H_15_ lipid moiety was identified as the geranyl group attached to the sulfur atom at position 2 of uridine (ges^2^U, **Fig.1** compound 1, 2 and 3).^24-27^ This geranyl group modifies about 0.4% of tRNAs specific for lysine, glutamine and glutamic acid in *Escherichia coli, Enterobacter aerogenes, Pseudomonas aeruginosa, and Salmonella enterica* serovar *Typhimurium*. Further studies revealed that the geranyl modification enhances tRNA translation fidelity and decreases frameshifting errors.^24^ Our following biophysical studies demonstrated that ges^2^U preferentially forms a base pair with G over A, with a significant enhancement in base pairing descrimination and codon recognition.^26^ It has also been proposed that the geranylated tRNA might be a transient intermediate in the selenation process, which is another important tRNA modification pathway in bacteria.^27^

In general, lipid-like hydrophobic prenyl groups, including geranyl, farnesyl, and geranylgeranyl, have been found to modify both proteins and RNAs with crucial functions.^28-30^ Many natural products such as caryophyllene^31^ and protein enzymes such as KRAS^32^ are all prenylated. Farnesyl and geranylgeranyl groups on cysteine in the CAAX motif in RAS super family proteins are essential in signaling transduction as they anchor the proteins to the cell membrane.^32^ The biological pathways and regulatory networks of these lipidized residues are relatively clear. Although it is known that geranylation in ges^2^U is mediated by the 2-selenouridine synthase (SelU), a conserved enzyme necessary for optimal bacterial growth, our understanding of the basic mechanisms that regulate geranylation process and its functions remains very limited. Specifically, the regulatory networks of geranylation with other reader, writer, and eraser enzymes remain largely unknown. Biochemical tools that enable the labeling of lipidized RNAs will allow for the analysis of hydrophobic modifications during normal and diseased biological processes. In addition, incorporating similar lipid analogs into tRNAs will provide a useful platform for the development of new molecular tools to label, detect, and isolate specific tRNAs for *in vitro, in situ* and *in vivo* applicaitons.^33-34^ To provide a toolset to fill in these knowledge gaps, we engineered two new methods, including an indirect two-step procedure called azidation-and-click tagging of fluorescent dye (ACT-Flu), and a direct metabolic incorporation and biorthogonal tagging (MIBT-Tag). These tools and methods can be used to analyze and exploit lipid modified RNAs, as well as to identify their interacting enzymes in living cells by taking advantage of the unique chemical reactivity of the prenyl groups ^35-37^ (overview in **Fig. 1**).

**Fig. 1.**
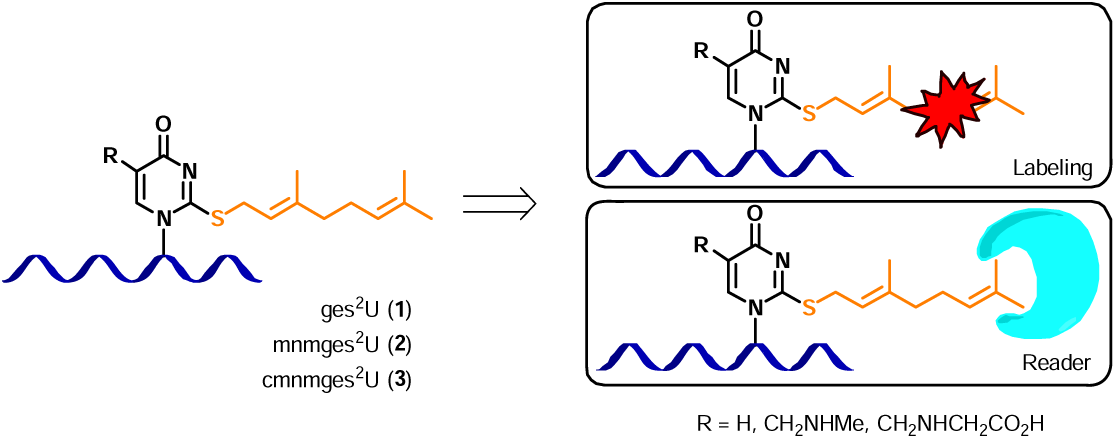
Labeling of geranylated RNA and identifying “reader” proteins. *Left:* The chemical structure of the geranyl modification on tRNA (represented as a ribbon). *Right:* Schematics of the fluorescent labeling on geranylated RNA and its reader proteins. R = H (ges^2^U, **1**), 5-methylaminomethyl (mnm^5^ges^2^U, **2**), or 5-carboxylmethyl amidomethyl (cmnm^5^ges^2^U, **3**).

## Results and Discussion

### Synthesis of geranyl pyrophosphate analogs

To validate the feasibility of geranyl tRNA labeling, we first synthesized a series of analogues including geranyl pyrophosphate variants **Ge2** and **Ge3**, a diphenylketone analogue **Ge4**, azido analogues **Ge5** and **Ge6**, and a DBCO derivative **Ge7** (**Fig. 2A, Schemes S1-5**). The pyrophosphate **Ge4** was designed as a photo-crosslinker, and all the other pyrophosphates were designed to mimic the natural geranyl pyrophosphate substrate Ge1. For the synthesis, **Ge1, Ge2**, and **Ge3** were all made from the corresponding bromide, and the crude products were purified by preparative HPLC. Synthesis of **Ge6** followed a two-step protocol. First, 3-azidopropylamine was reacted with 2-bromoacetyl bromide to yield bromide intermediate in 90% of yield, which was further reacted with hydrogen pyrophosphate to give **Ge6**. The analogs **Ge4, Ge5**, and **Ge7** were synthesized from geranyl acetate. Selenium dioxide and tert-butyl peroxide were used to oxidize the terminal methyl group on allylic position of geranyl acetate to generate the hydroxy intermediate. For **Ge4**, the diphenylketone acid was reacted with the hydroxide to give the ester. Deprotection of the acetyl group, bromination using tribromophosphine, and a treatment with hydrogen pyrophosphate in a three-step domino process with final HPLC purification yielded pure **Ge4**. For the synthesis of **Ge5**, a key intermediate hydroxide was subjected to the treatment with mesyl chloride and subsequently with sodium azide to generate the azido-intermediate. The aforementioned three-step domino process were subsequently applied to produce final **Ge5**. Similarly, a key intermediate hydroxide was subjected to react with triphosgene and DBCO-amine, followed by the three-step domino process to generate **Ge7**. All the compounds were analyzed by mass spectroscopy, ^1^H NMR, and ^31^P NMR (**Fig. S1-33**), confirming the identity of the desired products. Like the natural **Ge1** [-6.85 (d, *J* = 22.68 Hz), −10.57 (d, *J* = 22.68 Hz)], the analogs **Ge5** and **Ge6** have characteristic peaks in the NMR spectra at −6.64 (d, *J* = 22.68 Hz), −10.44 (d, *J* = 22.68 Hz), −5.87 (d, J = 22.7 Hz), and −11.19 (d, *J* = 22.3 Hz), which are typical for pyrophosphates (**Fig. 2B**).

**Fig. 2.**
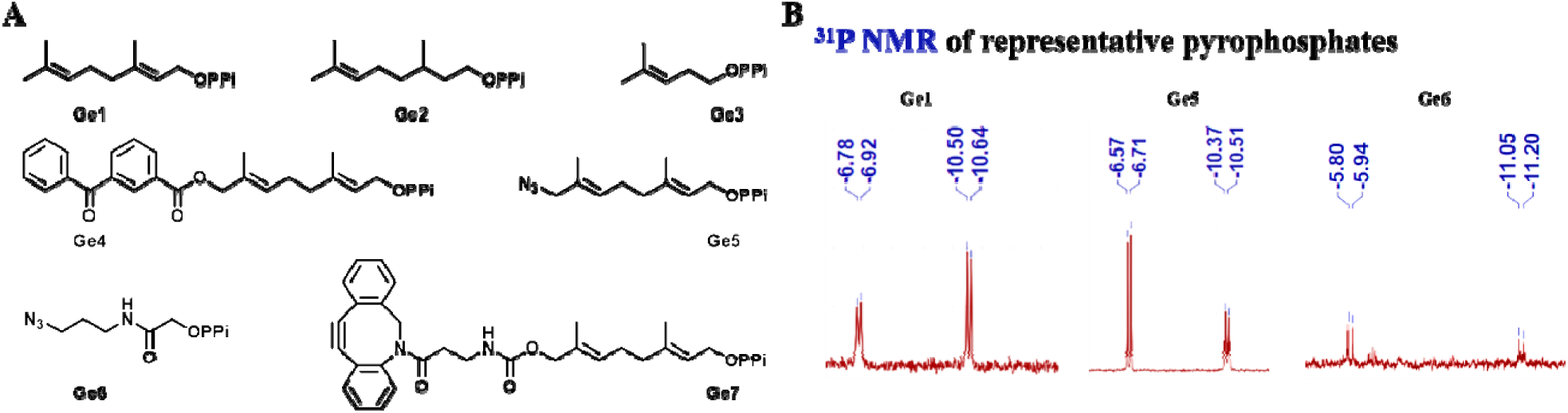
Chemical structures and ^31^P NMR spectra of geranyl pyrophosphate variants. (A) Geranyl pyrophosphate variants **Ge1-7**. (B) Example ^31^P NMR of geranyl pyrophosphate variants **Ge1, Ge5**, and **Ge6**. Experimental NMR conditions: CD_3_CN, 162 MHz, Varian instrument.

### SelU expression and modeling of substrates binding

Next, we expressed and purified SelU, the enzyme that installs the geranyl modification on tRNAs.^38^ Immunoblotting analysis confirmed the successful expression of SelU-His6 with the His-tag used for the affinity column purification (**Fig. S44, Table S2-3**). To investigate the substrate recognition criteria and better understand the geranylation process, we sought to predict the 3D structure of SelU using AlphFold (DeepMind). The protein is predicted to adopt an overall α helical structure (**Fig. E1-A (extended data)** and **Fig. S45-46**). SelU is known to contain a rhodanese domain, which has been shown to be the key working domain for both geranylation and selenation processes,^38^ and a P-loop domain with a Walker A motif, which is present in many ATP- and GTP-binding proteins and is also involved in substrate binding.^39-40^ Within the rhodanese domain, the two key residues Cys97 and Gly67 have been proven to be critical in both substrate binding and geranylation activation. Therefore, the detailed interacting network of Cys97 and Gly67 with the neighboring residues was analyzed (**Fig. E1-B**,**C**), which is consistent with the reported experimental data.^38^ With this apo-model in hands, we conducted the molecular docking studies with the synthesized geranyl-pyrophosphate ligand analogs. The data indicated that the predicted ligand binding pocket of SelU can well accommodate **Ge1, Ge5**, and **Ge6**, and the azido analogues are ideal substrates for geranylation by SelU (**Fig. E1-E** and **E1-F**). Although more accurate SelU-ligand interactions remain elusive with high-resolution complex structure studies, this AI-based prediction and docking strategy provided some general guidance for the mechanistic studies and further ligand design based on the structurally unknown proteins.

### Labeling of tRNA via geranyl derivatives

With these ‘clickable’ tags being recognized by SelU, we further conducted the fluorescent labeling of tRNA^Gln, Glu, Lys^ in the presence of the ligands **Ge5, Ge6, Ge7** and the purified SelU (the detailed processes are shown in **Scheme S7-8, Fig. S47-53**, and **Table S4**). When the reaction was performed in the presence of azido-pyrophosphate **Ge5**, efficient fluorescent signal of tRNA was observed with DBCO-Cy5 treatment, as shown in **Fig. 3A** and **Fig. S48**. SYBR green staining was used to validate the presence of RNAs. Similarly, **Ge6** is also a substrate for SelU, and the subsequent click reaction with DBCO-Cy5 resulted in fluorescent tRNAs (**Fig. 3B** and **Fig. S49**).

**Fig. 3.**
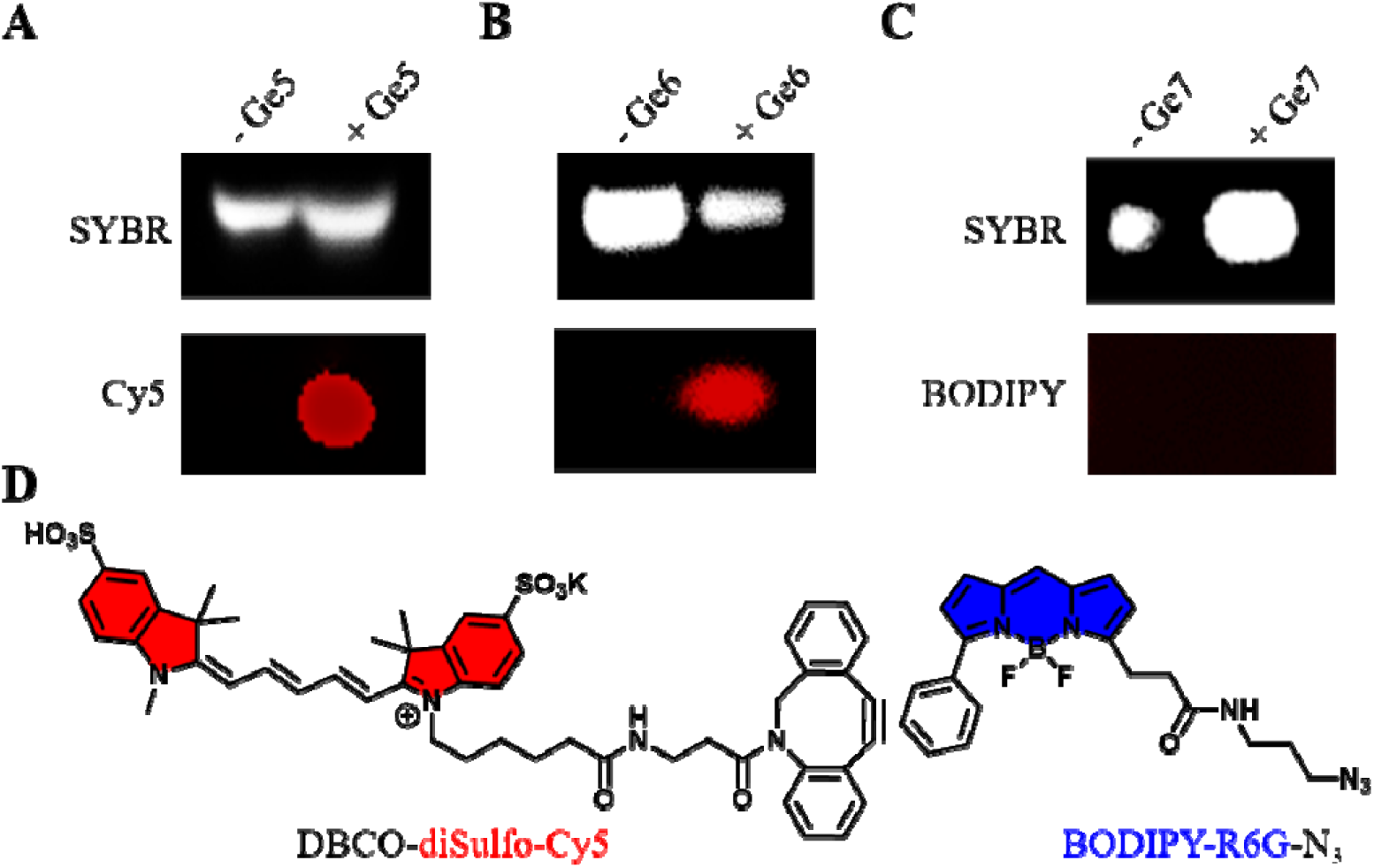
Labeling of tRNA using geranyl pyrophosphates Ge5, Ge6 and Ge7 by click reaction. (A) tRNA^Gln, Glu, Lys^ modified with pyrophosphate **G5** (10 *μ*M) was visualized via strain-promoted alkyne-azide cycloaddition (SPAAC) reaction in the presence of DBCO-diSulfo-Cy5 (10 *μ*M). (B) tRNA^Gln,Glu,Lys^ modified with pyrophosphate **G6** (10 *μ*M) was visualized via SPAAc reaction with DBCO-diSulfo-Cy5 (10 *μ*M) along with SYBR staining. (C) tRNA^Gln, Glu, Lys^modified with pyrophosphate **G7** (10 *μ*M) was visualized via SPAAC reaction using BODIPY-R6G (10 *μ*M). The to panel (SYBR staining) loaded with equivalent amounts of tRNA^Gln,Glu,Lys^ with and without geranyl pyrophosphates (Ge5-7). (D) The chemical structures of DBCO-diSulfo-Cy5 on the left and BODIPY-R6G on the right. All in-gel fluorescence assays were analyzed on 3% PAGE gel. At least three duplicates were performed, and *Zeiss Confocal Image* was used to measure in-gel fluorescent intensity.

After the labeling, the tRNAs were further treated with RNase T1 and analyzed by LC-MS/MS to confirm the presence of the fluorescent groups on nucleosides. The mass fragmentation results (**Fig. S52-54**), including 3398.8 (CCCUUUCACG, **Ge5**), 3268.8 (ACUUUUAAUCAAUUG, **Ge6**) and 3216.0/3293.0 (UUUUUGAUACCG, **Ge1**), were assigned as the corresponding RNA fragments cleaved at the G sites, which are the target cleavage sites of the RNase T1. All these data supports the idea that our ligand Ge5 and Ge6 can be successfully transferred to tRNA as tags for further fluorescence labeling. On the other hand, **Ge7** bears a large DBCO moiety and most likely interferes with the SelU binding, and as a result the fluorescent click labeling using BODIPY-R6G-N_3_ failed (**Fig. 3C, Scheme S5** and **Fig. S51**). Similarly, we also evaluated the labeling feasibility of ligands **Ge2** and **Ge3**. After the click reaction and RNase cleavage, both HPLC and MS-spec data showed no modified RNA residues (**Fig. S34-43** and **Table S5-7**), indicating that neither of these two substrates can be recognized by SelU, probably due to the higher flexibility of **Ge2** and shorter chain length of **Ge3**.

There are applications for this two step indirect ACT-Flu labeling in eukaryotic and human disease conditions. It has been recently shown in budding yeasts that the levels of thiolated wobble uridines change in response to hydroxyurea.^41^ Similarly, in human colorectal cancer cells, the thiolated-tRNA levels were found to change in response to the paromomycin treatment.^42^ In addition, it has also been reported that the human U34 writers linked to mcm^5^s^2^U provide resistance to targeted therapy in melanomas.^43^ Therefore, our labeling technology could be applied to study and monitor these thiolated tRNA levels in different cellular states and environmental stresses.

In a brief summary, through these experiments, we have identified two pyrophosphate analogs, **Ge5** and **Ge6**, that were suitable for SelU-mediated tRNA geranylation and fluorescent labeling. This two-step process, azidation and click tagging of fluorescent dye (ACT-Flu), could have great potential for in-cell tRNA labeling and detecting.

### Direct metabolic incorporation and biorthogonal tagging (MIBT-Tag) of prenyl-containing RNAs

In the meantime, we also accessed the feasibility of a direct labeling of prenyl-containing RNAs. Since some bacterial tRNAs contains geranyl groups on the wobble position of ^34^U, we investigated if a 4-phenyl-1,2,4-triazoline-3,5-dione (PTAD) activated fluorescence dye molecule could be directly attached to these terpene terminals through a known Ene-reaction, where PTAD reacts with conjugated diene group ^35-37^ (**Fig. 4A**). Indeed, as shown in **Fig. 4B**, a red fluorescent signal was observed when the total tRNAs isolated from *E.* Coli were treated with 10 μM of PTAD-DBCO-Cy5 (structure shown in **Fig. 4C**), while there was non-red-fluorescence in the absence of this fluorophore. SYBR staining confirmed the equivalent amount of tRNAs loaded in gel. We next applied this fluorophore probe directly to the ges^2^U-tRNA to test the labeling specificity. The labeled tRNAs were analyzed by MALDI-TOF (**Fig. 4D**). Mass peaks detected at 25724.9355 and 25417.4251 amu corresponded to PTAD-DBCO-Cy5-labeled tRNA^Gln,Glu,Lys^. In brief, the direct labeling of geranylated tRNA by the fluorescent probe PTAD-DBCO-Cy5 could be achieved based on the *in vitro* Ene-ligation.

**Fig. 4.**
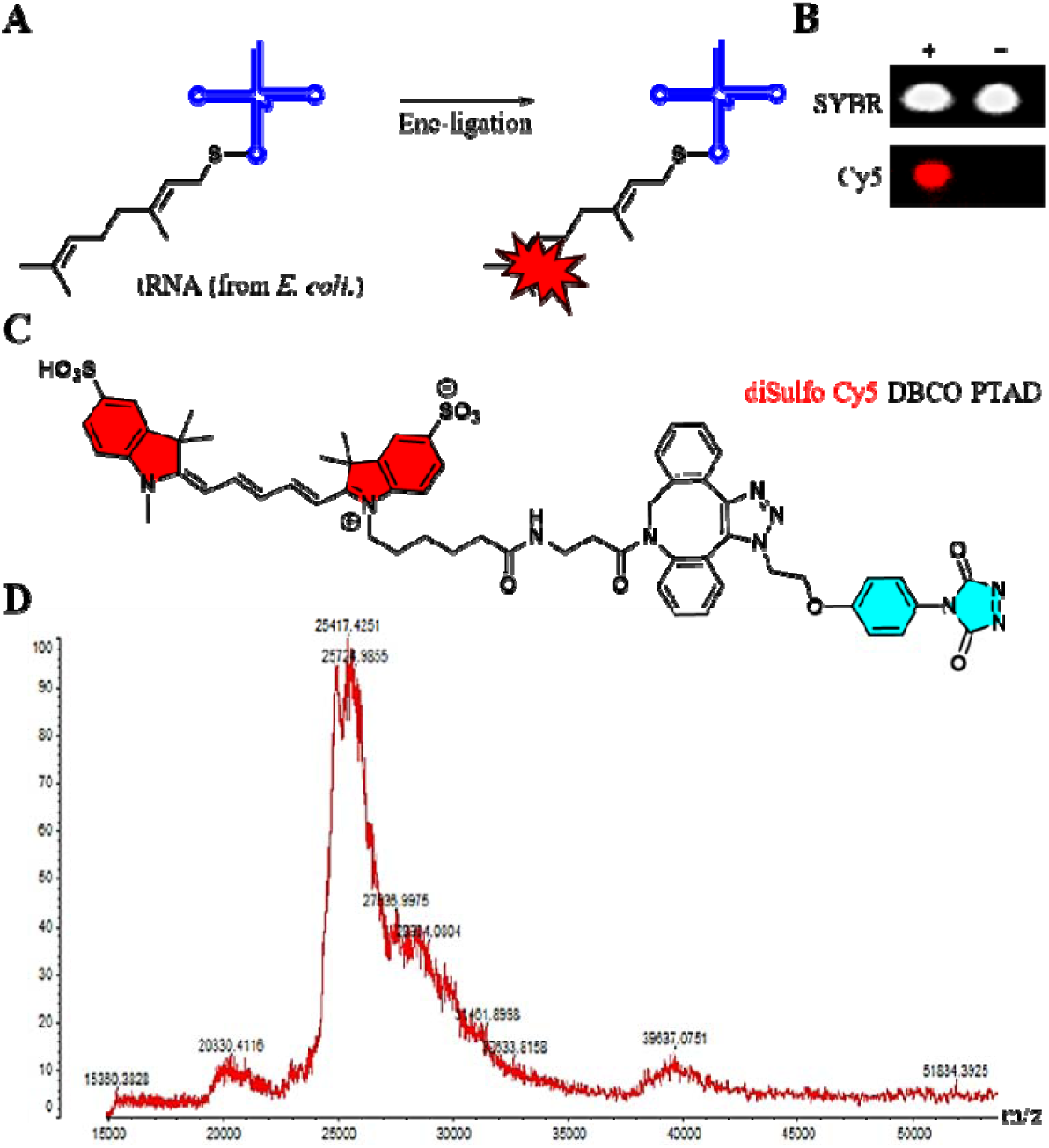
Direct labeling of tRNAs in E. coli. with diSulfo-Cy5-DBCO-PTAD. tRNAs (1.0 mg from *E. coli*., *Roche*) were reacted with diSulfo-Cy5-DBCO-PTAD (1.0 mM) for 5 min. 3% Agarose gel was used to visualize the geranylated tRNAs and confirmed by MALDI-TOF. Total tRNAs reacted directly with the probe diSulfo-Cy5-DBCO-PTAD (10 *μ*M). (A) Fluorescent labeling of geranylated tRNAs by Ene-ligation. (B) tRNA^*Gln, Glu, Lys*^ extracted from *E. coli*. reacted with pre-established probe followed by SPAAC reaction with diSulfo-Cy5-DBCO-PTAD (10 μM). The top panel (SYBR staining) showed the equivalent amounts of tRNA^*Gln, Glu, Lys*^ with and without probe were loaded. (C) Chemical structure of diSulfo-Cy5-DBCO-PTAD dye (*in-situ* generated by NBS). (D) Illustrated the HRMS mass 25724.9355 (cnmges2U-tRNA^Glu/Lys^, 2‰) and 25417.4251 (cmnmges2U-tRN^Gln^, 2‰). The mass spectra were analyzed on a *Shimadazu AXIMA Performance*-MALDI TOF/TOF.

### In-cell labeling of prenylated RNAs

To further apply our methodologies to the in-cell tRNA labeling, we constructed the *5s-pCMV3* (6333 bp, contains *5S rRNA*) and *SelU-pCMV-Myc-EGFP* (5593 bp, contains *E. coli* SelU tagged with EGFP) plasmids and tested the *in vivo* fluorescent labeling of geranylated tRNAs with pyrophosphates **Ge1** coupled with the direct labeling, and **Ge5** and **Ge6** with indirect ACT-Flu process (**Fig. 5A**). Specifically, the direct probe PTAD-DBCO-Cy5 was used to selectively tag the prenylated tRNAs catalyzed by *SelU* in the presence of natural geranyl pyrophosphate **Ge1** in HEK293T cell line. This direct labeling of prenylated RNAs turned out to be very efficient. The confocal image clearly showed the red fluorescence when transfected *SelU, 5s* (tRNA^Lys^) with pyrophosphate **Ge1**, as well as probe PTAD-DBCO-Cy5 (**Fig. 5B** and **S57**). Alternatively, in order to test the sequential two-step ACT-Flu approach, we also co-transfected *SelU* with pyrophosphate **Ge5** or **Ge6**, as well as the probe DBCO-Cy5 into HEK293T cell line. The resulting confocal image also displayed strong red florescence to indicate the successful labeling of the prenyl group (**Fig. S58**). When the *SelU-pCMV-Myc-EGFP-5s* plasmid was transfected in the presence of **Ge6**, both red and yellow fluorescence could be observed (**Fig. 5C**). As the control, no red fluorescence signal was detected in the absence of either *SelU* or *5s*. These combined results demonstrated the successful in-cell labeling capacity of our two methodologies.

**Fig. 5.**
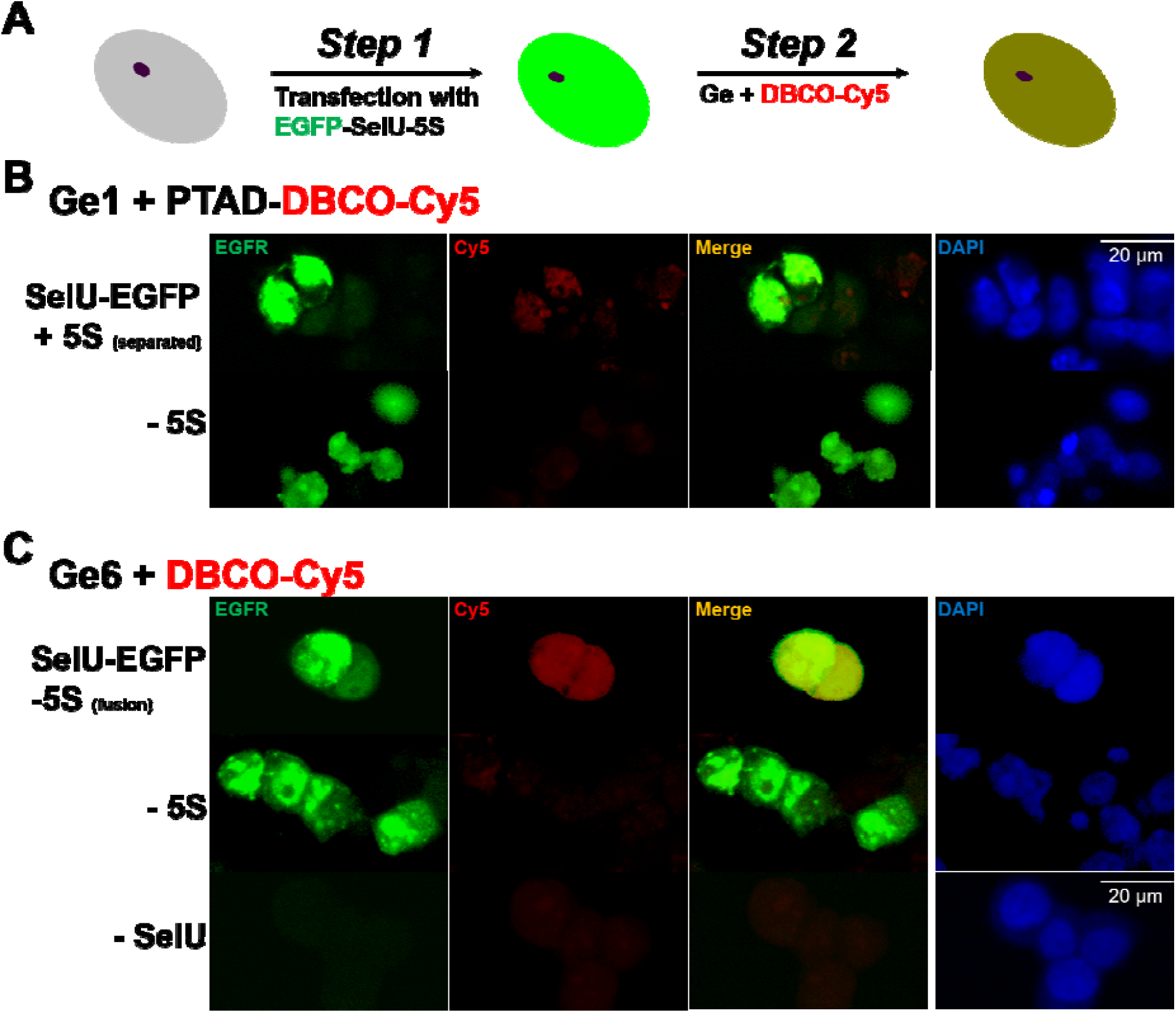
Fluorescence labeling of E. coli. tRNA^lys^-5S in 293T cells. (A) Overall illustrations of the fluorescence labeling of prenylated RNA *in vivo*. (B) Top panel: Transfection of fusion plasmid *SelU-EGFP* with transcript 5S in the presence of pyrophosphate **Ge1** (**1** mM), followed by the treatment of PTAD-DBCO-Cy5 probe (50 *μ*M). Bottom panel: Transfection of fusion plasmid *SelU-EGFP* without transcript 5S in the presence of pyrophosphate **Ge1** (**1** mM) and PTAD-DBCO-Cy5 treatment. Green fluorescence (left), Cy5 fluorescence (second), merged (third), and DAPI fluorescence (nuclear staining; right). (C) Transfection of fusion plasmid *SelU-pCMV-Myc-EGFP-5s* in the presence of pyrophosphate **Ge6**, followed by the treatment of DBCO-Cy5 probe (5 *μ*M) via click reaction. Imaging of HEK293T cells harboring the *SelU-pCMV-Myc-EGFP-5s plasmid* (top line, both red and green fluorescence), *SelU-pCMV-Myc-EGFP* plasmid (second line, only green fluorescence in the absence of transcript 5s); *pCMV-Myc-EGF* (bottom line, no green/red fluorescence in the absence of SelU). Green fluorescence (left), Cy5 fluorescence (second), merged (third), and DAPI fluorescence (nuclear staining; right). DAPI = 4’,6-dianidino-2-phenylindole. Scale bar: 20 μm.

### Extended application of direct labeling method: identification of new prenyl-regulatory proteins

Many different reader proteins as networks have been demonstrated to recognize and regulate a wide range of RNA modifications including m^6^A and m^5^C. To determine how geranylated tRNAs are regulated and further expand the biological applications of our molecular probes, we designed and synthesized three new probes **1-3** bearing the biotin and PTAD functionalities (**Fig. 6A** and **Scheme S11-13, Fig. S59-70**) that can selectively target and pulldown both geranylated tRNAs along with their reader proteins utilizing Ene-ligation chemistry. The probe 3 is a three-necked probe bearing an additional photo-reactive three-membered diazirine group as a carbene generate to crosslink with the neighboring amino acid residues. Using the *E. coli* cell lysate, probe **1-3** (10 *μ*M) were incubated for the optimized time respectively. By employing streptavidin magnetic beads pulldown assays, we have captured several latent proteins candidates. These proteins were then subjected to conventional LC-MS/MS proteomic analysis. We found that the protein size around 30-33 KDa in gel could show relatively weak but common signals using all the three probes (**Fig. 6B-D** and **Fig. S71-73**). The size similarity of these captured proteins also conveyed a key message about the binding specificity of geranylated RNAs to their reader proteins. The subsequent proteomic studies revealed that several of these proteins have been well identified (**Table S13-15**). Furthermore, we used bioinformation data STRING to analyze the according regulatory network involving these proteins. As shown in **Fig. 6E**, among the 44 proteins in the network, 18 have been co-found when using both probe 1/2 and the three-headed probe 3 (the dark green overlapping areas), suggesting that these 18 proteins are potentially the regulatory candidates. Within them, 12 have been identified and included in the STRING database, and the other 6 might be new players in this network. In brief, the functional probes 1-3 can be used to identify the geranylation-based regulatory proteins. The six new candidates, which are mainly the outmembrane proteins, remain to be further identified and elucidated in our currently onging works.

**Fig. 6.**
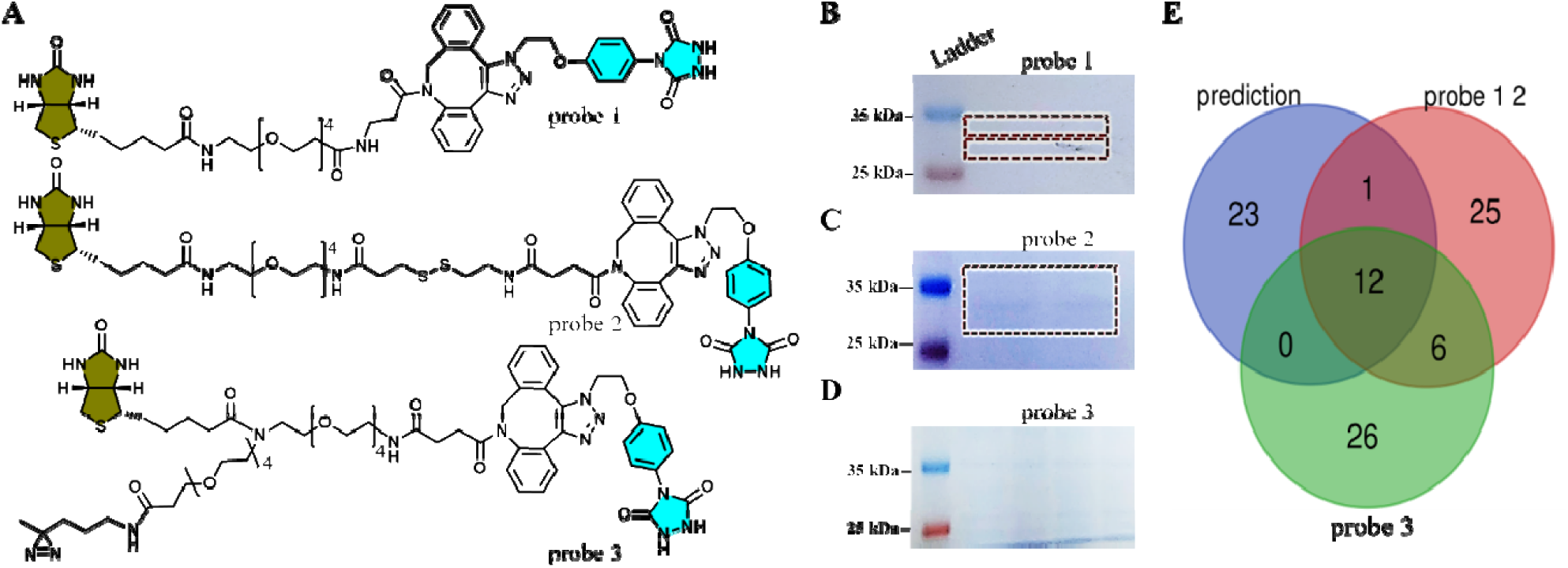
Identification of tRNA geranylation “Reader” proteins using probes 1-3 and direct lableing. (A) Chemical structures of the probes 1-3. (B)-(D) Immunoblotting of the pulldown assays using probes 1-3, respectively. (E) Venn diagram analysis of captured reader proteins from the pulldown assays or from prediction. Venn diagram representing geranylated tRNA regulatory proteins in our study. According to STRING database, proteins identified as tRNA geranylation reader proteins were categorized into “probe 1 or 2”, “probe 3” and “Prediction”. 18 proteins existed in the captured proteins using probe 1 or 2 and probe 3. Among them, 12 proteins are known (STRING database analysis) and 6 proteins have been identified as new reader candidates.

## Conclusion

In conclusion, we report two chemical labeling strategies for the lipid-modified RNAs by taking advantage of the natural SelU-mediated tRNA geranylation process and the special chemical reactivity of the prenyl-groups. We synthesized a series of ‘clickable’ geranyl pyrophosphate analogs and identified two candidates **Ge5** and **Ge6** that are suitable for indirect in-cell RNA labeling through a two-step process, azidation-and-click tagging of fluorescent dye, namely ACT-Flu process. Both fluorescent microscopy and molecular simulation studies confirmed that the two ligands could serve as ideal SelU substrates in the tRNA geranylation process, representing a new toolset for detecting and monitoring the 2-thiouridine residue, which is an important biomarker of bacteria infections and cancers. In addition, we developed a direct metabolic incorporation and biorthogonal tagging (MIBT-Tag) method by taking advantage of the PTAD-based Ene-ligation chemistry of the prenyl group. Both methods have been successfully applied to the in-cell RNA fluorescent labeling. Furthermore, by coupling the three new PTAD based functional probes with the direct labeling and proteomic studies, we have identified a series of new enzymes that are associated with the geranylation process. The biological importance of prenylation process has been increasingly appreciated with recent studies of OSA1, the double stranded RNA sensor that activates RNAase L, indicating that the OSA1 mediated prenylation^44-45^ process protects patients from severe illness infected by SARS-CoV-2 virus. The biochemical toolsets developed in our work has important *in vivo* application potentials. They will not only shed light on the functions of prenylation modifications on gene regulation and RNA biology, but have biomarker applications specific to the levels of prenylation in both healthy and diseased cells.

## Methods

### General information

All chemicals were purchased from Sigma-Aldrich, Acros, Inno-chem, Macklin Inc, Energy Chemical.

### SelU expression, and purifications

#### Expression

Strain pET28b-SelU (BL21) was cultivated in Kan (kanamycin) LB overnight at 37 °C and transferred 2.0 mL of strain liquid into fresh 200 mL Kan LB, cultivated at 220 r.p.m./min, 37°C for about 3 h to OD = 0.6. Then the 200 mL culture was induced with 1 mmol/L IPTG at 220 r.p.m./min, 28 °C for 5 h.

#### Purifications

The His×6 Ni Gravity Column (1.0 ml) was equilibrated and all buffers was settled to the working temperature. The column was then washed with 5-10 column volumes of His×6 Ni Equilibration Buffer (50 mM sodium phosphate, 6 M guanidine-HCl, 300 mM NaCl, 20 mM imidazole; pH = 7.4). The clarified sample was added to the column and the top stopper was carefully connected to the top of the column. The target protein was allowed to bind by slowly inverting the column for 1 h. The column was installed in a vertical position and the resin was settled at the bottom of the column. The column was washed with 10 column volumes of His×6 Ni Equilibration Buffer followed by 10 column volumes of His×6 Ni Wash Buffer (50 mM sodium phosphate, 6.0 M guanidine-HCl, 300 mM NaCl, 40 mM imidazole; pH = 7.4). The target protein was eluted with approximately 10 column volumes of Elution Buffer (50 mM sodium phosphate, 6 M guanidine-HCl, 300 mM NaCl, 300 mM imidazole; pH = 7.4) and collect 1 mL once in a tube. The sample was analyzed and identified by 5% agarose gel electrophoresis.

### NMR spectroscopy

Measurements were carried out using ^1^H, ^13^C NMR, and ^31^P NMR. Chemical shifts (δ) were reported in parts per million (ppm) and referenced to the solvent signals. Data are reported as follows: chemical shift, multiplicity (s=singlet, d=doublet, t=triplet, and m=multiplet) and coupling constants (in hertz). Bruker AM-400 spectrometer (400 MHz) and a Bruker AVANCE NEO 600 (600 MHz) was used for NMR data collection and spectral interpretation.

### General synthetic procedures for pyrophosphates (Ge1-7)

To a stirred solution of (*E*)-1-bromo-3,7-dimethylocta-2,6-diene (50.0 mg, 0.220 mmol) in CH_3_CN (1.5 mL) was slowly added tris(tetra-*N*-butylammonium) hydrogen pyrophosphate (200 mg, 0.264 mmol). The mixture was stirred for 5 h at room temperature under nitrogen atmosphere. The crude product was purified by HPLC (see below **Table E1**) and lyophilized to dryness.

### Molecular Docking Study of SelU with pyrophosphates Ge

The molecular docking studies were performed by using AutoDock Vina 1.1.2.^Ref.M1^ The ligand sites were detected by using AutoLigand,^Ref.M2^ and the ligands (the pyrophosphate **Ge1/Ge5/Ge6**) site containing Cys97 and Gly67 was used to define the binding pocket by establishing a grid box centered on X: −26.5 Y: 2.6 Z: 7.8 Å with the dimensions of X: 26.9 Y: 26.0 Z: 19.1 Å. The magnesium ion was first docked into the binding pocket. The docked position of the magnesium ion with the lowest docked energy was selected to prepare a protein-ion complex. Then each ligand was docked into the binding pocket with the presence of the magnesium ion. The docking poses of ligands were also selected for subsequent analyses according to the lowest docked energies.

## General experimental procedures for SelU-mediated geranylation Enzymatic fluorescent labeling analysis with pyrophosphates Ge1-7

Partially purified SelU-His×6 (15 μg, ca. 0.35 nmol), 20 μg (ca. 3.74 nmol) tRNA and geranyl pyrophosphate ammonium salt (Sigma, 5 equiv.) was dissolved in 100 μl of buffer containing 10 mM Tricine-KOH, pH = 7.2, 0.2 mM dithiothreitol (DTT), and 100 mM MgCl_2_, and the sample was incubated at 25 °C for 24 h. The Ge-tRNA product was monitored by RP-HPLC using a Kinetex C18 column (5 μ, 100 A, 150 × 4.6 mm; in the gradient of *Buffer B*: 0-10 min 0% B; 10-40 min 0-35% B; 40-45 min 35-100% B; 45-50 min, 100% B; 50-55 min, 100-0% B; 55-60 min, 0% B. *Buffer A*: 0.1 M CH_3_CO_2_NH_4_; pH = 6.8. *Buffer B*: 0.1 M CH_3_CO_2_NH_4_ with 40% CH_3_CN; the collected fraction was desalted and the geranylated product was tested by fluorescent labeling or MALDI-TOF MS.

### With pyrophosphates Ge2,3,7

RNase T1 is an endoribonuclease that specifically degrades single-stranded RNA at G residues. It cleaves the phosphodiester bond between 3’-guanylic residue and the 5’-OH residues of adjacent nucleotide with the formation of corresponding intermediate 2’, 3’-cyclic phosphate. The reaction products are 3’-GMP and oligonucleotides with a terminal 3’-GMP. RNase T1 does not require metal ions for activity. The Ge-tRNA product was hydroxylated to give nucleosides with varied modifications, which were further analyzed by AB Sciex 4500, Wuhan biological sample 4500 Q-trap (*Wuhan Biobank Co*., *Ltd*).

### With pyrophosphates Ge1,5,6

SelU-catalyzed tRNA geranylation with pyrophosphates **Ge5-7** as substrates, the in-gel fluorescent imaging was performed therefore. In the fluorescent labeling studies when using **Ge5-6** and **Ge7** as substrates, DBCO-Cy5 was used for labeling of **Ge5** or **Ge6**, while BODIPY-R6G-N_3_ was used for labeling of **Ge7**. Confocal parameters are: Blue channel (E.X. 492 nm. E.M. 507 nm), red channel (E.X. 649 nm. E.M. 660 nm) and green channel (nm).

## Direct fluorescent labeling of prenylated tRNA by using PTAD-DBCO-Cy5 probe

tRNA (1.0 mg) isolated from *E. coli* using Zymo purification kit strictly following the instruction protocol was dissolved in PBS buffer (0.1 M, pH = 7.4) and the PTAD-DBCO-Cy5 probe (1.0 mM) generated by in-situ oxidation with *N*-bromosuccinimide (NBS, 1 M in DMF) was added in one portion at 4 °C. The mixture was incubated at 4 °C for 15 min in a sterling tube. And excess fluorescent probe was removed by 3K filter (Millipore) and the resultant nucleic acid was analyzed by in-gel fluorescence.

### Mass Spectra identification of prenyl-containing nucleic acid labeled with PTAD-DBCO-Cy5

tRNA^Glu^ from aforementioned procedure was further analyzed by MALDI-TOF/TOF mass spectra (*AXIMA-PerformanceMA, Qinghua University*). The detailed are presented below:

### MALDI matrix application

3-Hydroxypicolinic Acid (3-HPA, Sigma Aldrich, USA) was chosen as MALDI matrix for DNA detection. The matrix solution was prepared by dissolving 20 mg 3-HPA and 45 mg dihydrogen ammonium citrate (DHAC) in 1 mL mixture solution of 50% acetonitrile/50% water. A home-built inkjet printing device was developed to print an array of matrix droplets on sample. This device consisted of a piezoelectric inkjet print head for matrix solution ejection (*Fuji Electrics Systems Co*., *Ltd, Japan*) and an XY-motorized stage where the ITO glass was placed (*MMU-30X, Chuo Precision Industrial Co*., *Ltd, Japan*). A laboratory-made software was used to control the inkjet waveform and the movement of x–y stage. ITO glass with cells was firstly washed with 100 mM ammonium acetate solution to remove non-volatile salts which might affect ionization efficiency. After air drying, a 6×6 matrix array of circular regions ∼300 μm in diameter was printed on the ITO glass and imaged with fluorescent microscope (*DMI 4000B, Leica, Germany*). For quantitative analysis, DNA internal standard and matrix solution were loaded into different channels of the inkjet printing head. The internal standard was printed prior to the matrix on the same position.

### MALDI-MS analysis

MALDI-MS analysis was performed in an AXIMA Performance MALDI-TOF/TOF mass spectrometer (*Shimadzu Co. Ltd*., *Japan*). This instrument was equipped with a 337 nm nitrogen laser. ITO glass with cells was attached to the stainless MALDI plate by conductive tapes. Data were acquired in a linear negative mode and signals between m/z 20000–30000 were collected. Raster scans on cell surfaces were performed automatically using the mass spectrometry software (*Shimadzu Biotech*., *Japan*). The scan area was 200 μm × 200 μm with the sampling distance of 50 μm. For each coordinate, mass spectra resulting from 20 laser shots at 5 Hz were accumulated to obtain an average mass spectrum.

### *In vivo* **labeling of prenylated nucleic acids**

#### Construction of plasmids

*pCMV3-5s* (or *5s-pCMV3*, 6333 bp).

See Figure S56A for details.

*pCMV3-Myc-SelU-EGFP* (or *SelU-pCMV-Myc-EGFP*, 5593 bp).

See Figure S56B for details.

Merged *pCMV3-Myc-SelU-EGFP-5s* (or *5s-SelU-pCMV-Myc-EGFP*, 8197 bp).

See Figure S56C for details.

### HEK293T cell culture and plating

The HEK293T cells was washed twice with PBS, and trypsin was added for 1 min. Then 2.0 mL of DMEM medium was added into the mixed well, was centrifuged at 1,000 r.p.m for 5 min, and the supernatant was discarded. Fresh medium was added to resuspend the cell pellet and cells were seeded in a confocal dish at the density of 0.5 ×1 0^5^, the cells were gently mixed using pipette, and then placed in CO_2_ incubator.

### Transfection of HEK293T cells

150 mM sterile NaCl solution was prepared as a diluent for DNA and ***Vigofect*** *(Vigorous Biotechnology Beijing Co*., *Ltd*.). The culture medium in the confocal dish was then replaced with fresh complete culture medium before transfection, and the culture was incubated at 37 °C under 5% CO_2_ atmosphere. The pCMV-Myc-SelU-GFP group, pCMV-Myc-SelU-5s-GFP and **Ge1** or **Ge5** or **Ge6** group were settled as control groups.

### Configure the transfection working solution

First, 2.5 μg pCMV-Myc-SelU-GFP or pCMV-Myc-SelU-5s-GFP was added into the diluent (total 100 μl) and stored at room temperature for 5 min. Next, 2.5 μl VigoFect was added to the diluent (total 100 μl), and the solution was mixed gently and placed at room temperature for 5 min. Next, the diluted VigoFect was added into the diluted DNA solution gently, and the resulting transfection working solution was incubated at room temperature for 15 min. Then, working solution was added to the culture solution, mixed gently and incubated for 24 h. After 24 hours of transfection, the pyrophosphates **Ge1, Ge5, Ge6** (1 mM) were added into the medium for 4 h. Cells were washed with PBS 3 times and then fixed with 4% paraformaldehyde for 15 min, and subsequently permeabilized with 0.5% Triton X-100 for 15 min. Next, tTMSN_3_ (50 μM) and Selectfluor (50 μM) were added into **Ge1** group for 30 min, cells were then incubated with DBCO-Cy5 (5 μM) for 30 min and then stained with 4⍰, 6-diamidino-2-phenylindole (DAPI) for 15 min in darkness at room temperature (RT). Samples were washed with PBS three times and examined under a confocal microscope (*LSM780, Carl Zeiss*).

### Amplification, expression, and extraction of *pCMV-Myc-SelU-GFP, pCMV-Myc-SelU-5s-GFP* plasmid

pCMV-Myc-SelU-GFP or pCMV-Myc-SelU-5s-GFP transformation used 1 μl plasmid being added to competent DH5α cells at 4 °C. The resulting cell – plasmid mixture was then mixed gently and kept in an ice-water bath for 30 min, then heat shockedat 42 °C for 90s, placed on ice for 5 min. 400 μl of LB solution was added, and the cells were incubated at 37 °C, 220 r.pm./min for 45 min. Then the bacterial solution was spread onto the Kan-containing LB solid culture plate. The plate was placed in 37 °C incubator for 12-16 h until a single colony appeared. Next, in the ultra-clean table, a single colony from the plate was picked and inoculated into 5 mL of LB solution (kan), incubated overnight at 37 °C, 220 r.p.m./min, until the culture solution was turbid. Then, plasmid extraction was performed.

### Plasmid extraction adopts Dynake Biological Plasmid Extraction Kit

(1) 250 μl of buffer BL was added to the adsorption column AC and centrifuged at 12,000/g for 1 min to activate the silica gel membrane. (2) 4 mls of the overnight bacterial culture was collected, centrifuged at 12,000/g for 1 min, and the bacterial cells were collected. (3) 200 μl of buffer S1 was added to resuspend the bacterial pellet, the solution was vortexed and centrifuged. (4) 200 μl of buffer S2 was added, mixed for 7 times to fully lyse the bacteria and (5) 200 μl of buffer S3 was added, and was mixed for 7 times. Next, the solution with white flocculent precipitate was centrifuged at 12,000/g for 15 min. (6) The supernatant was aspirated and transferred into the adsorption column AC, centrifuged at 12000/g for 1 min, the waste liquid was discarded, and the adsorption column AC was put back into the empty collection tube. (7) 700 μl of buffer W2 was added into the adsorption column AC, centrifuged at 12,000/g for 1 min, and the waste liquid was discarded, with this step repeated once. (8) The adsorption column AC was put back into the empty collection tube and centrifuged at 12000/g for 2 min. (9) The adsorption column AC was transferred to a clean 1.5 mL centrifuge tube, kept at 25 °C for 2 min, then 30 μl of Eluent was added to the middle of the adsorption membrane. It was then kept at 25 °C for 2 min, centrifuged at 12,000/g for 2 mi, where the plasmid was obtained, and the concentration was determined.

### General procedures for probe synthesis

General procedure: To a stirred solution of **7a** (5 mg, 0.001 mmol) in dry DMF (0.2 mL) was added **7b** (1 mg, 0.001 mmol) under nitrogen atmosphere. This mixture was stirred at room temperature overnight. 72 hours later, PTAD-N_3_ (**7d**, 2 mg, 0.007 mmol) was slowly added (in anhydrous DMF). The mixture was stirred at room temperature overnight. Compound **7e** was obtained through HPLC purification. [M+H]^+^ = 1355.6458. Probe **7f** was generated from **7e** which was formed by in-situ oxidation with NBS in DMF for 5 min, and was used directly without further purification considering the property of PTAD derivative in aqueous solution.

### General procedure for capturing of geranylated tRNA with its interacted proteins

In general, **First**, tRNA geranylation with *SelU in vitro*. 500 *μ*g *E. coli* tRNA (Roche) was incubated with 20 *μ*l purified *SelU* protein (1 mM), 100 *μ*l geranyl pyrophosphate (**Ge1**, 1 mM) in 100 *μ*l buffer (10 mM Tricine-NaOH, pH 8.0, 0.2 mM DTT, and 100 mM MgCl_2_, 2% glycerol at 25 °C) for 24 h. Next, the geranylated tRNA was incubated with NBS-oxidated probe (i.e., **7f**) for 10 min under icy-water bath. Then the *E. coli* lysate was co-incubated with the mixture at 4 °C for 30 min, and was irradiated with UV light (365 nm) for 15 min. The mixture was incubated at 4 °C while kept rotating overnight. Subsequently, the washed beads were added into the mixture, the beads-protein mixture was incubated and rotated for 4 h. Then the unbounded protein was removed using a magnetic stand. The sample was analyzed by SDS-PAGE, and the indicated band was collected and further analyzed by proteomics.

### Proteomic analysis and regulatory enzymes illustration

#### Technique route

In this project, a series of cutting-edge technologies, such as high-performance liquid chromatography (HPLC) classification technology and mass-spectrometry-based proteomics technology, were combined to detect and analyze the protein modification of a sample. The technical route details is showed in Supplementary Material. Representative procedure shows below:

#### Trypsin hydrolysis

The film was cut into 1 mm^3^ rubber blocks using a surgical blade and packed into a clean EP tube. Next the rubber blocks were washed with water 2-3 times, and 200 *μ*l of decolorizing solution (35% acetonitrile and 50 mM NH_4_HCO_3_) was added, with slight vibration used to decolorize. Next 70% acetonitrile (200 *μ*l) was added for dehydration, followed by DTT (200 *μ*l, 25 mM, dithiothreitol). The mixture was placed at room temperature for 1 h, the supernatant was discarded and NH_4_HCO_3_ (200 *μ*l, 50 mM) and IAM (iodoacetamide) were added and it was left at room temperature in the dark for 10 min. Next the supernatant was discarded, and 200 *μ*l of 70% acetonitrile was added for dehydration, and finally trypsin was added at the mass ratio of 1: 50 (trypsin: protein), and enzymolysis occurred at 37 °C overnight. Then the trypsin was added in the mass ratio of 1: 100 (trypsin: protein), and the enzymatic hydrolysis was continued for 4 h.

#### Database search and analysis

The String database and proteins identified as mitochondrial proteins were classified into ‘matrix’, ‘mitochondrial inner membrane (MIM)’, ‘intermembrane space (IMS)’ and ‘mitochondrial outer membrane (MOM)’.

### Cytotoxicity assays

When the density of HEK293T cells reached 80%, 96 plates were plated. The cells were washed twice with PBS, trypsin was added to digest the cells for 1 min, then 2 ml of complete medium was added, and was pipetted evenly, and the plate ws centrifuged at 1000 r.p.m. for 3 min. The supernatant was removed, 1 mL of fresh medium was added to the culture, and the cells were counted to verify there were 9 × 10^4^ cells. Each well was inoculated in the 96-well plate, and 100 μL of DMEM medium was added to each well, and a calculated amount of cell suspension was added, followed by pipetting and mixing, and incubation. Blank and **Ge1-Ge6** groups were settled (N = 4, each). After 24 h, to the medium was added **Ge1, Ge5, and Ge6** (1.0 mM). After culturing for 6 h, 10 μL of CCK-8 was added into each well in the dark, the edge of the plate was gently tapped to mix the solution, and was placed in the incubator for 1 h, and the OD value of each well was detected with a microplate reader at a wavelength of 450 nm. The cell viability was calculated as cell viability % = the average OD value of the experimental group / the average OD value of the blank group × 100%.

#### Reporting Summary

Further information on research design is available in the Nature Research Reporting Summary linked to this Article.

## Data availability

The genome sequence data that support the findings of this study are available in NCBI GenBank database under accession nos. AYSJ00000000, VNHN00000000, WSEY00000000, WSFA00000000 and WSFB00000000. For the corresponding genomes, see below **Table E2**. All other data generated or analyzed in this study are available within the Article and its Supplementary Information and Source Data. Source data are provided with this paper.

## Acknowledgements

We are grateful to Professors Sydney M. Hecht, Guangmin Yao, Yan Li and Peng Chen for their helpful discussions and comments on this manuscript. We thank Professors Ping Yin and Hua Li for SelU crystallization trials. We also thank Dr. Zhiqiang Wu for the mass-spec analysis of nucleoside samples, and the Analytical and Testing Center of HUST and Tongji Medical College Public Technical Service Platform, and the chemical core facility at UAlbany for NMR, mass-spec and confocal microscopy imaging support. We thank the startup fund of HUST, Shenzhen Huazhong University of Science and Technology Research Institute (to R. W.), and US NSF (MCB-1715234 and CHE-1845486 to J.S.) for the supports.

## Author contributions

R.W. and J.S. designed the project. J.W.Z., Y.Y.L, H.L.Z., and J.J.Z. performed compound isolation and structural elucidation under the supervision of R.W., J.S. H.L.Z. performed the chemical synthesis of PTAD-based probes 1-3. H.L.Z, Y.Y.L, and J.W.Z. conducted bioassays. R.W., and J.S. conducted molecular docking assays and structure analysis. Y.Y.L, and J.W.Z. conducted NMR spectra measurements. H.L.Z, Y.Y.L, J.W.Z., Y.Y.Z., and T.J.B. conducted SelU expression and purification, and some bioinformatic studies. R.W., L.W., and J.S. wrote the manuscript, with input from all co-authors.

## Additional information

**Supplementary information** The online version contains supplementary material available at https://doi.org/.

**Correspondence and requests for materials** should be addressed to Rui Wang (Huazhong University of Science and Technology) or Sheng Jia (University at Albany SUNY).

**Peer review information** Nature Chemistry thanks anonymous, reviewer(s) for their contribution to the peer review of this work.

**Reprints and permissions information** is available at www.nature.com/reprints.

## Extended Data

**Fig. E1.**
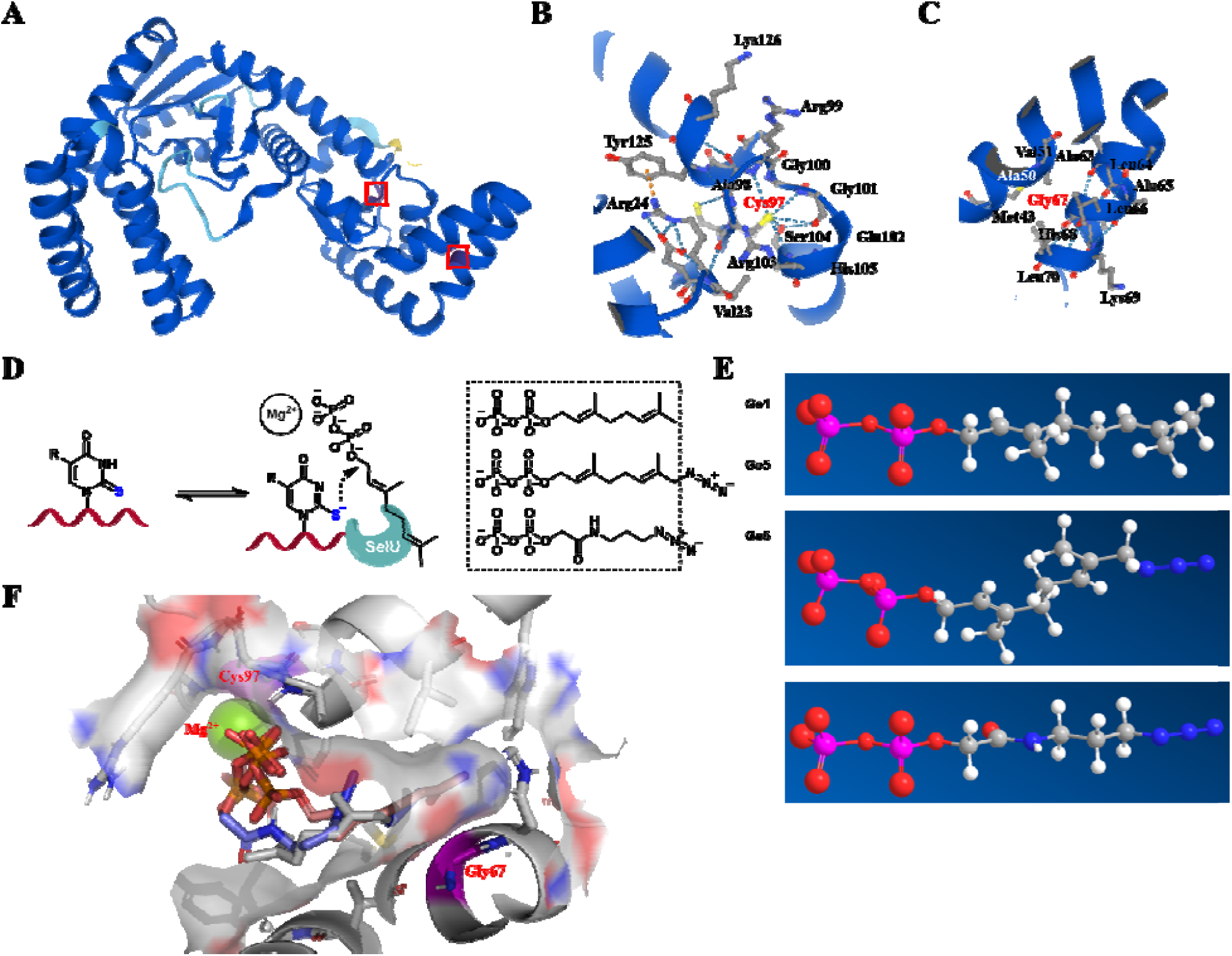
Structure prediction of the core domain in RNA geranylation enzyme *SelU*. (A) Ribbon diagram for the predicted structure of SelU based on the prediction from *AlphaFold*. (B) Predicated Cys97 interaction network. (C) Analysis of *SelU* critical sites Gly67. (D) Plausible mechanism of tRNA geranylation catalyzed by *SelU* enzyme in the presence of magnesium and prenyl pyrophosphate. The pyrophosphate structural comparisons between **Ge5, Ge6** and the natural substrate **Ge1** were also presented. (E) 3D structural comparisons of the pyrophosphate variants **Ge1, Ge5** and **Ge6**. (F) Molecule docking result from *Autodock Vina* demonstrated the structural similarities of SelU-ligand complexes with **Ge1, Ge5** and **Ge6**.

**Table E1.**
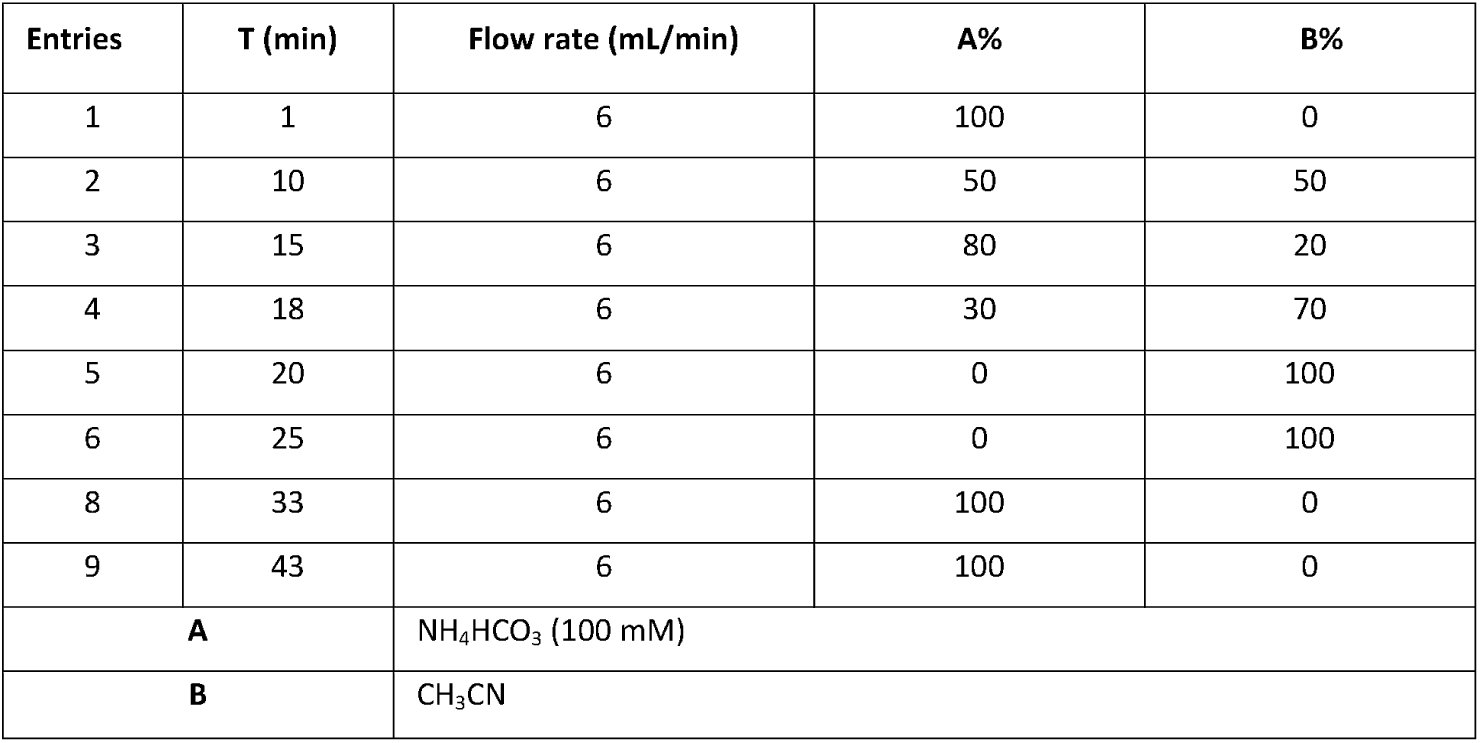
HPLC condition for isolation of pyrophosphates. Typical NMR data: ^31^**P NMR of Ge1** (162 MHz, D_2_O) δ - 6.85 (d, *J* = 22.68 Hz), −10.57 (d, *J* = 22.68 Hz).

| Entries | T (min) | Flow rate (mL/min) | A% | B% |
| --- | --- | --- | --- | --- |
| 1 | 1 | 6 | 100 | 0 |
| 2 | 10 | 6 | 50 | 50 |
| 3 | 15 | 6 | 80 | 20 |
| 4 | 18 | 6 | 30 | 70 |
| 5 | 20 | 6 | 0 | 100 |
| 6 | 25 | 6 | 0 | 100 |
| 8 | 33 | 6 | 100 | 0 |
| 9 | 43 | 6 | 100 | 0 |
| **A** | | $\text{NH}_4\text{HCO}_3$ (100 mM) | | |
| **B** | | $\text{CH}_3\text{CN}$ | | |

**Table E2.**
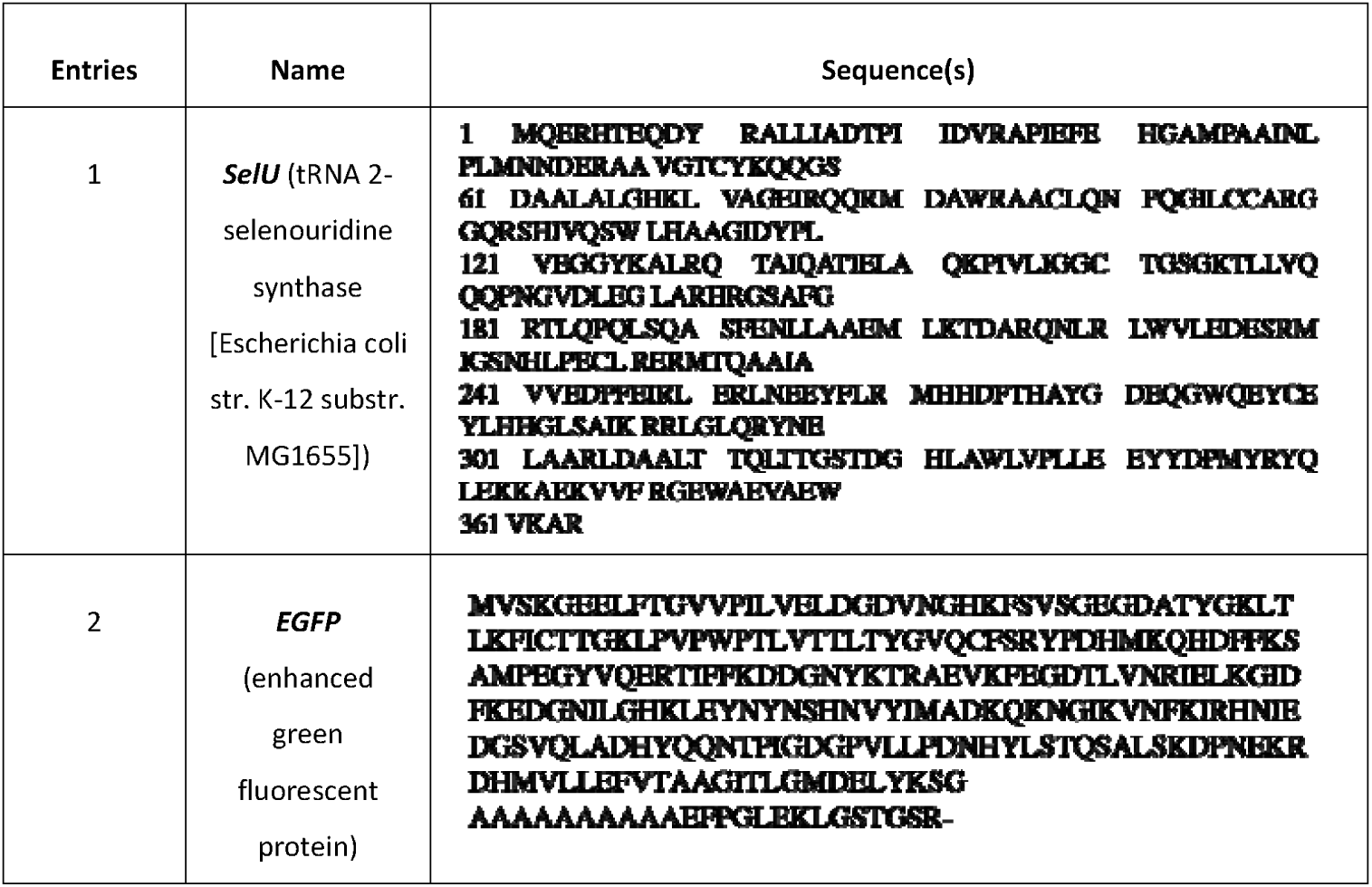
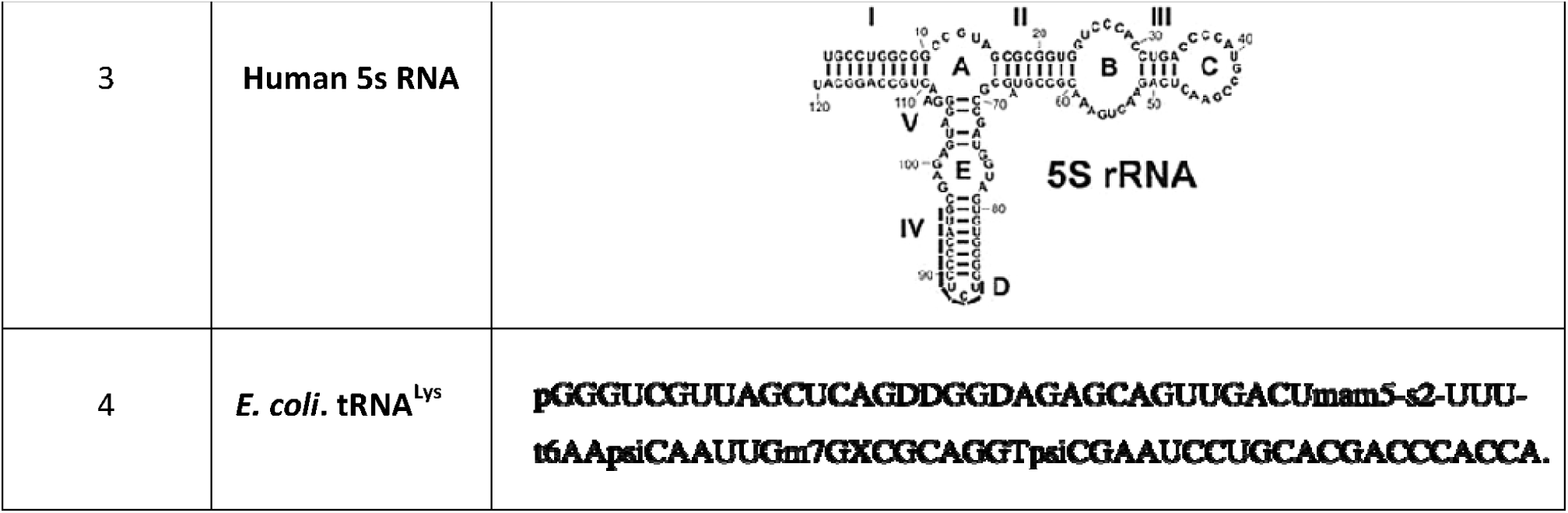
Summary of the gene sequences used in this work.

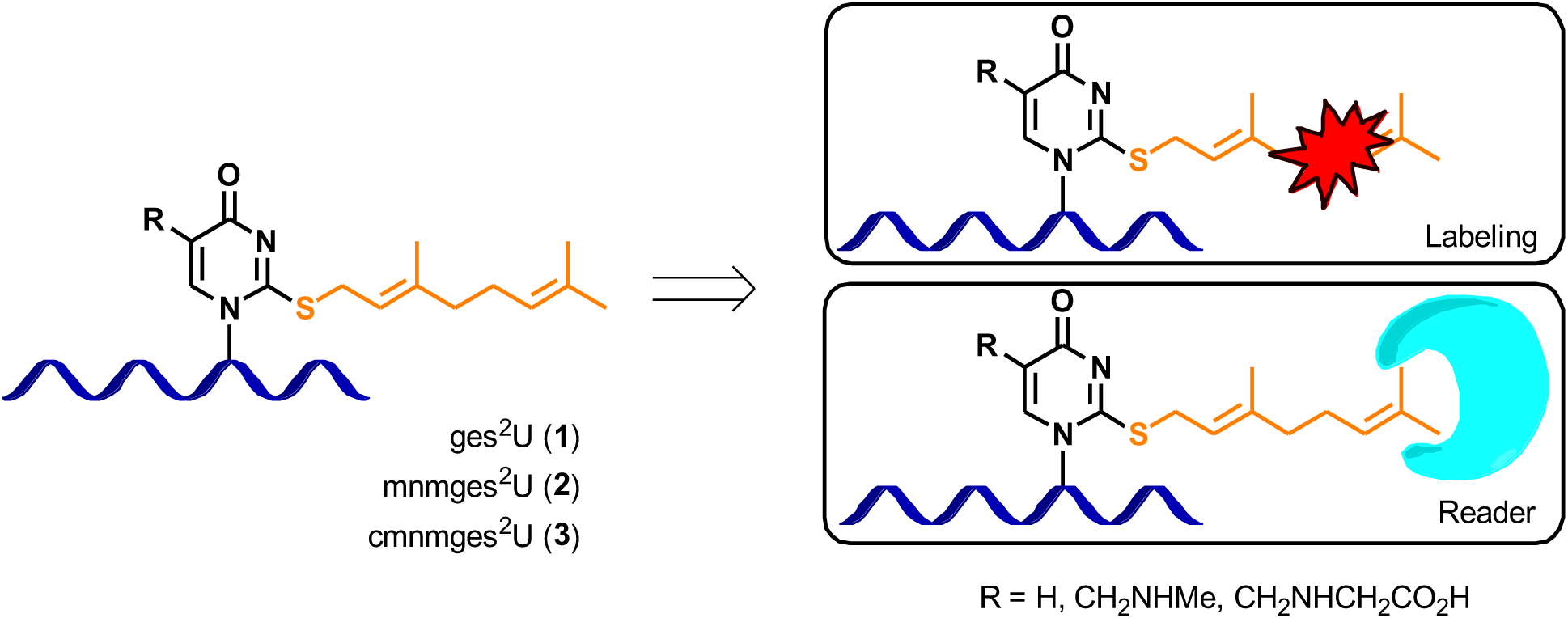

