## Supplemental Information for "Nature-Inspired Chemical Probes for in-cell Labeling of Lipidized RNA and Identification of New Regulatory Enzymes"

Table of Contents

Materials_••••••__•••••••••••••••••••••••••••••••••••••••••••••••••••••••••••••••••••••••••••••••••••••••••••••••••••••••••••••••••••••••••••••••••••••••••••••••••••••••_Page 2

Synthesis of geranyl variants (Ge1-7)_••••••••••••••__•••••••••••••••••••••••••••••••••••••••••••••••••••••••••••••••••••••••••••••••••••••••••••••••_Page 3

Synthesis of the standard nucleosides _•••••••••••••••••••••••••••••••••••••••••••••••••••••••••••••••••••••••••••••••••••••••••••••••••••••••••••_Page 29

Construction of *SelU* enzyme_•••••••••••••••••••••••••••••__•••••••••••••••••••••••••••••••••••••••••••••••••••••__••••••••••••••••••••••••••••••••••••••_Page 38

Structure elucidation of *SelU* and its molecular docking with pyrophosphate variants_•__••__••••••••••••••••••••••••_Page 46

*SelU*-mediated geranylation of tRNA in the fluorescent labeling investigations_••••••••••••••••••••••••••••••••••••••_Page 48

Direct labeling of prenylated tRNA using this Ene-ligation_••••••••••••••••••••••••••••••••••••••••••••••••••••••••••••••••••••••••_Page 62

Cell-specific labeling of *SelU*-mediated tRNA geranylation_••••••••••••••••••••••••••••••••••••••••••••••••••••••••••••••••••••••_Page 63

Installation of photo linking probe in tRNA Geranylation for studies of Reader proteins_•••••••••••••••••••••_Page 68

Proteomic and bioinformatic studies to identify geranylation regulatory proteins_••••••••••••••••••••••••••••••••••_Page 78

Reference_•••••••••••••••••••••••••••••••••••••••••••••••••••••••••••••••••••••••••••••••••••••••••••••••••••••••••••••••••••••••••••••••••••••••••••••••••••••••••_Page 84

**Materials**. Cas9 Nuclease, *Streptococcus pyogenes* (product# M0646), T7 Endonuclease I (product # M0302) Ribonucleotide solution mix (NTPs) and deoxy-ribonucleoside triphosphates (dNTPs) were purchased from *New England Biolabs* (USA). Transcript Aid T7 High Yield Transcription kit (product # K0441) and Glycogen (product # R0561) were purchased from *Thermo Fisher Scientific*. Pyrobest™ DNA Polymerase and PrimeSTAR HS DNA Polymerase were purchased from *TaKaRa Shuzo Co. Ltd.* (Tokyo, Japan). DNA Clean & Concentrator™-5 kit (product # D4014) was purchased from *Zymo Research Corp.* The DNeasy Blood & Tissue Kit was purchased from *QIAGEN*. The oligonucleotides at HPLC purity were obtained from *TaKaRa Company* (Dalian, China). The nucleic acid stains Super GelRed (No.: S-2001) was bought from *US Everbright Inc.* (Suzhou, China). DPBA (Cas # 17261-28-8), TCEP (Cas # 51805-45-9), 4-(2-hydroxyethyl)-piperazine-1-ethanesulfonic acid (HEPES, Cas # 7365-45-9) and Thiazolyl Blue Tetrazolium Bromide (MTT, Cas # 298-93-1) were purchased from *Sigma-Aldrich Inc.* (Shanghai, China). DPBS (Cas # 63995-75-5) was purchased from *TCI Development Co., Ltd* (Shanghai). The concentration of DNA or RNA was quantified by NanoDrop 2000c (*Thermo Scientific*, USA). Gel Imaging was performed using Pharos FX Molecular imager (*Bio-Rad*, USA).

**SYNTHEIS OF GERANYL PYROPHOSPHATEE VARIANTS**

**1.1** Synthesis of (*E*)-3,7-dimethylocta-2,6-dien-1-yldiphosphate tetrabutylammonium trimer (**Ge1**).

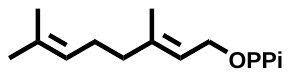

To a stirred solution of (*E*)-1-bromo-3,7-dimethylocta-2,6-diene (50.0 mg, 0.220 mmol) in CH_3_CN (1.5 mL) was slowly added tris(tetra-*N*-butylammonium) hydrogen pyrophosphate (200 mg, 0.264 mmol). The mixture was stirred for 5 h at r.t. under N_2_ atmosphere. The crude product was purified by HPLC (Table 1) and lyophilized to dryness.

**^1^H NMR** (400 MHz, D_2_O) δ 5.37 (t, *J* = 6.8 Hz, 1H), 5.11 (t, *J* = 6.7 Hz, 1H), 4.38 (t, *J* = 6.4 Hz, 2H), 2.06 (d, *J* = 7.4 Hz, 2H), 2.01 (d, *J* = 6.7 Hz, 2H), 1.60 (d, *J* = 4.4 Hz, 6H). **^31^P NMR** (162 MHz, D_2_O) δ -6.85 (d, *J* = 22.68 Hz), -10.57 (d, *J* = 22.68 Hz). **MS (ESI)**: Calculated [M+Na] ^+^ = 337.2, Found [M+Na] ^+^ = 337.0.

| **Entries** | T (min) | Flow rate (mL/min) | A% | B% |
| --- | --- | --- | --- | --- |
| **1** | 1 | 6 | 100 | 0 |
| **2** | 10 | 6 | 50 | 50 |
| **3** | 15 | 6 | 80 | 20 |
| **4** | 18 | 6 | 30 | 70 |
| **5** | 20 | 6 | 0 | 100 |
| **6** | 25 | 6 | 0 | 100 |
| **8** | 33 | 6 | 100 | 0 |
| **9** | 43 | 6 | 100 | 0 |
| **A** | | NH_4_HCO_3_ (100 mM) | | |
| **B** | | CH_3_CN | | |

**Table S1.** HPLC isolation condition.

1.2 Synthesis of 3,7-dimethyloct-6-en-1-yl pyrophosphate (**Ge2**).

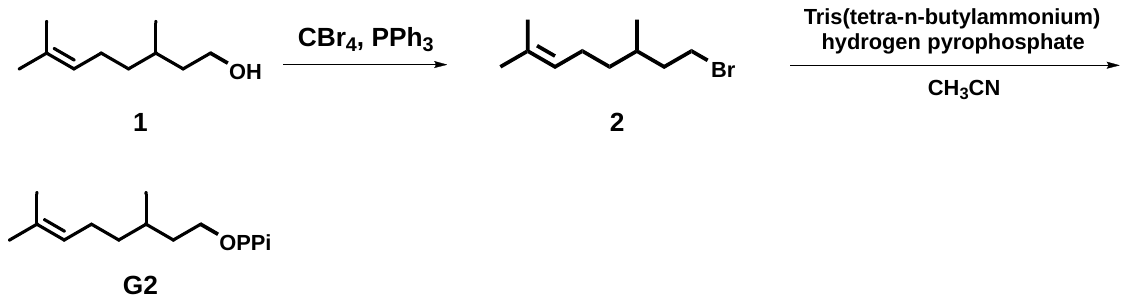

**Scheme S1**. Synthesis of pyrophosphate **Ge2.**

To a stirred solution of 3,7-dimethyloct-6-en-1-ol (150 mg, 1.05 mmol) was slowly added carbon tetrabromide (414 mg, 1.27 mmol) and triphenyl phosphorus (387 mg, 1.48 mmol) in CH_2_Cl_2_ (3.0 mL). The mixture was stirred at 0 ℃ for overnight under N_2_ atmosphere. The crude product was purified by silica gel flash chromatography to give 8-bromo-2,6-dimethyloct-2-ene (150 mg, 0.684 mmol, yellow oil). Then a solution of tris(tetra-n-butylammonium) hydrogen pyrophosphate (423 mg, 0.468 mmol) in CH_3_CN (1.5 mL) was added into 8-bromo-2,6-dimethyloct-2-ene (80.0 mg, 0.389 mmol). The mixture was stirred for 5 h at r.t. under N_2_ atmosphere. The crude product was purified by HPLC (Table 1) and lyophilized to dryness.

**^1^H NMR** (400 MHz, CDCl_3_) δ 5.13 - 5.02 (m, 1H), 4.10 (q, *J* = 7.1 Hz, 1H), 3.52 - 3.30 (m, 2H), 2.01 - 1.91 (m, 2H), 1.95 - 1.83 (m, 2H), 1.67 (s, 3H), 1.59 (s, 3H).

Compound **Ge2**: **^1^H NMR** (400 MHz, D_2_O) δ 5.21 (t, *J* = 7.2 Hz, 2H), 4.02 - 3.85 (m, 4H), 2.01 - 1.96 (m, 3H), 1.65 (s, 6H), 1.47 - 1.38 (m, 3H), 1.15 (dd, *J* = 14.7, 6.6 Hz, 2H), 1.06 (t, *J* = 7.5 Hz, 1H), 0.85 (m, 3H). **^31^P NMR** (162 MHz, D_2_O) δ - 8.45 (d, *J* = 17.3 Hz) - 10.53 (d, *J* = 21.06 Hz). **MS (ESI)**: Calculated [M+H] ^+^ = 336.2, Found [M+H] ^+^ = 336.3.

1.3 Synthesis of 4-methylpent-3-en-1-yl diphosphate tetrabutyl ammonium trimer (**Ge3**).

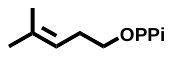

To a stirred solution of 5-bromo-2-methylpent-2-ene (50.0 mg, 0.310 mmol) was slowly added a solution of tris(tetra-n-butylammonium) hydrogen pyrophosphate (331 mg, 0.367 mmol) in CH_3_CN (1.5 mL). The mixture was stirred for 5 h at r.t. under N_2_ atmosphere. The crude product was purified by HPLC (Table 1) and lyophilized to dryness.

**^1^H NMR** (400 MHz, D_2_O) δ 5.17 (s, 1H), 3.81 (s, 2H), 2.27 (s, 2H), 1.84 (d, *J* = 101.2 Hz, 6H). **^31^P NMR** (162 MHz, D_2_O) δ -6.52 (d, *J* = 22.68 Hz), -10.59 (d, *J =* 21.06). **MS (ESI)**: Calculated [M+H] ^+^ = 536.4, Found [M+H] ^+^ = 536.8.

1.4 Synthesis of pyrophosphate **Ge4.**

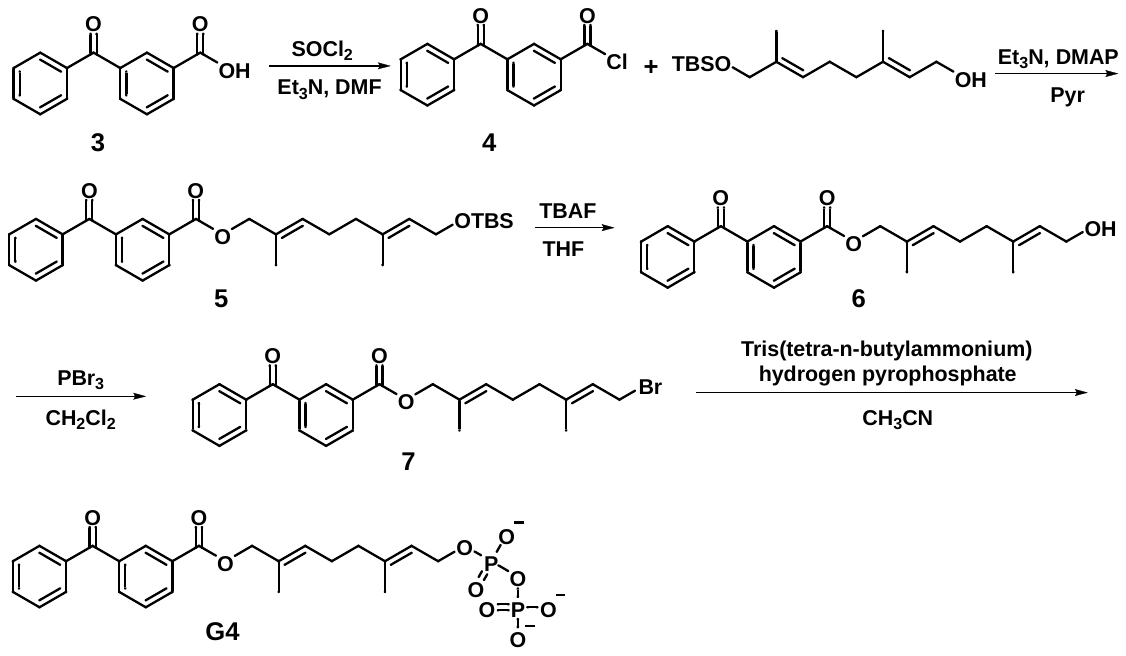

**Scheme S2**. Synthesis of pyrophosphate **Ge4.**

3-Benzoylbenzoic acid (450 mg, 2.10 mmol), SOCl_2_ (0.3 mL, 3.97 mmol) in CH_2_Cl_2_ (4.4 mL) and few drops of DMF was stirred for overnight at refluxed temperature. The reaction was washed with NaOH (0.1 M) and was extracted with dichloromethane and dried over Na_2_SO_4_. The solvents were removed and was used directly for the next step.

Approximately compound **4** (200 mg), (*2E,6E*)-8-((tert-butyldimethylsilyl) oxy)-3,7-dimethylocta-2, 6-dien-1-ol (125 mg, 0.400 mmol), Et_3_N (1.0 mL) and DMAP (4.0 mg) in pyridine (2.0 mL) was stirred for 5 h at 65 ℃. The reaction was quenched with water, washed with brine and extracted with CH_2_Cl_2_ and dried over Na_2_SO_4._ The crude product was purified by silica-gel flash chromatography. The rest of **3** was put in reaction at the same condition, quenched with water, washed with brine and extracted with CH_2_Cl_2_ and dried over Na_2_SO_4_. The crude product was purified by silica gel flash chromatography compound **5** (230 mg, 0.510 mmol), TBAF (300 *µ*L, 1.29 mmol) in THF (1.5 mL) was stirred overnight 30 ℃ under N_2_ atmosphere. The reaction was quenched with sat. NH_4_Cl. THF was removed and the residue was extracted with ethyl acetate and then dried over Na_2_SO_4_. The residue was purified by silica gel flash chromatography to give compound **6** (200 mg).

To a solution of **6** (100 mg, 0.260 mmol) in dichloromethane (2.0 mL) at 0 ℃, PBr_3_ (26.0 *µ*L, 0.270 mmol) was slowly added into the mixture. The reaction was stirred at r.t. overnight. Quenched with water washed with NaHCO_3_ and brine, extracted with dichloromethane dried over Na_2_SO_4_. The crude product was purified by silica gel flash chromatography to give compound **7** (300 mg, 0.068 mmol).

Compound **7** (300 mg, 0.068 mmol) in CH_3_CN (0.5 mL) was slowly added tris(tetra-n-butylammonium) hydrogen pyrophosphate (73.0 mg, 0.0820 mmol) in CH_3_CN (0.5 mL) at 0 ℃ under N_2_ atmosphere. The reaction was then stirred at r.t. for 3 h and was further purified by HPLC (Table 1).

Compound **5**: **^1^H NMR** (400 MHz, CDCl_3_) δ 8.44 (d, *J* = 7.2 Hz, 1H), 8.26 (t, *J* = 5.2 Hz, 1H), 8.02 - 7.96 (m, 1H), 7.80 (d, *J* = 8.9 Hz, 2H), 7.61 (t, *J* = 8.7 Hz, 1H), 7.56 (d, *J* = 7.7 Hz, 1H), 7.50 (t, *J* = 7.6 Hz, 2H), 5.47 (t, *J* = 7.0 Hz, 1H), 5.37 (t, *J* = 6.7 Hz, 1H), 4.86 (d, *J* = 7.1 Hz, 2H), 3.99 (s, 2H), 2.21 - 2.13 (m, 2H), 2.14 - 2.07 (m, 2H), 1.59 (s, 5H), 0.89 (s, 9H), 0.05 (s, 6H). **^13^C NMR** (101 MHz, CDCl_3_) δ 165.85 (s), 142.57 (s), 137.93 (s), 137.08 (s), 134.84 (s), 133.97 (s), 133.22 (s), 132.80 (s), 131.00 (s), 128.47 (s), 123.58 (s), 118.23 (s), 68.50 (s), 39.26 (s), 25.96 (s), 18.43 (s), 16.59 (s), 13.47 (s), 0.01 (s).

Compound **6**: **^1^H NMR** (400 MHz, CDCl_3_) δ 8.24 (d, *J* = 7.9 Hz, 2H), 7.96 (t, *J* = 6.5 Hz, 2H), 7.48 (t, *J* = 7.6 Hz, 5H), 5.50 (dd, *J* = 15.2, 8.3 Hz, 1H), 5.44 (d, *J* = 7.0 Hz, 1H), 4.83 (d, *J* = 7.1 Hz, 2H), 4.69 (s, 2H), 2.24 - 2.16 (m, 2H), 2.15 - 2.06 (m, 2H), 1.75 (s, 2H), 1.70 (s, 3H). **^13^C NMR** (101 MHz, CDCl_3_) δ 196.37 (s), 166.36 (s), 142.63 (s), 138.46 (s), 137.56 (s), 134.56 (d, *J* = 4.0 Hz), 133.73 (s), 133.35 (s), 131.52 (d, *J* = 5.1 Hz), 130.86 (s), 130.60 (s), 129.68 (s), 129.23 - 128.90 (m), 119.10 (s), 71.43 (s), 62.70 (s), 39.41 (s), 30.23 (s), 17.13 (s), 14.63 (s).

Compound **7**: **^1^H NMR** (400 MHz, CDCl_3_) δ 8.43 (d, *J* = 1.5 Hz, 2H), 8.25 (d, *J* = 7.7 Hz, 2H), 7.80 (d, *J* = 7.0 Hz, 5H), 5.22 (dd, *J* = 17.3, 1.0 Hz, 1H), 5.10 - 5.02 (m, 1H), 4.38 (d, *J* = 6.0 Hz, 2H), 3.95 (s, 2H), 2.15 - 2.08 (m, 2H), 2.07 - 2.02 (m, 2H), 1.74 (d, *J* = 5.5 Hz, 6H). **^13^C NMR** (101 MHz, CDCl_3_) δ 195.98 (s), 166.03 (s), 138.14 (s), 137.23 (s), 134.22 (s), 133.34 (s), 133.02 (s), 132.27 (s), 131.52 (s), 131.15 (s), 130.93 (s), 130.34 (d, *J* = 15.1 Hz), 129.69 (s), 128.72 (d, *J* = 9.6 Hz), 121.33 (s), 68.73 (s), 63.97 (s), 36.31 (s), 35.60 (s), 29.81 (s), 25.90 (s), 19.60 (s), 14.84 (s).

Pyrophosphate **Ge3**: **^1^H NMR** (400 MHz, D_2_O) δ 8.32 (d, *J* = 6.5 Hz, 4H), 8.11 (d, *J* = 8.1 Hz, 2H), 7.83 (d, *J* = 7.6 Hz, 5H), 7.76 (t, *J* = 8.1 Hz, 2H), 7.65 (t, *J* = 7.7 Hz, 4H), 5.61 (t, *J* = 5.8 Hz, 2H), 4.44 (s, 2H), 4.36 (d, *J* = 5.8 Hz, 3H), 2.13 - 2.07 (m, 2H), 1.89 - 1.82 (m, 2H), 1.72 (s, 6H). **^31^P NMR** (162 MHz, D_2_O) δ -9.71 (d, *J* = 26.3 Hz), -10.86 (d, *J* = 24.3 Hz). **MS (ESI)**: Calculated [M+H] ^+^ = 536.4, Found [M+H] ^+^ = 536.8.

1.5 Synthesis of pyrophosphate **Ge5.**

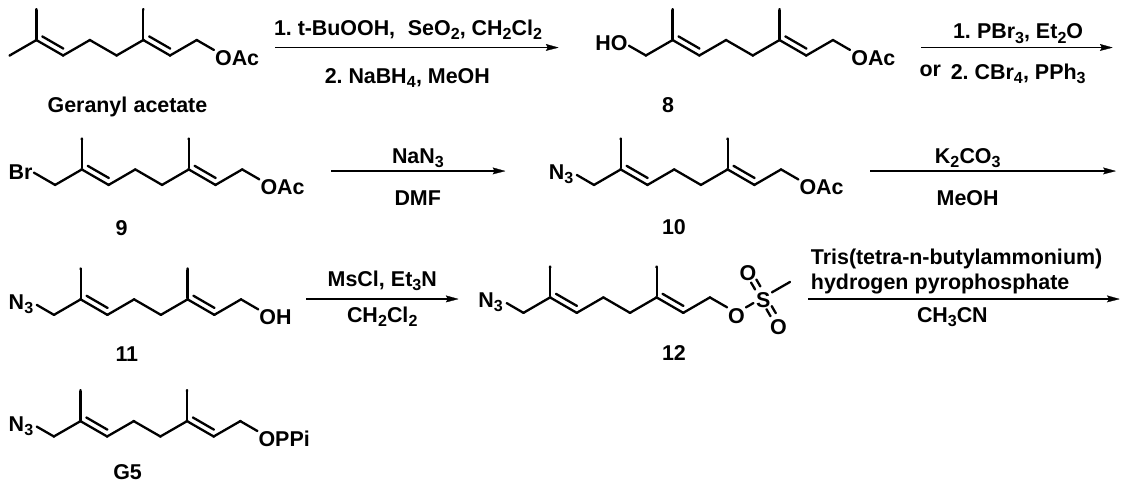

**Scheme S3**. Synthesis of pyrophosphate **Ge5.**

Pyrophosphate **Ge5**: (*E*)-3,7-dimethylocta-2,6-dien-1-ylacetate (201.0 mg, 1.03 mmol), *t*-BuOOH (122 *µ*L, 2.06 mmol), SeO_2_ (17.1 mg, 0.150 mmol) in CH_2_Cl_2_ (2.0 mL) stirred for 3 days at r.t. Cooled to 0 ^o^C, added NaBH_4_ (each time 100 mg, separately added for 6 times), was stirred for 5 h, till there was no foaming. Then the mixture was washed with H_2_O, extracted with CH_2_Cl_2,_ dried over Na_2_SO_4_. The crude product was purified by silica gel flash chromatography to yield compound (**8**, 120 mg, 0.566 mmol), PBr_3_ (1.00 mg, 4.85 mmol) in Et_2_O (0.8 mL). The solution was stirred for 4 h at ice-bathing under N_2_ atmosphere. The crude residue was purified by silica gel flash chromatography (crude).

Compound **9** (50.0 mg, 0.210 mmol), NaN_3_ (23.0 mg, 0.370 mmol) was dissolved in DMF (1.00 mL) stirred overnight refluxed at 75 ^o^C and purified by silica gel flash chromatography to offer **10** (50.0 mg, 0.211 mmol crude product).

To a stirred solution of compound **10** was added K_2_CO_3_ (58.0 mg, 0.421 mmol) in MeOH (2.0 mL) stirred for 3 h at r.t., washed with D.I.H_2_O and extracted with ethyl acetate and dried over Na_2_SO_4_. The mixtures were added into anhydrous CH_3_OH (5.0 mL) and was stirred overnight at r.t. under N_2_ atmosphere. The mixture was treated with NaHCO_3_ (sat.), dried over Na_2_SO_4_ (anhydrous), the crude product was purified by silica gel flash chromatography using eluent (10% ethyl acetate in PE) to give compound **11** (50.0 mg, 0.20 mmol crude product).

Compound **11** (50.0 mg, 0.200 mmol crude product) was dissolved in dichloromethane (1.0 mL), MsCl (20.0 *µ*L) and Et_3_N (40.0 *µ*L) was added to the mixture at 0°C. The reaction was stirred for overnight and reduced under *vacuo*, was further purified by silica gel flash chromatography which was used directly for next step.

To a stirred solution of compound **12** (50.0 mg, 0.183 mmol) was slowly added tris(tetra-*N*-butylammonium) hydrogen pyrophosphate (220 mg, 0.243 mmol) resolved in CH_3_CN (1.0 mL). The resulted mixture was stirred for 5 h at r.t. under N_2_ atmosphere. The crude product was purified by HPLC (Table 1) and lyophilized to dryness.

Alcohol **8**: **^1^H NMR** (400 MHz, CDCl_3_) δ 5.35 (m, 2H), 4.57 (s, 2H), 3.97 (s, 2H), 2.16 (dd, *J =* 14.6, 7.0 Hz, 2H), 2.07 (t, *J =* 5.9 Hz, 2H), 2.04 (s, 3H), 2.03 (s, 2H), 1.65 (s, 4H). **^13^C NMR** (101 MHz, CDCl_3_) δ 171.21 (s), 141.73 (s), 135.22 (s), 124.74 (s), 118.47 (s), 68.31 (s), 61.33 (s), 39.01 (s), 25.63 (s), 20.88 (s), 16.21 (s), 13.54 (s). **HRMS (ESI)**: Calculated [M+Na] ^+^ = 235.1310, Found [M+Na] ^+^ = 235.1308.

Bromide **9**: **^1^H NMR** (400 MHz, CDCl_3_) δ 5.56 (dd, *J* = 12.3, 6.2 Hz, 1H), 5.33 (t, *J* = 7.6 Hz, 1H), 4.57 (d, *J* = 7.1 Hz, 2H), 3.95 (s, 2H), 2.18 - 2.08 (m, 7H), 1.74 (s, 3H), 1.69 (s, 3H). **^13^C NMR** (101 MHz, CDCl_3_) δ 171.28 (s), 141.50 (s), 132.66 (s), 130.65 (s), 119.08 (s), 61.48 (s), 41.74 (s), 38.75 (s), 26.60 (s), 21.24 (s), 16.63 (s), 14.87 (s).

Azide **10**: **^1^H NMR** (400 MHz, CDCl_3_) δ 5.46 - 5.40 (m, 1H), 5.38 (m, 1H), 4.96 (dd, *J* = 2.6, 1.3 Hz, 2H), 4.56 (d, *J* = 7.1 Hz, 2H), 3.77 (dd, *J* = 7.5, 5.4 Hz, 2H), 2.32 - 2.20 (m, 7H), 1.70 (d, *J* = 0.8 Hz, 6H). **^13^C NMR** (101 MHz, CDCl_3_) δ 171.17 (s), 142.28 (s), 130.46 (s), 129.56 (s), 114.61 (d, *J* = 14.6 Hz), 68.80 (d, *J* = 6.7 Hz), 62.86 (s), 59.41 (s), 33.15 (s), 25.20 (s), 20.99 (s).

Compound **11**: **^1^H NMR** (400 MHz, CDCl_3_) δ 5.50 - 5.43 (m, 1H), 5.42 (t, *J* = 6.6 Hz, 2H), 4.98 (dd, *J* = 3.0, 1.5 Hz, 2H), 3.81 (d, *J* = 5.8 Hz, 2H), 2.05 (td, *J* = 14.8, 6.9 Hz, 4H), 1.72 (d, *J* = 0.8 Hz, 6H). **^13^C NMR** (101 MHz, CDCl_3_) δ 131.25 (s), 129.97 (s), 115.13 (d, *J* = 16.3 Hz), 69.35 (s), 61.61 (s), 59.99 (s), 40.33 (s), 37.28 (s), 33.87 (s), 30.23 (s), 29.62 (s), 25.81 (s), 19.96 (s), 18.04 (s), 15.14 (s).

Pyrophosphate **Ge5**: **^1^H NMR** (400 MHz, D_2_O) δ 5.52 - 5.48 (m, 2H), 3.60 - 3.56 (m, 2H), 2.17 (m, 5.3 Hz, 2H), 2.11 - 2.04 (m, 2H), 1.68 (s, 3H), 1.65 (s, 3H). **^31^P NMR** (162 MHz, D_2_O) δ -6.64 (d, *J* = 22.7 Hz), -10.44 (d, *J* = 22.7 Hz). **MS (ESI)**: Calculated [M+H] ^+^ = 375.2, Found [M+H] ^+^ = 375.5.

1.6 Synthesis of pyrophosphate **Ge6.**

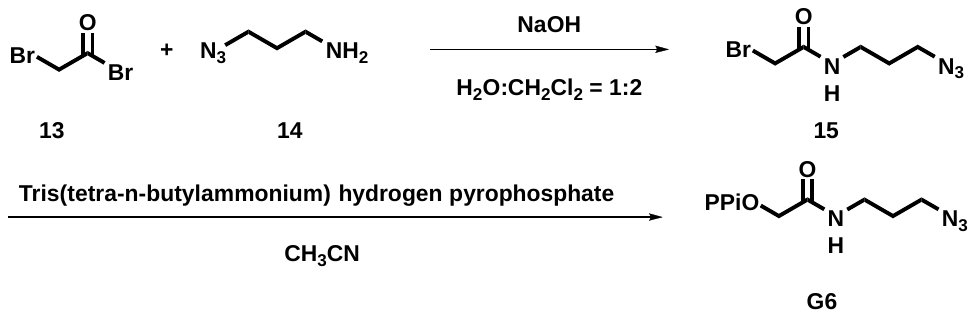

**Scheme S4**. Synthesis of pyrophosphate **Ge6.**

2-Bromoacetyl bromide **13** (0.696 mL, 7.99 mmol, 8 eq.), 3-azidopropan-1-amine **14** (100 mg, 0.999 mmol) and NaOH (200 mg, 5.00 mmol) in CH_2_Cl_2_/ H_2_O (2.5 mL-1.2 mL) was stirred for overnight at r.t. The aqueous layer was extracted with CH_2_Cl_2_, washed with Na_2_CO_3_ (50 mM), dried over Na_2_SO_4,_ and then was purified by silica gel flash chromatography to give compound **15.**

To a stirred solution of compound **15** (50.0 mg, 0.226 mmol) was slowly added tris(tetra-n-butylammonium) hydrogen pyrophosphate (220.1 mg, 0.243 mmol) resolved in CH_3_CN (1.0 mL). The resulted mixture was stirred for 5 h at r.t. under N_2_ atmosphere. The crude product was purified by HPLC (Table 1) and lyophilized to dryness.

Compound **15**: **^1^H NMR** (400 MHz, CDCl_3_) δ 4.22 (t, *J* = 6.0 Hz, 2H), 4.12 (t, *J* = 14.3, 7.3 Hz, 3H), 2.46 - 2.36 (m, 2H), 2.33 - 2.25 (m, 2H).

Pyrophosphate **Ge6**: **^1^H NMR** (400 MHz, D_2_O) δ 4.88 (d, *J* = 6.4 Hz, 2H), 3.50 - 3.29 (m, 2H), 1.17 (dd, *J* = 14.9, 7.4 Hz, 2H), 1.07 (t, *J* = 7.4 Hz, 2H). **^31^P NMR** (162 MHz, D_2_O) δ - 5.87 (d, *J* = 22.7 Hz), - 11.19 (d, *J* = 22.3 Hz). **MS (ESI)**: Calculated [M+H] ^+^ = 316.1, Found [M+H] ^+^ = 316.5.

1.7 Synthesis of pyrophosphate **Ge7**.

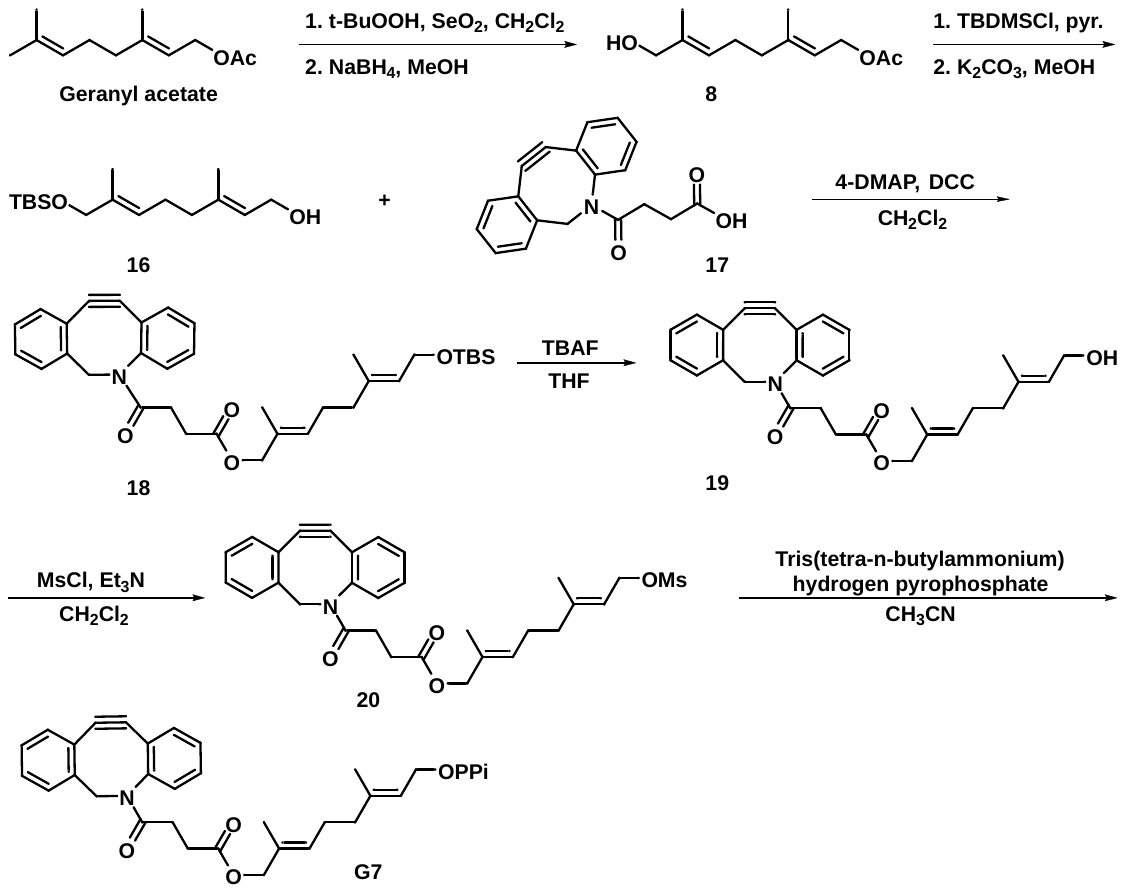

**Scheme S5**. Synthesis of pyrophosphate **Ge7.**

(*2E,6E*)-8-((tert-butyldimethylsilyl) oxy)-3,7-dimethylocta-2,6-dien-1-yl acetate (**16**). To a stirred solution of compound **8** (500 mg, 2.35 mmol) in aqueous pyridine (6.0 mL) was added *tert*-butyl chlorodimethylsilane (423 mg, 2.82 mmol). The resulted mixture was stirred for 2 h at 0 ℃ under N_2_ atmosphere. Upon competition of the reaction, the solvents were removed under *vacuo*, the concentrated product was taken up in CH_2_Cl_2_, then the crude product was purified by flash chromatography on silica gel column using petroleum ether-ethyl acetate (14: 1) as eluent to give compound **16** (600 mg) in 78% yield.

Add compound **16** (60.0 mg, 0.196 mmol), compound **17** (50.0 mg, 0.163 mmol), dicyclohexylcarbodiimide (80.0 mg, 0.326 mmol) and *N*-(4-pyridyl)-dimethylamine (10.0 mg, 0.150 mmol) into a stirred solution of dichloromethane (1.5 mL), then the mixture was stirred overnight at r.t. under N_2_ atmosphere. Upon competition of the reaction, HCl (0.1 M) was used to adjust pH = 5 - 6, and washed with saturated NaHCO_3_ and saturated aqueous NaCl, extracted with dichloromethane, dried over Na_2_SO_4._ The crude product was concentrated in rotary evaporator, was further purified by flash chromatography on silica gel column using petroleum ether-ethyl acetate (60: 1 to 30: 1) as eluent to give compound **18** (60.0 mg) in 64% yield.

To a stirred solution of compound **18** (60.0 mg, 0.105 mmol) in tetrahydrofuran (1.0 mL) was added tetrabutylammonium fluoride (40.0 *µ*L, 0.130 mmol). The resulted reaction mixture was stirred overnight at r.t. under N_2_ atmosphere. Upon competition of the reaction, the reaction was quenched with saturated NH_4_Cl and was further reduced in rotary evaporator. The crude product was extracted with CH_2_Cl_2_ and dried over Na_2_SO_4_. Then the crude product was concentrated in rotary evaporator, was further purified by silica gel flash chromatography using petroleum ether-ethyl acetate (30: 1 to 10: 1) as eluent to give alcohol **19** (320 mg) in 83% yield.

Alcohol **19** (50.0 mg, 0.109 mmol crude product) was dissolved in dichloromethane (1.0 mL), MsCl (20.0 *µ*L) and Et_3_N (40.0 *µ*L) was added to the mixture when cooled to 0 °C. The reaction was stirred overnight and reduced under *vacuum* to give compound **20** which was used directly for next step.

To a stirred solution of **20** was slowly added tris(*tetra*-*N*-butylammonium) hydrogen pyrophosphate (101 mg, 0.112 mmol) resolved in CH_3_CN (1.0 mL). The resulted mixture was stirred for 5 h at r.t. under N_2_ atmosphere. The crude product was purified by HPLC (Table 1) and lyophilized to dryness.

Compound **18**: **^1^H NMR** (400 MHz, CDCl_3_) δ 7.68 (d, *J* = 7.6 Hz, 1H), 7.51 (dd, *J* = 5.7, 3.2 Hz, 1H), 7.40 (dd, *J* = 8.9, 5.1 Hz, 2H), 7.37 (d, *J* = 1.4 Hz, 1H), 7.35 (d, *J* = 1.6 Hz, 1H), 7.35 - 7.32 (m, 1H), 5.34 (t, *J* = 6.8 Hz, 1H), 5.26 (t, *J* = 7.6 Hz, 1H), 5.16 (d, *J* = 13.9 Hz, 1H), 3.99 (s, 2H), 3.66 (d, *J* = 8.3 Hz, 1H), 2.74 (ddd, *J* = 16.2, 8.3, 6.4 Hz, 1H), 2.63 (ddd, *J* = 17.1, 8.3, 6.3 Hz, 1H), 2.31 (dt, *J* = 17.2, 6.1 Hz, 1H), 2.15 - 2.09 (m, 2H), 1.95 (dd, *J* = 10.2, 6.2 Hz, 2H), 1.92 (d, *J* = 6.0 Hz, 2H), 1.73 - 1.69 (m, 2H), 1.67 (d, *J* = 4.8 Hz, 2H), 1.65 (s, 3H), 0.89 (s, 8H), 0.04 (s, 6H).

Alcohol **19**: **^1^H NMR** (400 MHz, CD_3_OD) δ 7.54 (d, *J* = 8.1 Hz, 1H), 7.52 - 7.47 (m, 1H), 7.36 (d, *J* = 2.2 Hz, 2H), 7.29 - 7.19 (m, 2H), 7.15 (d, *J* = 7.2 Hz, 1H), 7.12 (d, *J* = 2.4 Hz, 1H), 5.25 (dd, *J* = 10.0, 7.1 Hz, 2H), 3.80 (s, 2H), 3.66 - 3.56 (m, 1H), 2.41 - 2.17 (m, 4H), 2.00 - 1.88 (m, 4H), 1.86 (s, 3H), 1.57 (s, 3H), 1.54 (s, 3H).

Pyrophosphate **Ge7**: **^1^H NMR** (400 MHz, CDCl_3_) δ 7.69 (dd, *J* = 17.0, 8.2 Hz, 4H), 7.61 - 7.56 (m, 4H), 7.37 (s, 2H), 4.30 (d, *J* = 1.5 Hz, 6H), 4.10 - 4.08 (m, 2H), 3.75 (s, 2H), 3.62 (s, 2H), 2.20 - 2.08 (m, 4H), 2.04 (s, 3H), 1.97 (s, 1H), 1.79 (s, 3H), 1.76 (s, 3H). **^31^P NMR** (162 MHz, D_2_O) δ -6.61 (d, *J* = 22.0 Hz), -10.74 (d, *J* = 19.5 Hz). **MS (ESI)**: Calculated [M-H] ^+^ = 613.5, Found [M-H] ^+^ = 613.0.

**NMR Spectra I of Pyrophospahte Analogues**

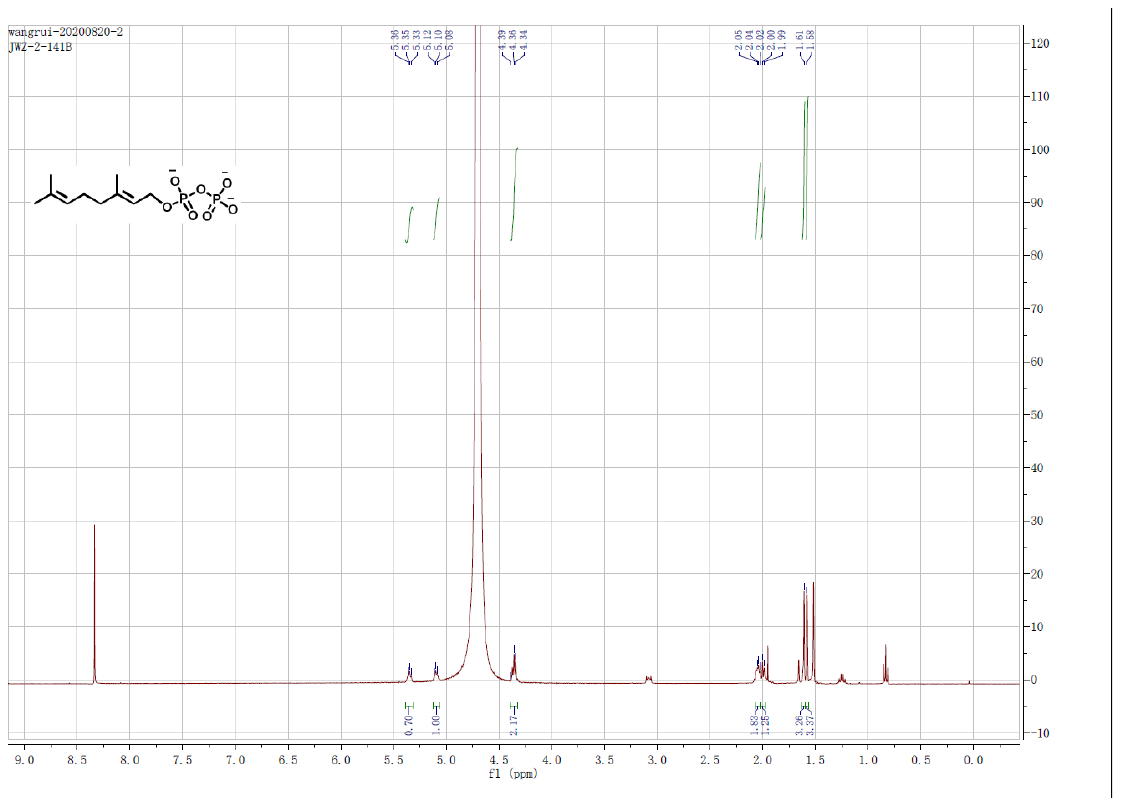

**Figure S1**. **^1^H NMR** spectra of pyrophosphate **Ge1**.

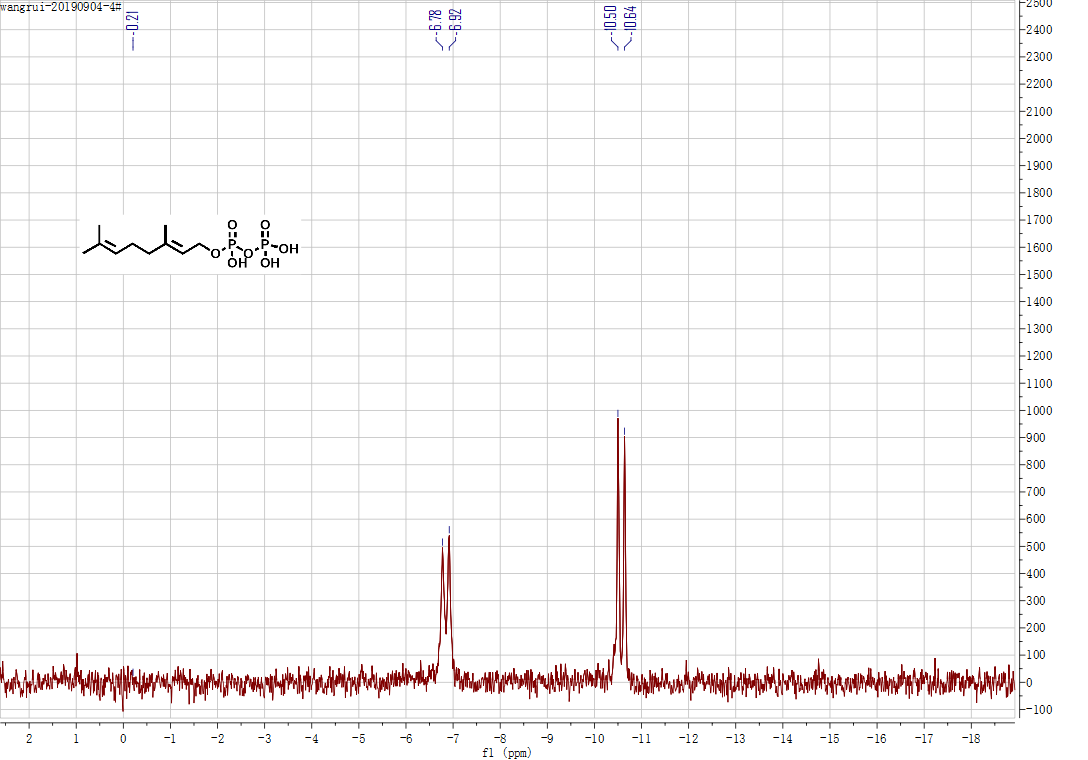

**Figure S2**. **^31^P NMR** spectra of pyrophosphate **Ge1.**

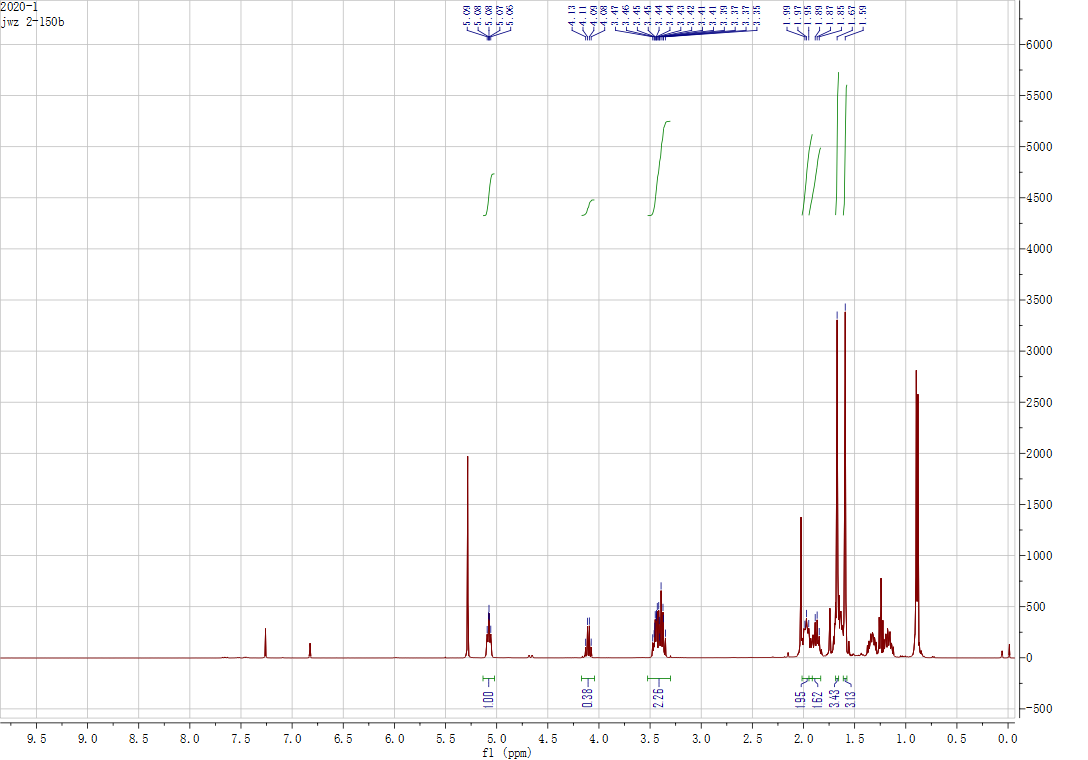

**Figure S3. ^1^H NMR** spectra of bromide **2.**

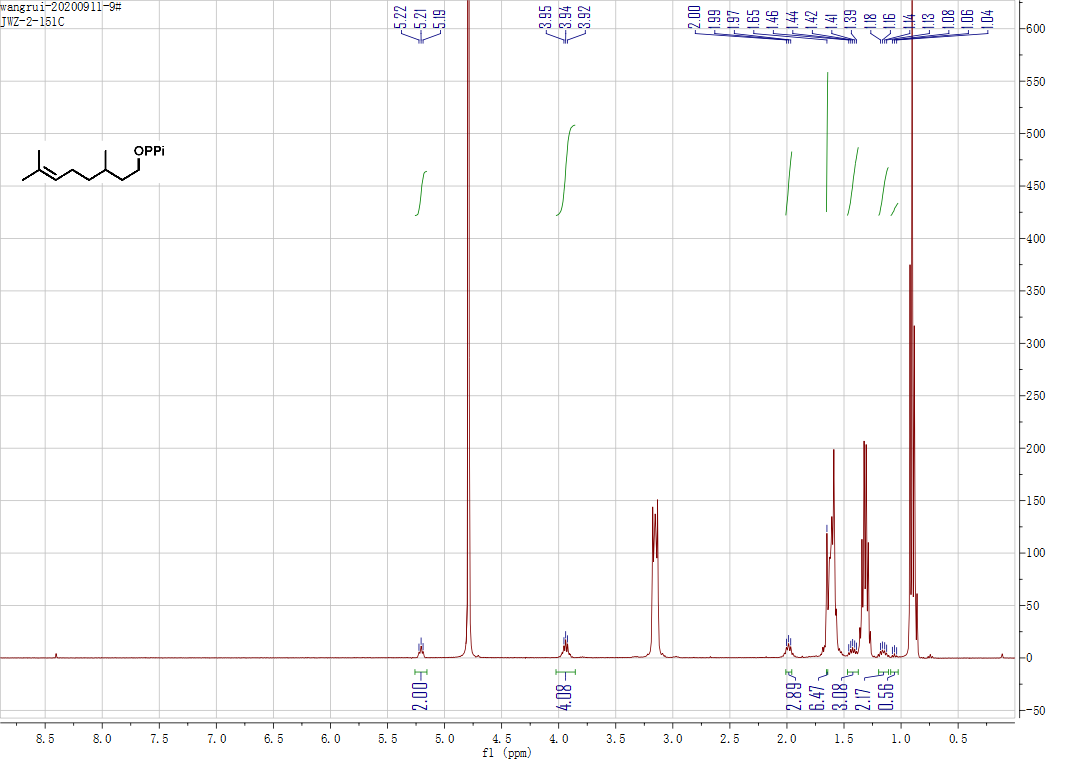

**Figure S4**. **^1^H NMR** spectra of pyrophosphate **Ge2**.

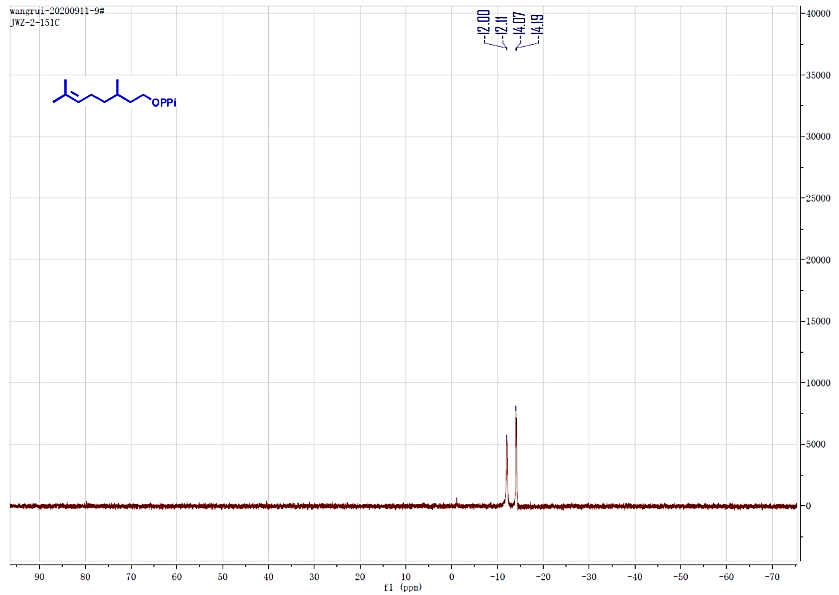

**Figure S5**. **^31^P NMR** spectra of pyrophosphate **Ge2**.

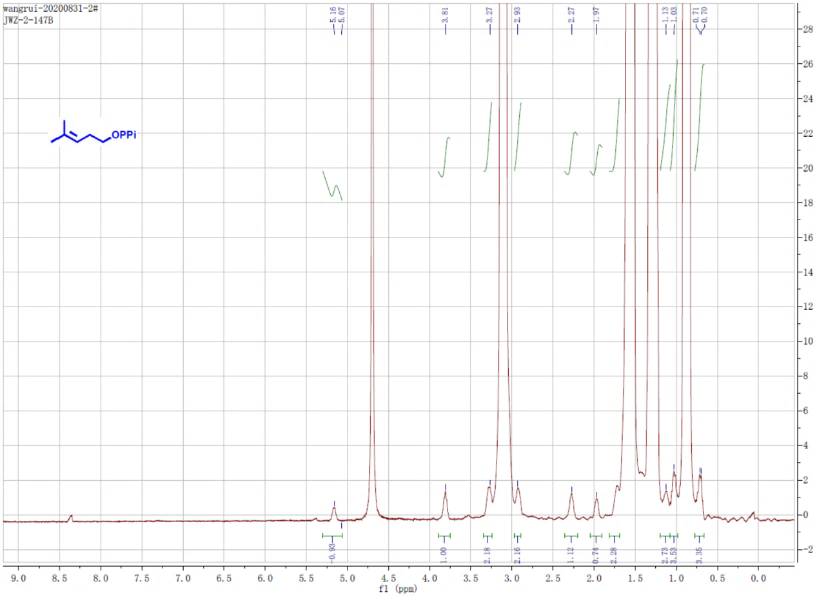

**Figure S6**. **^1^H NMR** spectra of pyrophosphate **Ge3**.

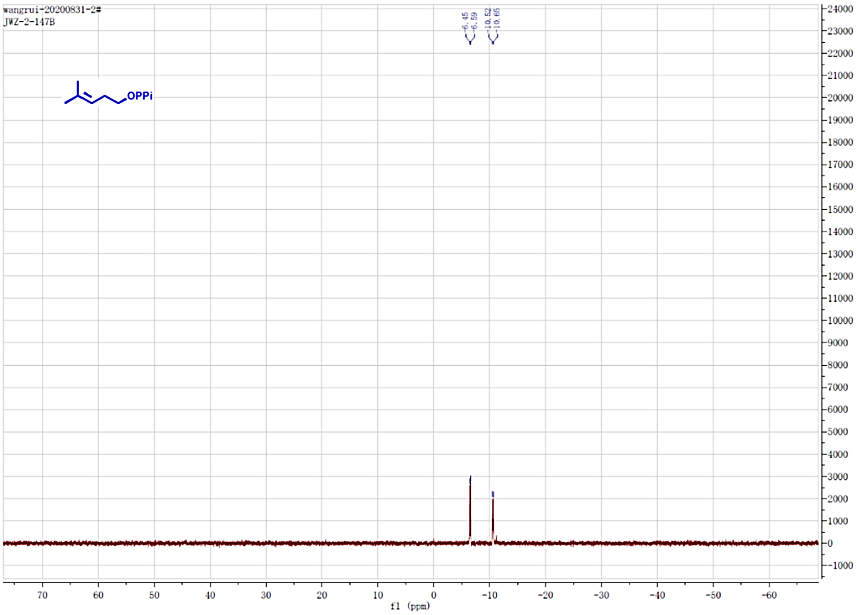

**Figure S7**. **^31^P NMR** spectra of pyrophosphate **Ge3.**

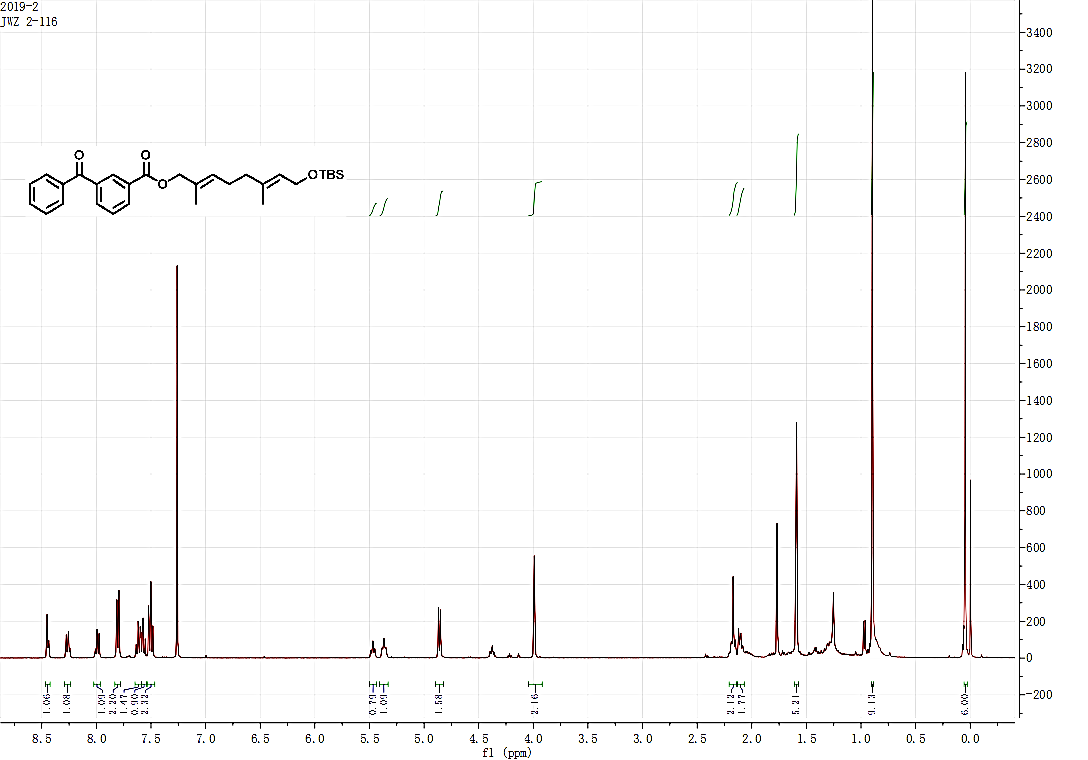

**Figure S8**. **^1^H NMR** spectra of compound **5**.

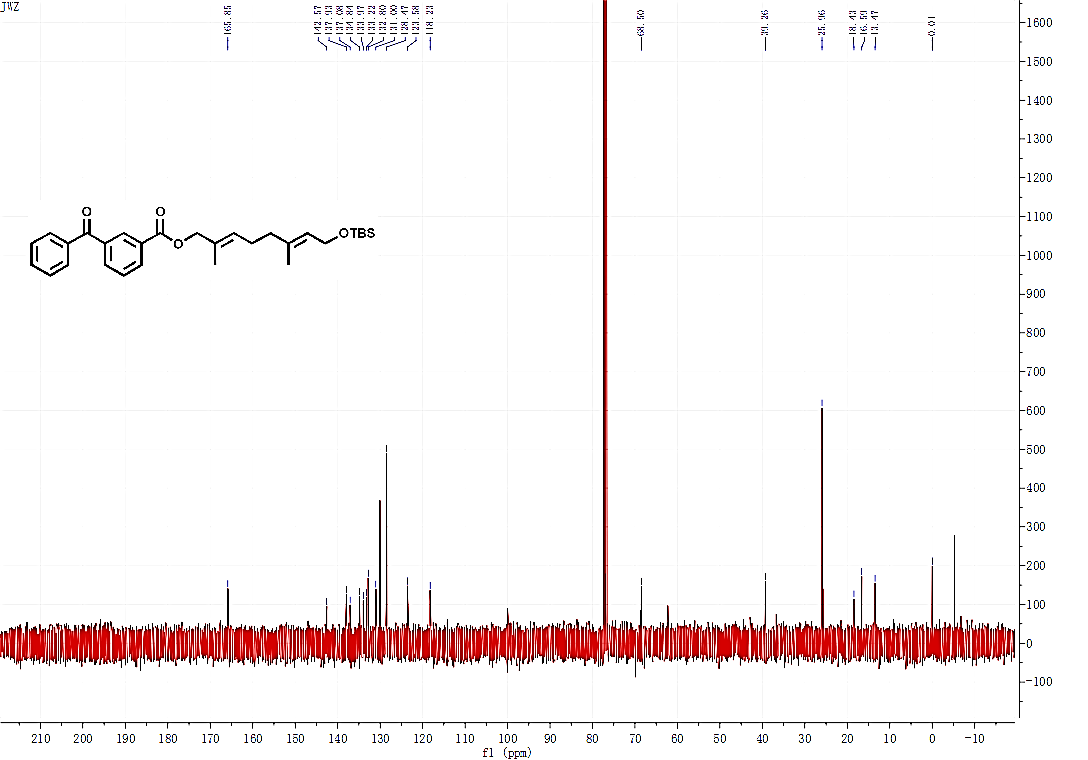

**Figure S9**. **^13^C NMR** spectra of compound **5**.

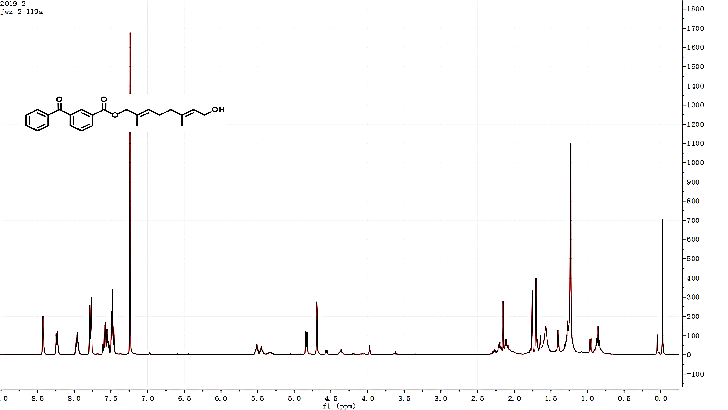

**Figure S10**. **^1^H NMR** spectra of alcohol **6**.

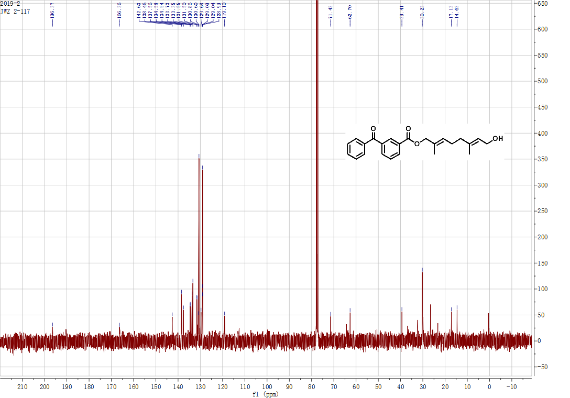

**Figure S11**. **^13^C NMR** spectra of alcohol **6**.

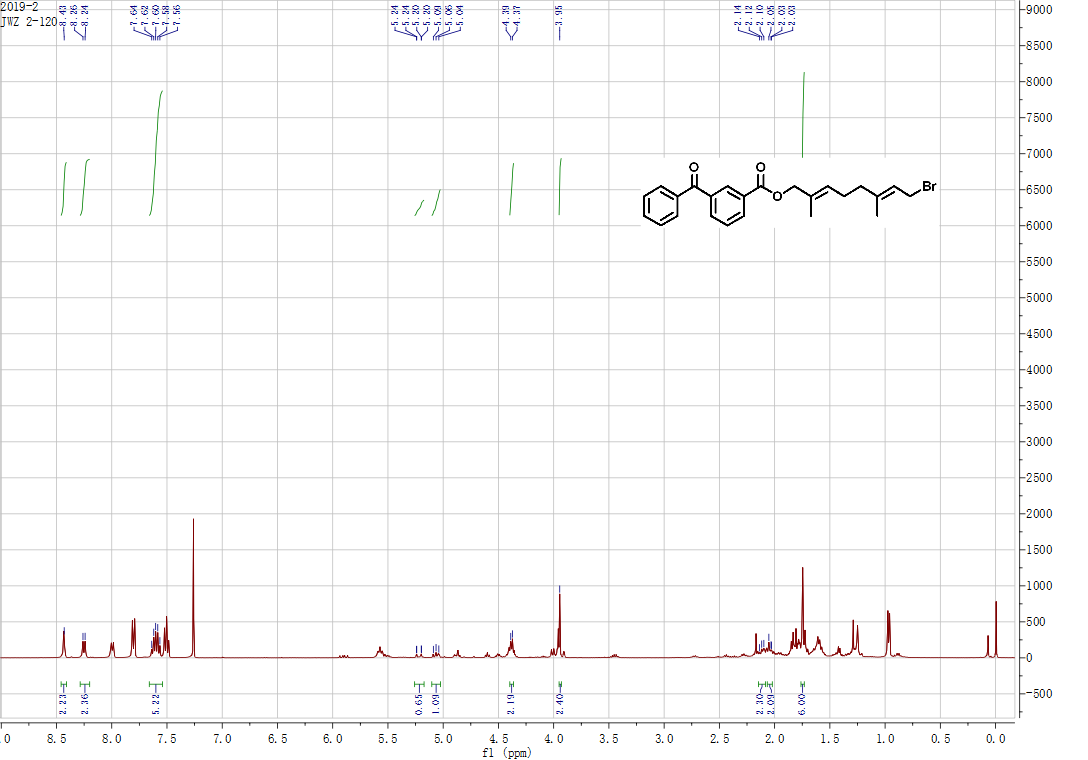

**Figure S12**. **^1^H NMR** spectra of bromide **7**.

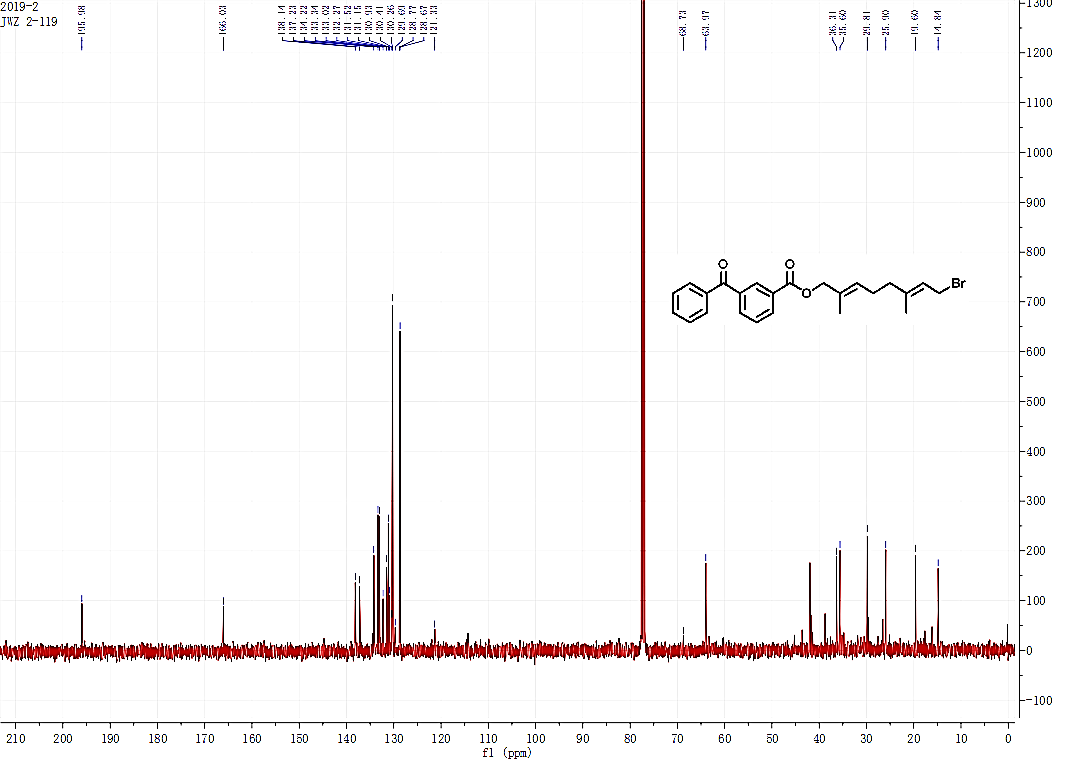

**Figure S13**. **^13^C NMR** spectra of bromide **7**.

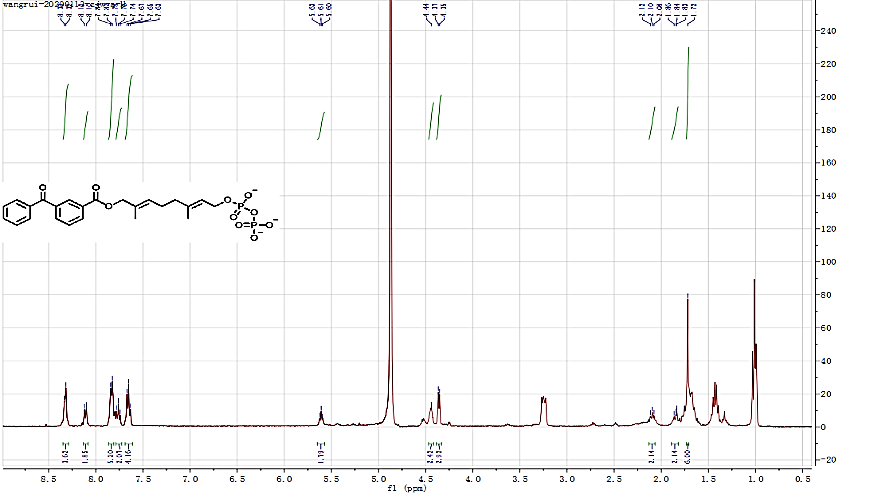

**Figure S14**. **^1^H NMR** spectra of pyrophosphate **Ge4**.

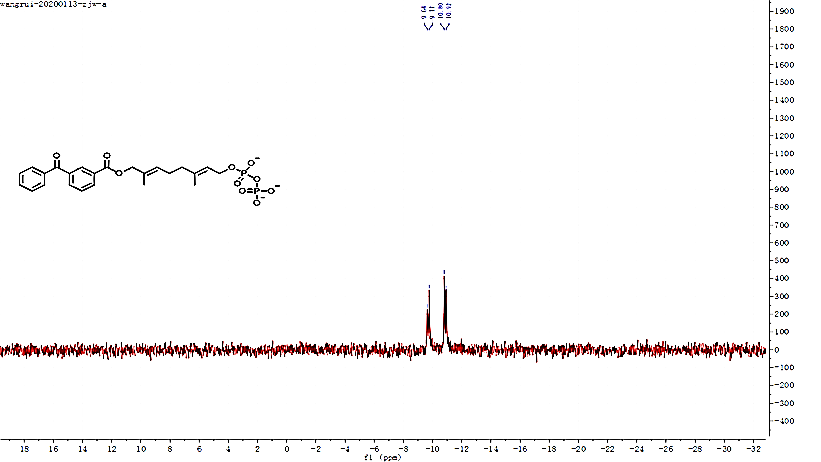

**Figure S15**. **^31^P NMR** spectra of pyrophosphate **Ge4**.

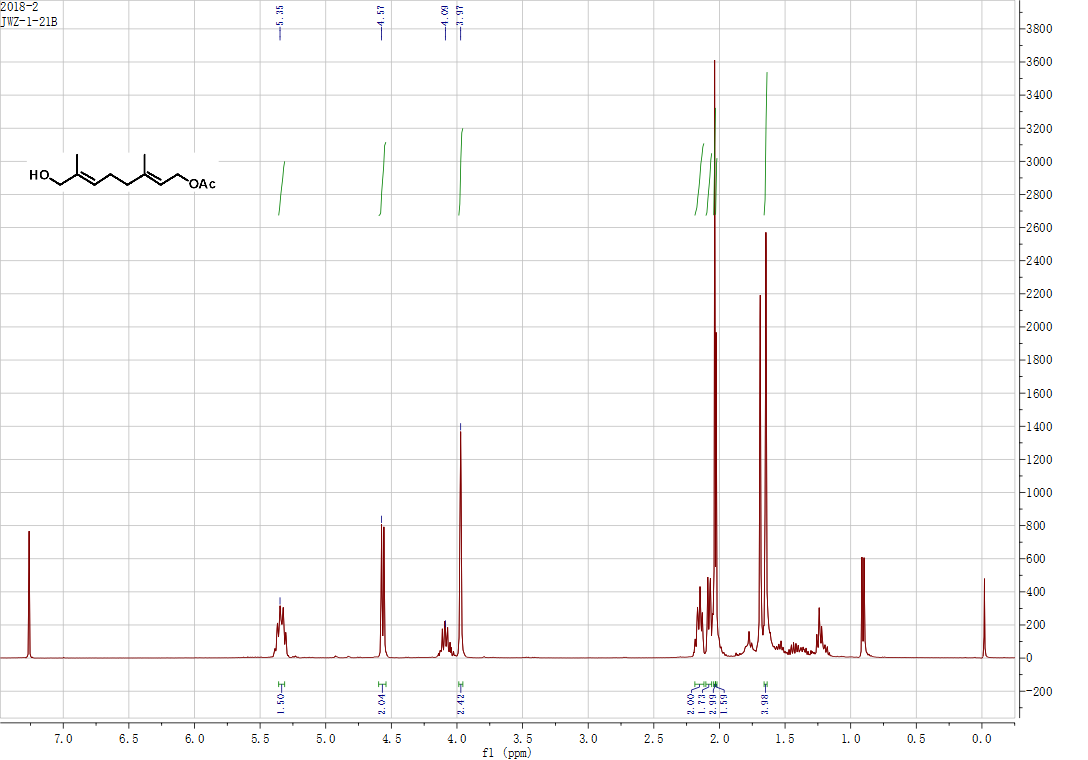

**Figure S16**. **^1^H NMR** spectra of alcohol **8**.

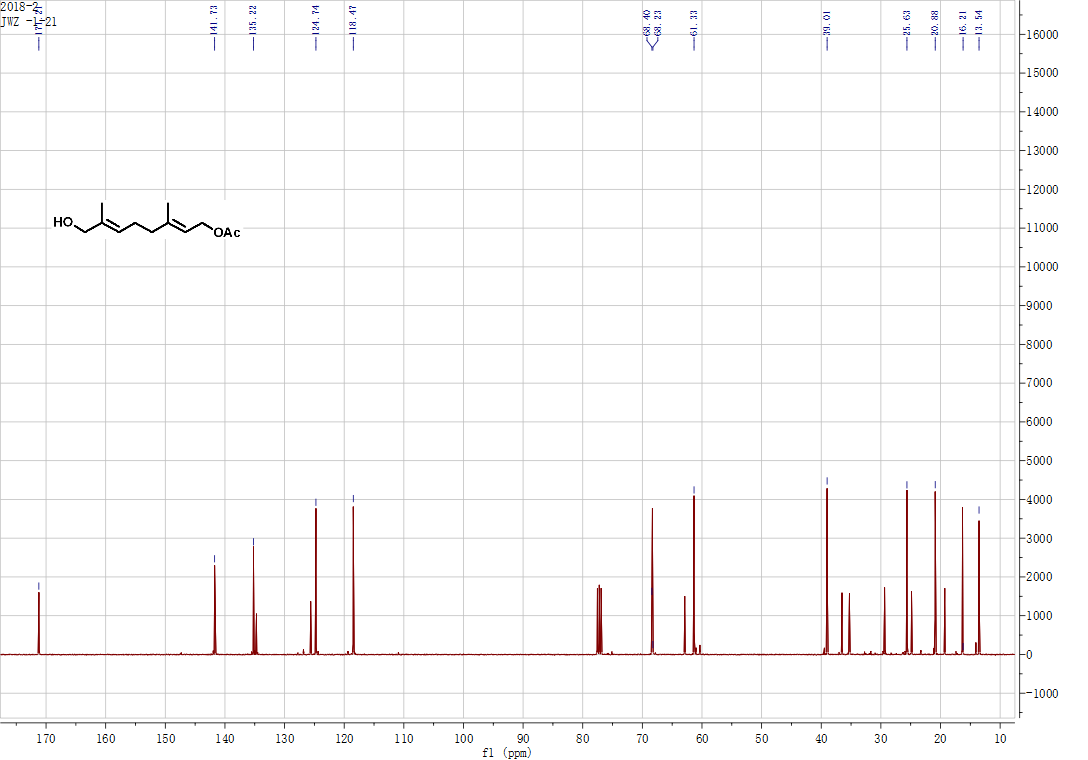

**Figure S17**. **^13^C NMR** spectra of alcohol **8.**

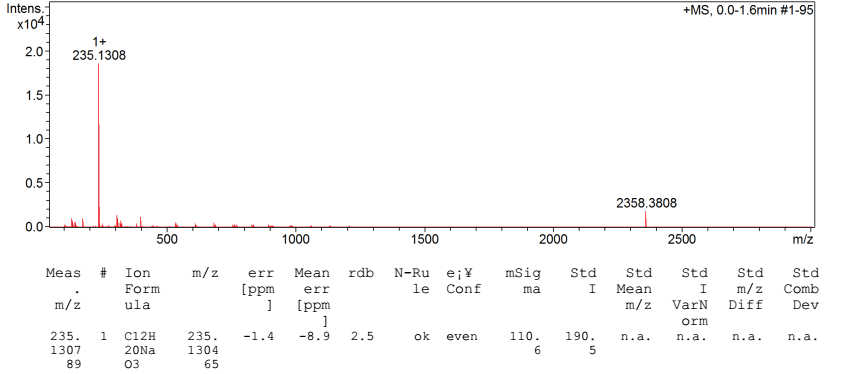

**Figure S18**. High resolution mass spectra of alcohol **8**.

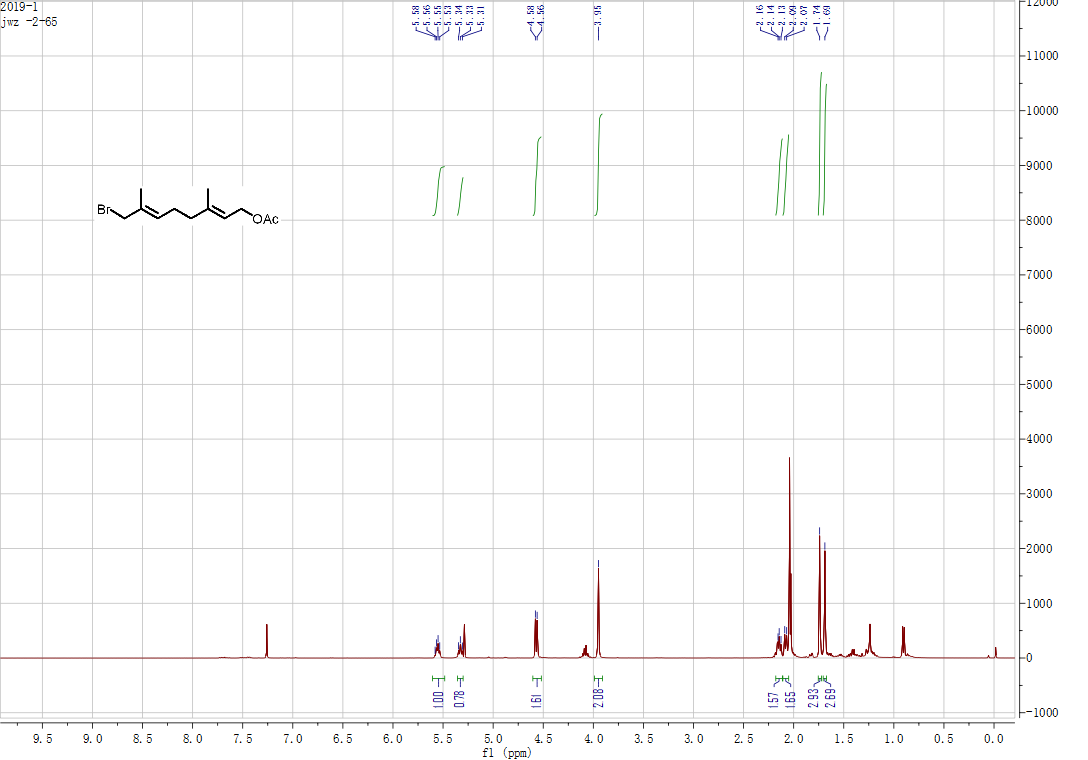

**Figure S19**. **^1^H NMR** spectra of bromide **9**.

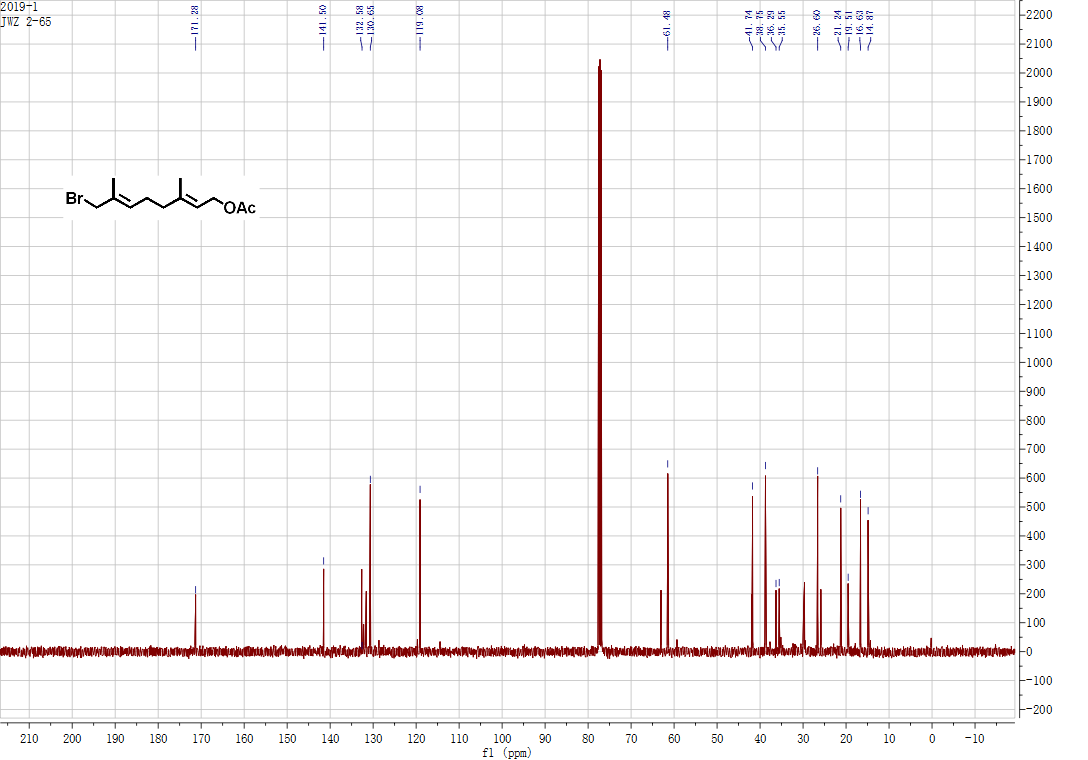

**Figure S20**. **^13^C NMR** spectra of bromide **9.**

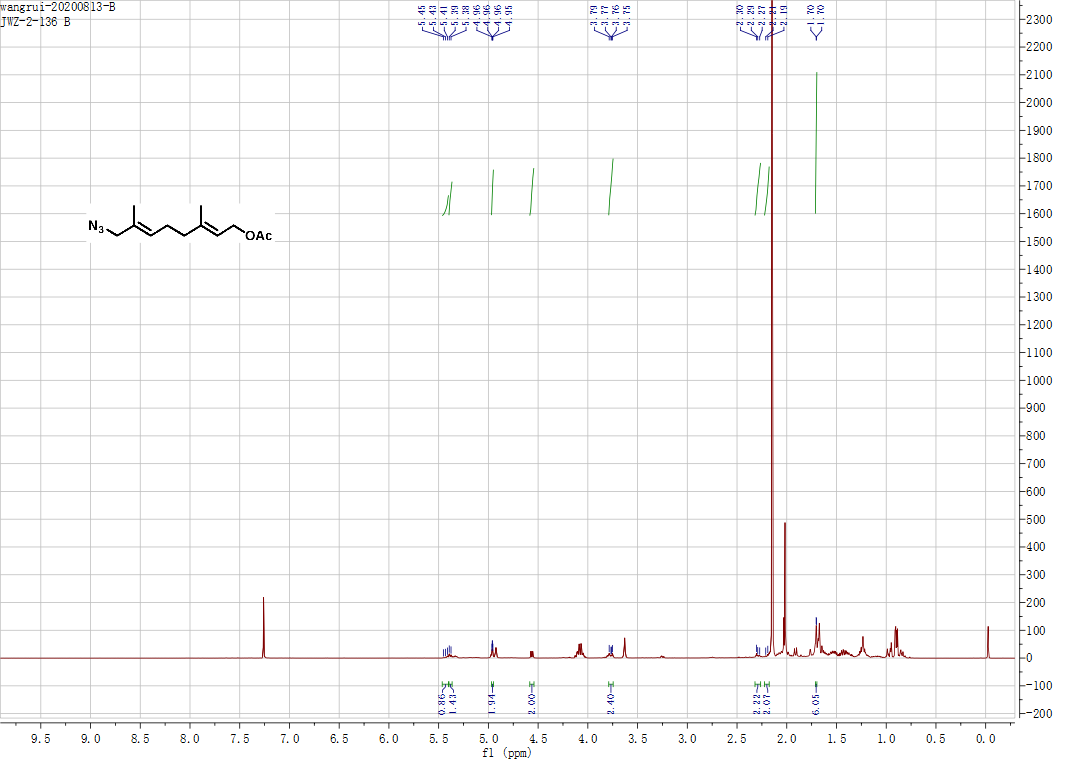

**Figure S21**. ^1^H NMR spectra of azide **10**.

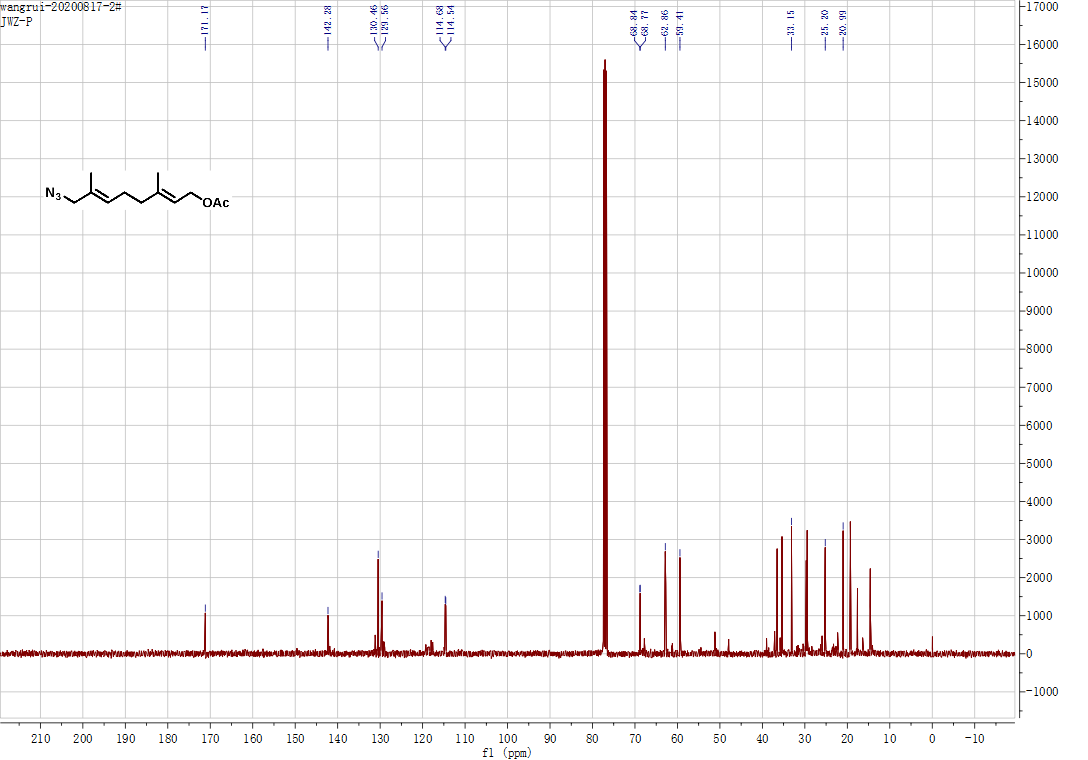

**Figure S22**. **^13^C NMR** spectra of azide **10.**

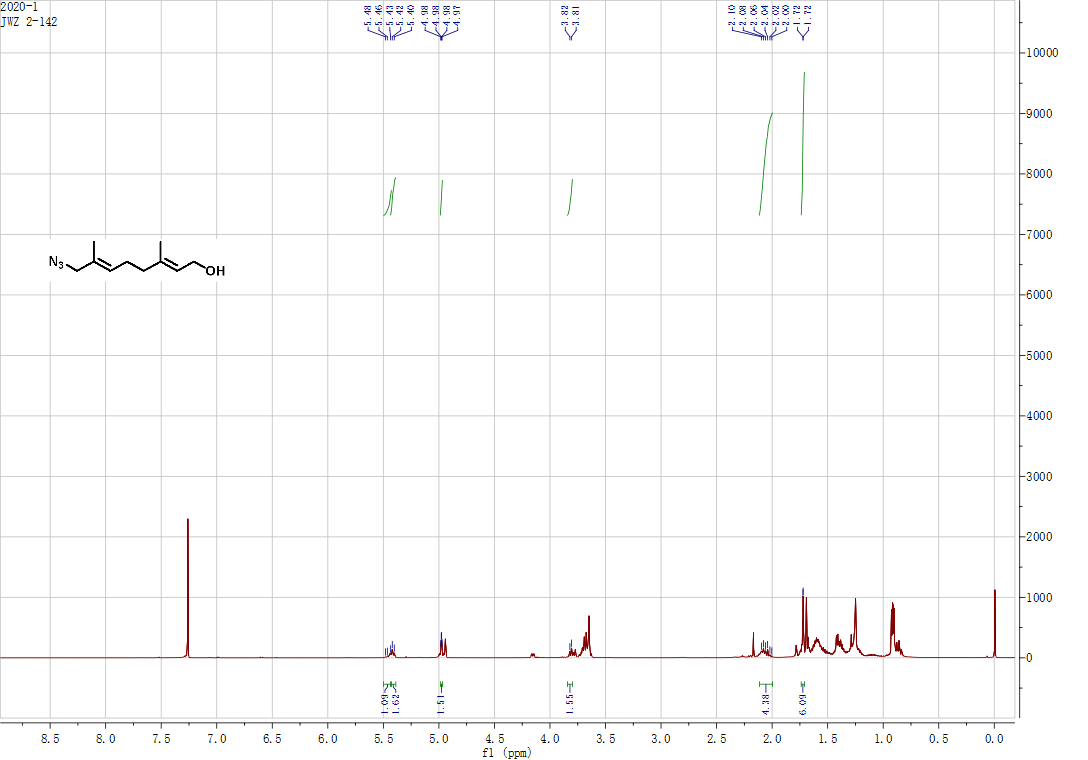

**Figure S23**. ^1^H NMR spectra of azide **11**.

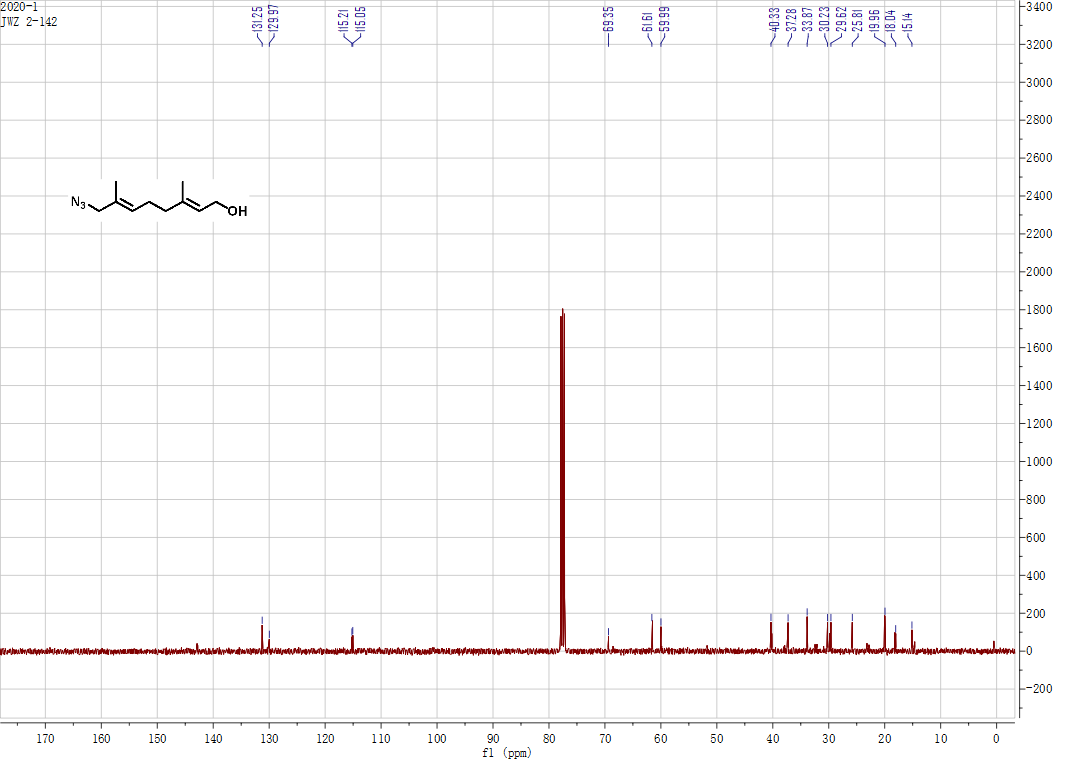

**Figure S24**. **^13^C NMR** spectra of azide **11.**

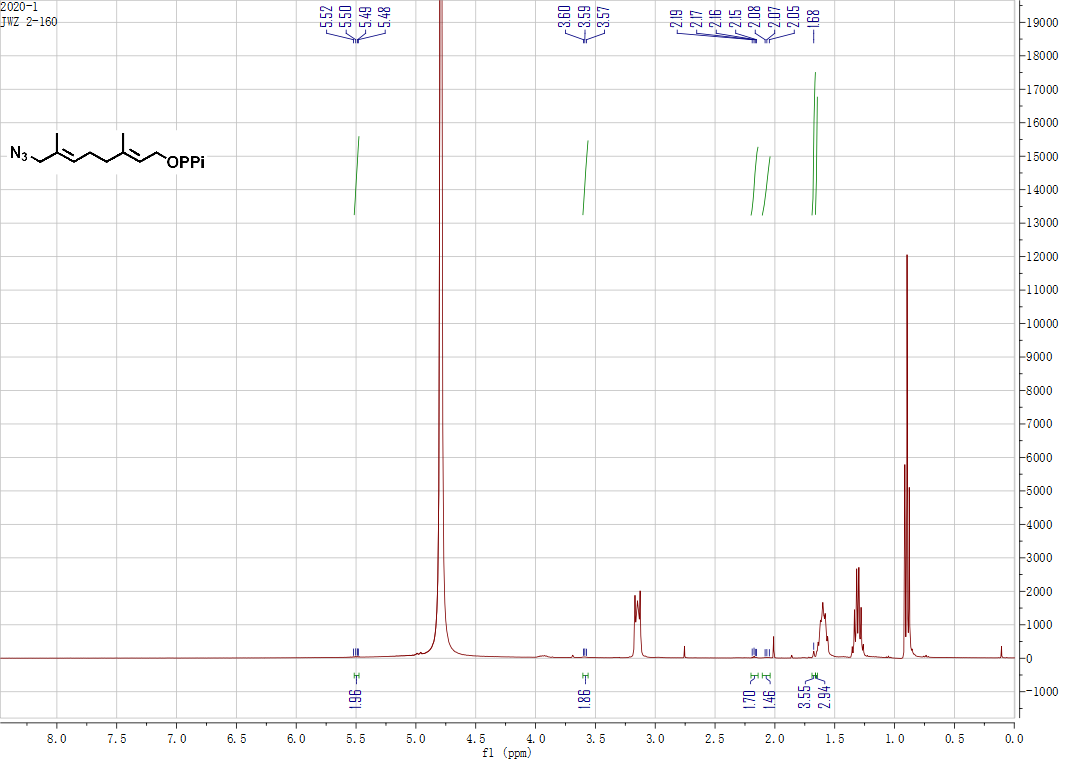

**Figure S25**. **^1^H NMR** spectra of pyrophosphate **Ge5**.

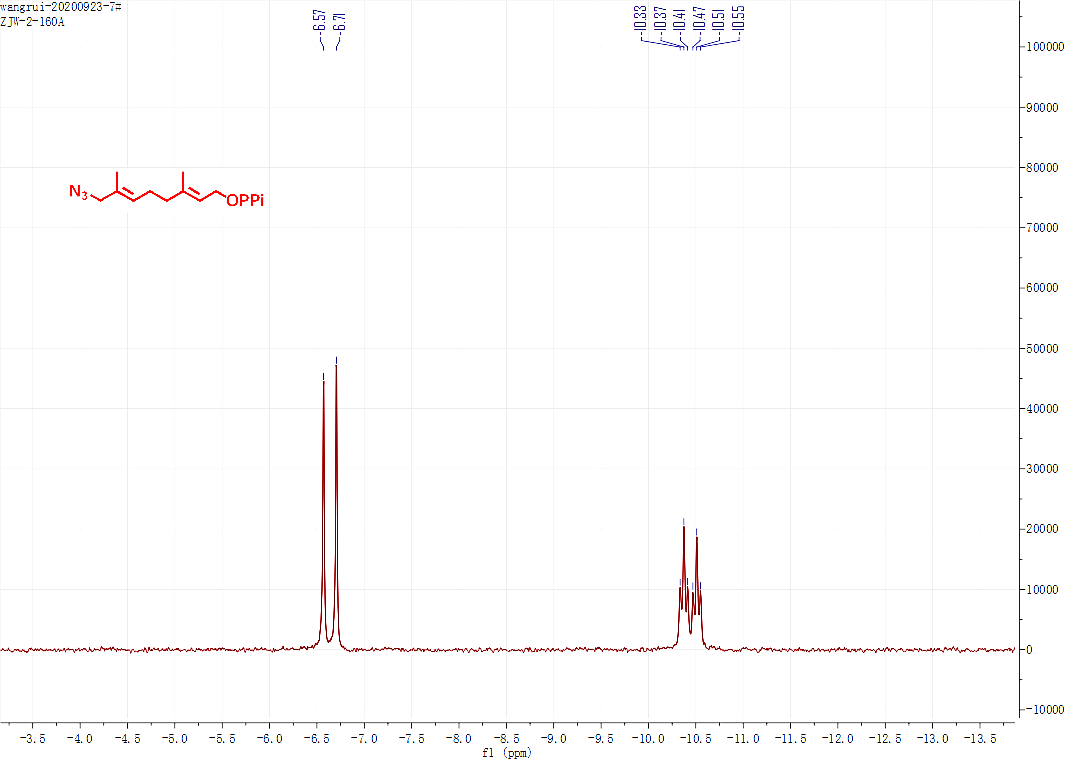

**Figure S26**. **^31^P NMR** spectra of pyrophosphate **Ge5**.

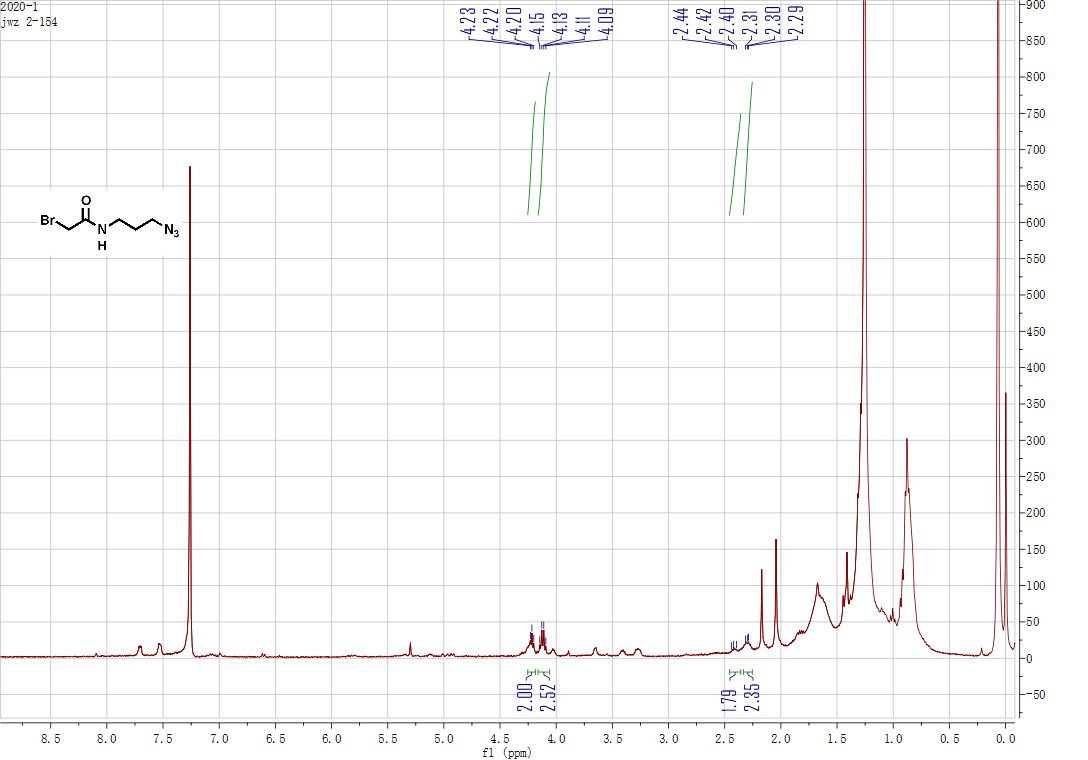

**Figure S27**. **^1^H NMR** spectra of azide **15.**

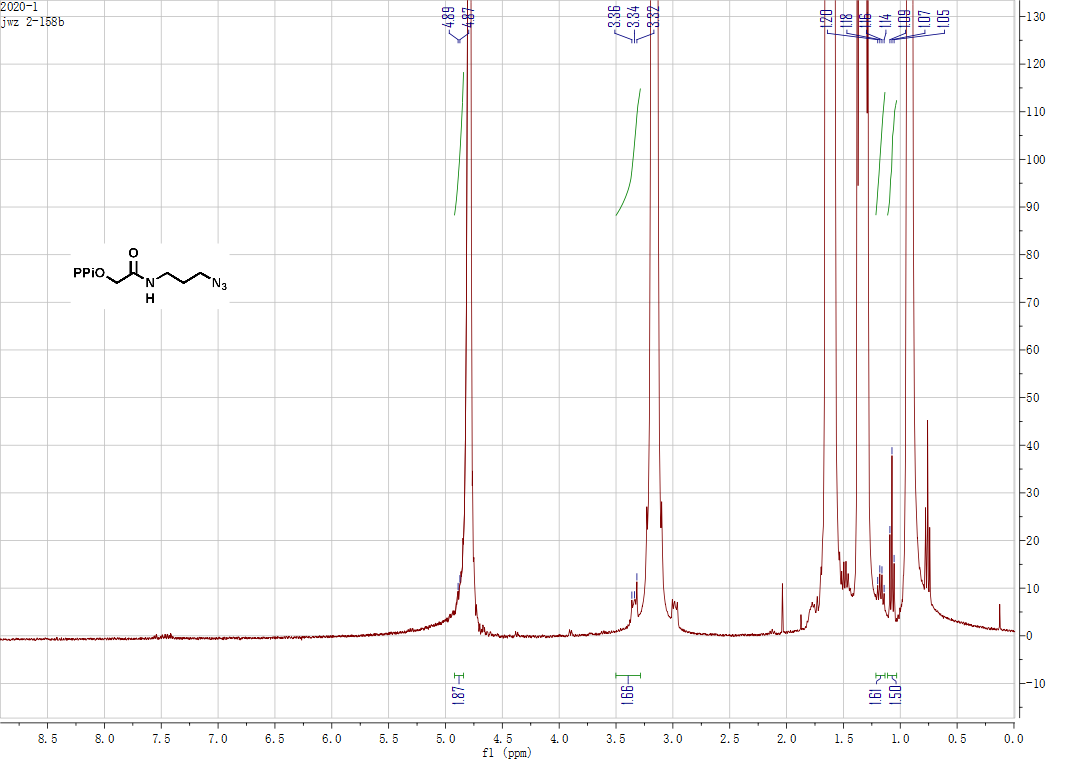

**Figure S28**. **^1^H NMR** spectra of pyrophosphate **Ge6**.

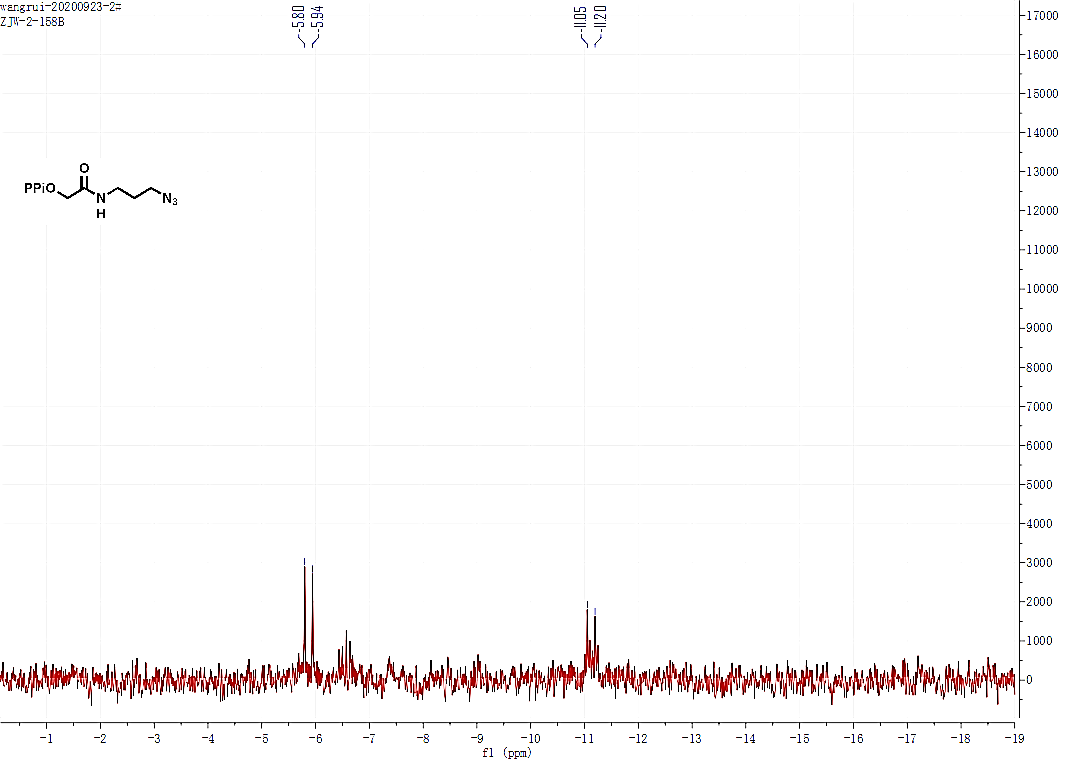

**Figure S29**. **^31^P NMR** spectra of pyrophosphate **Ge6**.

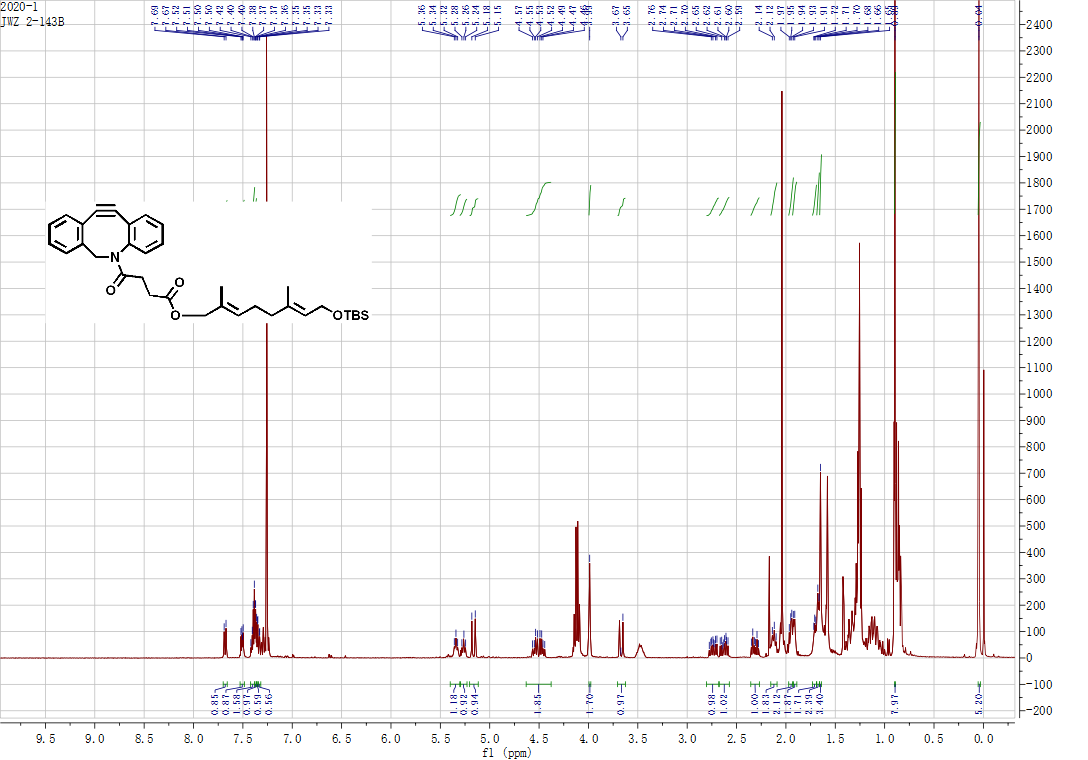

**Figure S30**. **^1^H NMR** spectra of compound **18**.

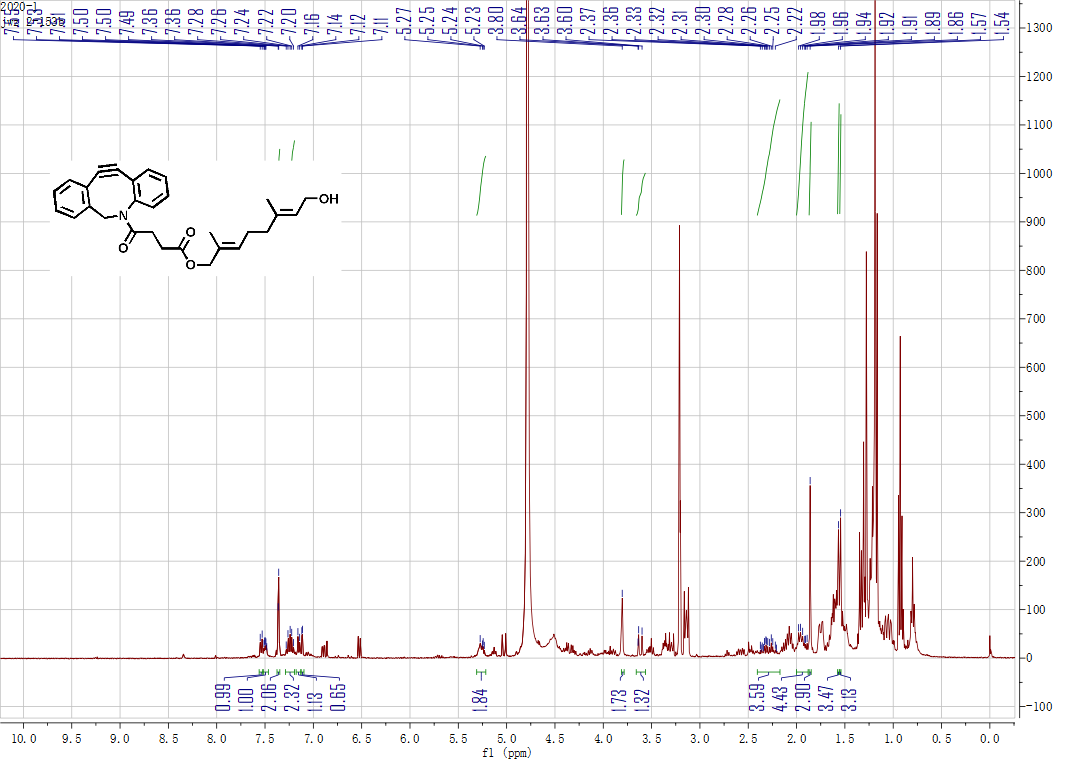

**Figure S31**. **^1^H NMR** spectra of alcohol **19**.

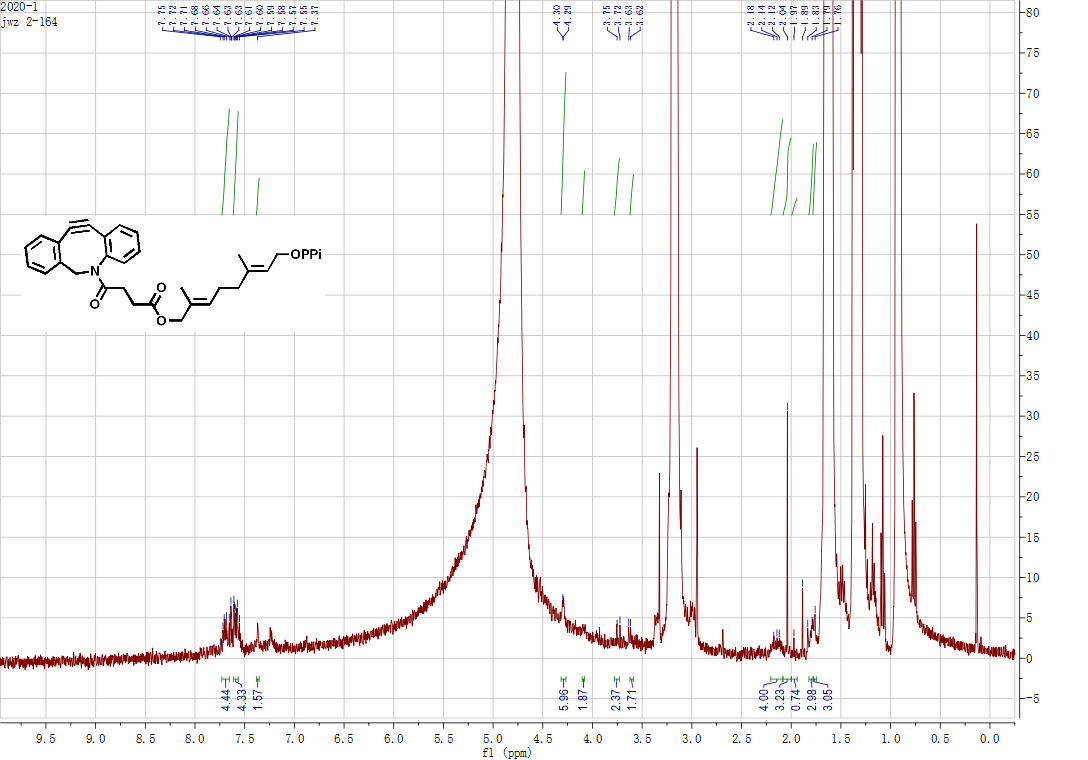

**Figure S32**. **^1^H NMR** spectra of pyrophosphate **Ge7**.

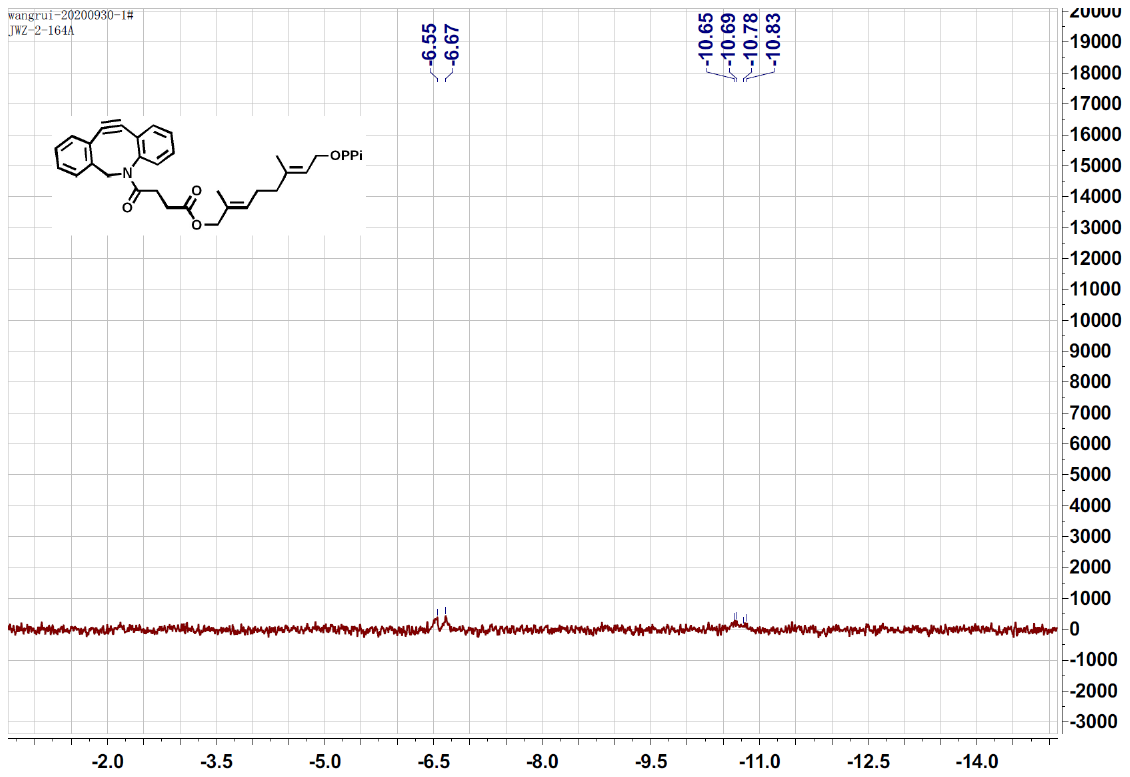

**Figure S33**. **^31^P NMR** spectra of pyrophosphate **Ge7.**

**SYNTHESIS OF THE STANDARD NUCLEOSIDES**

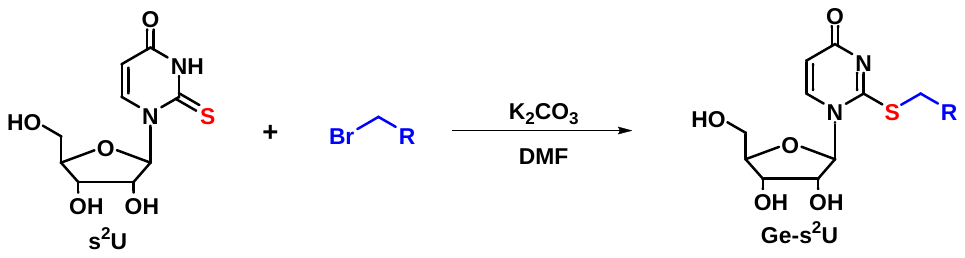

**Scheme S6**. Synthetic routes for standard nucleoside.

General procedure: To a stirred solution of **s^2^U** (13.0 mg, 0.05 mmol) in anhydrous DMF (0.1 mL) was added K_2_CO_3_ (0.2 mmol) at r.t. Bromide (0.2 mmol) was injected in one-potion and the mixture was stirred overnight at 85 ^o^C. The solvents were removed under *vacuo* and was further purified by HPLC to produce the **Ge-s^2^U**.

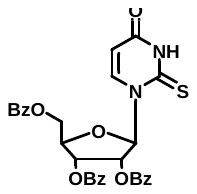

1.1 Synthesis of 2-thioxo-2,3-dihydropyrimidin-4(*1H*)-one (**21**).

To a stirred 4-((trimethylsilyl)oxy)-2-((trimethylsilyl)thio) pyrimidine (3.81 g, 29.8 mmol) in 1,1,1,3,3,3-Hexamethyldisilazane was added trimethyl chlorosilane (4.0 mL, 31.7 mmol). The resulted mixture was stirred at 135 ℃ of refluxed temperature overnight under N_2_ atmosphere until the mixture became clear. Solvent was removed through rotary evaporator and directly for next step.

(*2S,3R,4R,5R*)-2-acetoxy-5-((benzoyloxy)methyl)-tetrahydrofuran-3,4-diyldibenzoate (10.0 g, 19.82 mmol) dissolved in dichloroethane (80.0 mL) was cooled to -20℃ and added into previous product. Then the tin (IV) chloride (8.0 mL) was slowly added into the mixture. The resulted mixture was stirred at r.t. overnight under N_2_ atmosphere. The combined organic layers were washed with saturated aqueous Na_2_SO_4_, extracted with CH_2_Cl_2_, dried over Na_2_SO_4,_ and concentrated on rotary evaporator after using TLC to confirm that the starting material was consumed. Then the mixture was quenched with saturated NaHCO_3_, extracted with CH_2_Cl_2_ and dried over Na_2_SO_4_. The crude product was purified by silica gel flash chromatography using dichloromethane in ethyl acetate (20: 1) as eluent to give pure compound **21** (4.80 g).

**^1^H NMR** (400 MHz, CDCl_3_) δ 9.96 (s, 1H), 8.08 (d, *J =* 7.8 Hz, 3H), 8.00 (d, *J =* 7.8 Hz, 3H), 7.93 (d, *J =* 7.9 Hz, 3H), 7.70 (d, *J =* 8.2 Hz, 2H), 7.61 - 7.55 (m, 3H), 7.55 - 7.50 (m, 3H), 7.45 - 7.33 (m, 7H), 4.90 (d, *J =* 2.2 Hz, 1H), 4.86 (d, *J =* 2.2 Hz, 1H), 4.78 (dt, *J =* 4.8, 2.4 Hz, 2H), 4.70 (d, *J =* 3.0 Hz, 1H), 4.67 (d, *J =* 2.9 Hz, 1H). **^13^C NMR** (101 MHz, CDCl_3_) δ 176.09 (s), 165.95 (s), 165.30 (s), 165.00 (s), 158.97 (s), 139.04 (s), 129.95 (d, *J =* 17.6 Hz), 129.58 (s), 128.93 (s), 128.58 (s), 107.39 (s), 90.40 (s), 80.72 (s), 74.40 (s), 70.28 (s), 63.04 (s). **MS (ESI)**: Calculated [M+Na] ^+^ = 595.6, Found [M+Na] ^+^ = 595.3.

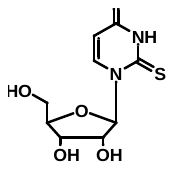

1.2 Synthesis of 1-((2R,3R,4S,5R)-3,4-dihydroxy-5-(hydroxymethyl) tetra hydrofuran-2-yl)-2-thioxo-2,3-dihydropyrimidin-4(*1H*)-one (**22**).

To a stirred solution of compound **11** (3.30 g, 5.76 mmol) in aqueous menthol (14.0 mL) was added sodium methoxide (6.1 mL, 34.6 mmol) at 0℃. The resulted mixture was stirred at r.t. under N_2_ atmosphere. Dowex 50WX8-400 was added to adjust the pH = 7 - 8. The bi-phase was filtered and reduced in evaporator rotary. Water and ethyl acetate were added to extract the crude product and wash off anisole. The mixture was lyophilized to dryness to afford crude product **12** in 57% yield.

**^1^H NMR** (400 MHz, CD_3_OD) δ 8.36 (d, *J* = 8.1 Hz, 1H), 6.60 (s, 1H), 5.95 (d, *J* = 8.1 Hz, 1H), 4.21 (dd, *J* = 4.9, 2.5 Hz, 1H), 4.16 - 4.12 (m, 1H), 4.09 - 4.04 (m, 1H), 3.96 (dd, *J* = 12.5, 2.2 Hz, 1H), 3.80 (dd, *J* = 12.4, 2.4 Hz, 1H). **^13^C NMR** (101 MHz, CDCl_3_) δ 178.39 (s), 163.24 (s), 143.40 (s), 107.17 (s), 95.63 (s), 86.45 (s), 77.23 (s), 70.56 (s), 61.62 (s). **HRMS (ESI)**: Calculated [M+Na] ^+^ = 283.0365, Found [M+Na] ^+^ = 283.0363.

1.3 Synthesis of 1-((2R,3R,4S,5R)-3,4-dihydroxy-5-(hydroxymethyl) tetrahydrofuran-2-yl)-2-(((E)-3,7-dimethylocta-2,6-dien-1-yl)-thio)-pyrimidin-4(*1H*)-one (**23**).

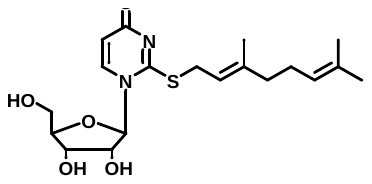

To a stirred solution of nucleoside **22** (5.0 mg, 0.019 mmol), geranyl bromide (13.3 mg, 0.057 mmol) and DIPEA (7.0 mg, 0.057 mmol) in MeOH (0.5 mL). The mixture was stirred overnight at r.t. under N_2_ atmosphere. The crude residue was purified by preparative thin layer chromatography and confirmed by ^1^H NMR and MS (ESI).

**^1^H NMR** (400 MHz, CDCl_3_) δ 8.23 (d, *J* = 8.0 Hz, 1H), 6.05 (s, 1H), 6.03 (s, 1H), 5.87 (d, *J* = 3.3 Hz, 1H), 5.58 - 5.57 (m, 1H), 5.51 (s, 1H), 5.06 (s, 1H), 4.86 (s, 1H), 4.37 (s, 1H), 4.07 (s, 1H), 4.06 (s, 1H), 2.00 - 1.99 (m, 2H), 1.99 - 1.97 (m, 2H), 1.75 (s, 3H), 1.74 (s, 3H). **MS (ESI)**: Calculated [M+H] ^+^ = 397.5, Found [M+Na] ^+^ = 398.2.

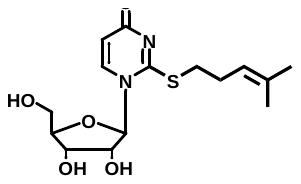

1.4 Synthesis of 1-((2R,3R,4S,5R)-3,4-dihydroxy-5-(hydroxymethyl)-tetrahydrofuran-2-yl)-2-((4-methylpent-3-en-1-yl)-thio)-pyrimidin-4-(*1H*)-one (**24**).

To a stirred solution of nucleoside **22** (5.0 mg, 0.0190 mmol), 5-bromo-2-methylpent-2-ene (13.3 mg, 0.0570 mmol) and DIPEA (7.0 mg, 0.057 mmol) in MeOH (0.5 mL). The mixture was stirred overnight at r.t. under N_2_ atmosphere. The residue was purified by preparative thin layer chromatography and confirmed by ^1^H NMR and mass spectra.

**^1^H NMR** (400 MHz, CDCl_3_) δ 8.18 (s, 1H), 6.19 (d, *J* = 5.3 Hz, 1H), 4.47 - 4.42 (m, 2H), 4.40 (t, *J* = 3.6 Hz, 1H), 4.36 (d, *J* = 4.2 Hz, 1H), 4.32 (d, *J* = 6.6 Hz, 1H), 4.13 - 4.01 (m, 2H), 3.43 (dd, *J* = 15.2, 11.8 Hz, 2H), 2.94 - 2.83 (m, 2H), 1.68 (d, *J* = 16.1 Hz, 6H). **MS (ESI)**: Calculated [M+Na] ^+^ = 365.4, Found [M+Na] ^+^ = 365.3.

1.5 Synthesis of *N*-(3-azidopropyl)-2-((1-((2R,3R,4S,5R)-3,4-dihydroxy-5-(hydroxymethyl)-tetrahydrofuran-2-yl)-4-oxo-1,4-dihydropyrimidin-2-yl)-thio) acetamide (**25**).

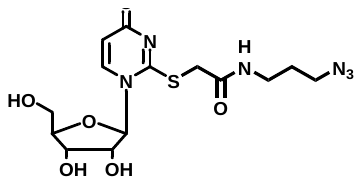

To a stirred solution of nucleoside **22** (5.0 mg, 0.0190 mmol), 5-bromo-2-methylpent-2-ene (13.3 mg, 0.0570 mmol) and DIPEA (7.00 mg, 0.057 mmol) in MeOH (0.5 mL). The mixture was stirred overnight at r.t. under N_2_ atmosphere. The residue was purified by preparative thin layer chromatography and confirmed by ^1^H NMR and MS (ESI).

**^1^H NMR** (400 MHz, CDCl_3_) δ 8.07 (d, *J* = 6.9 Hz, 1H), 5.78 (d, *J* = 3.4 Hz, 1H), 5.33 (s, 1H), 4.87 (dd, *J* = 12.9, 2.4 Hz, 1H), 4.77 (d, *J* = 2.7 Hz, 1H), 4.67 (dd, *J* = 12.5, 2.9 Hz, 1H), 3.69 - 3.55 (m, 1H), 2.20 (dd, *J* = 11.9, 5.1 Hz, 1H), 2.02 - 1.97 (m, 1H), 1.40 (d, *J* = 4.8 Hz, 2H), 1.30 (d, *J* = 5.3 Hz, 2H). **MS (ESI)**: Calculated [M+Na] ^+^ = 423.4, Found [M+Na] ^+^ = 423.0.

1.6 Synthesis of 1-((2R,3R,4S,5R)-3, 4-dihydroxy-5-(hydroxymethyl)-tetrahydrofuran-2-yl)-2-((3,7-dimethyloct-6-en-1-yl) thio)-pyrimidin-4 (*1H*)-one (**26**).

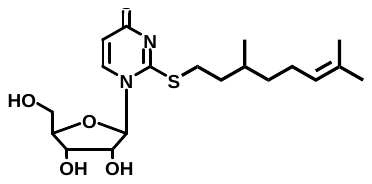

To a stirred solution of nucleoside **22** (5.0 mg, 0.019 mmol), 5-bromo-2-methylpent-2-ene (13.3 mg, 0.057 mmol) and DIPEA (7.0 mg, 0.057 mmol) in MeOH (0.5 mL). The mixture was stirred overnight at r.t. under N_2_ atmosphere. The residue was purified by preparative thin layer chromatography and confirmed by ^1^H NMR and MS (ESI).

**^1^H NMR** (400 MHz, CDCl_3_) δ 8.34 (d, *J* = 8.2 Hz, 1H), 6.74 (d, *J* = 8.4 Hz, 1H), 6.04 - 5.87 (m, 1H), 5.33 (s, 1H), 4.94 (dd, *J* = 26.2, 14.0 Hz, 1H), 4.66 (d, *J* = 12.8 Hz, 1H), 3.80 - 3.74 (m, 2H), 2.16 (s, 2H), 1.86 (s, 2H), 1.61 (s, 4H), 1.59 (s, 4H), 0.99 (d, *J* = 6.6 Hz, 3H). **MS (ESI)**: Calculated [M+H_2_O+H] ^+^ = 417.5, Found [M+H_2_O+H] ^+^ = 417.1.

**NMR Spectra II of Nucleosides**

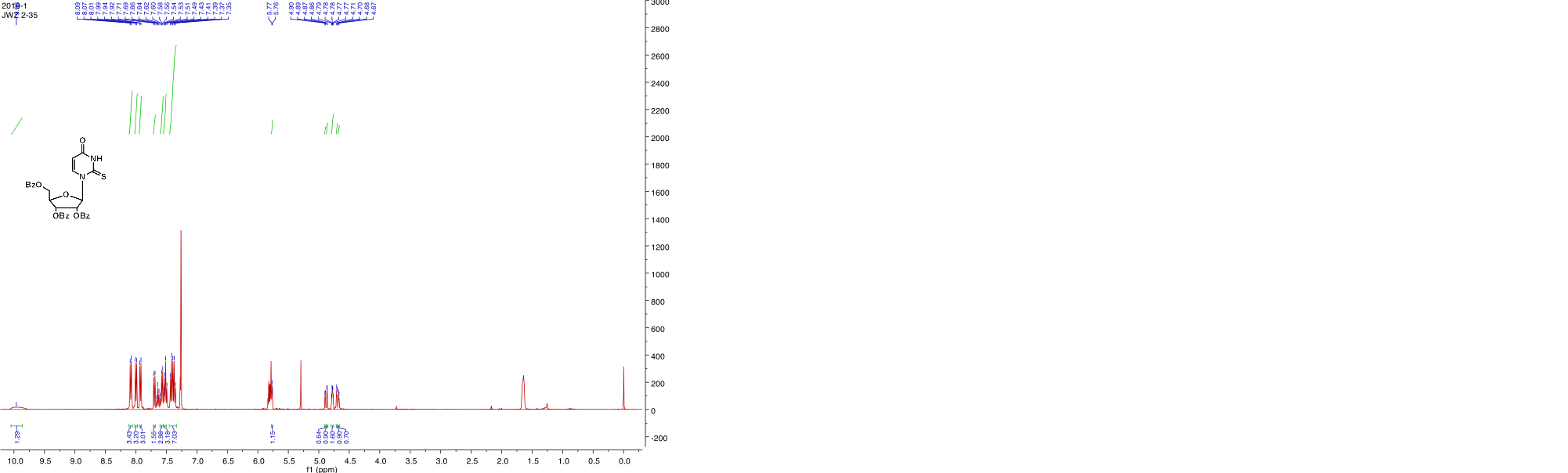

**Figure S34**. **^1^H NMR** spectra of **21**.

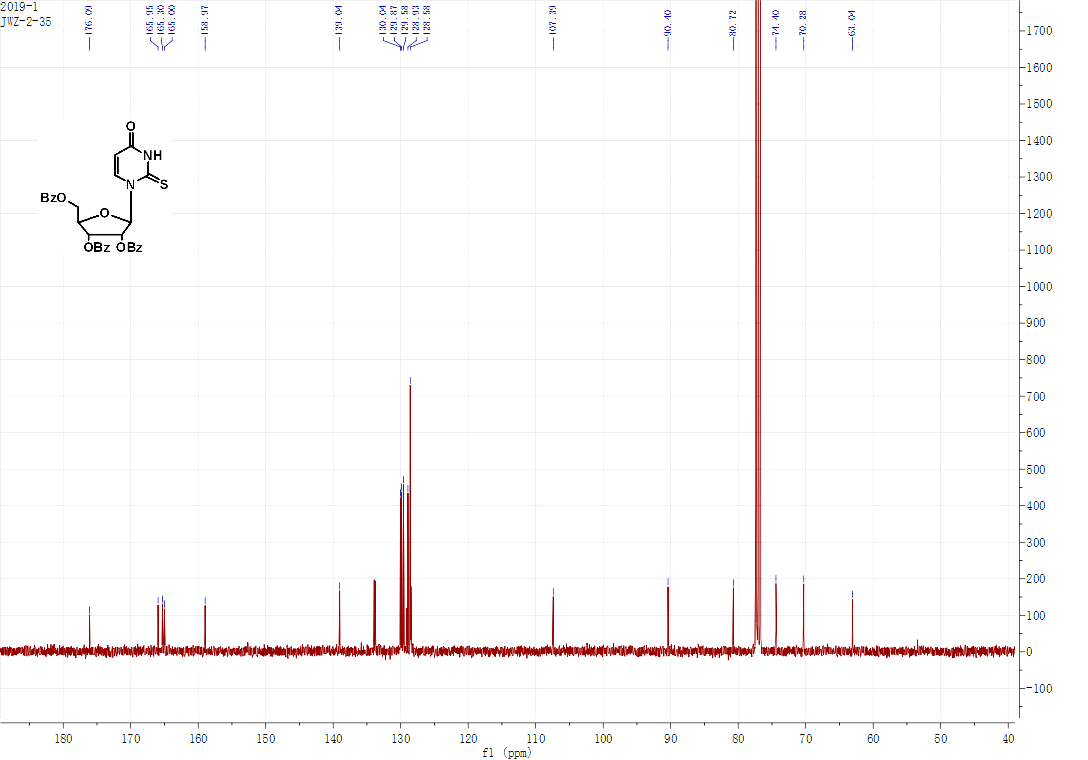

**Figure S35**. **^13^C NMR** spectra of **21**.

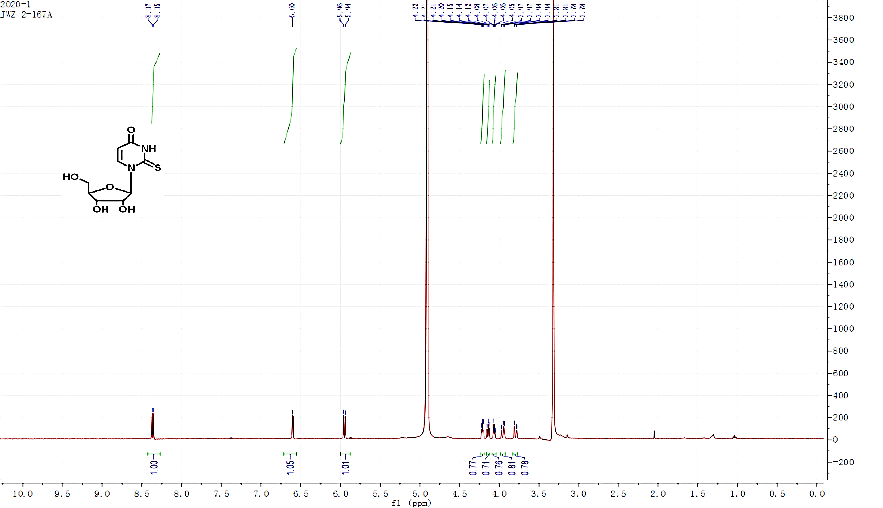

**Figure S36**. **^1^H NMR** spectra of nucleoside **22**.

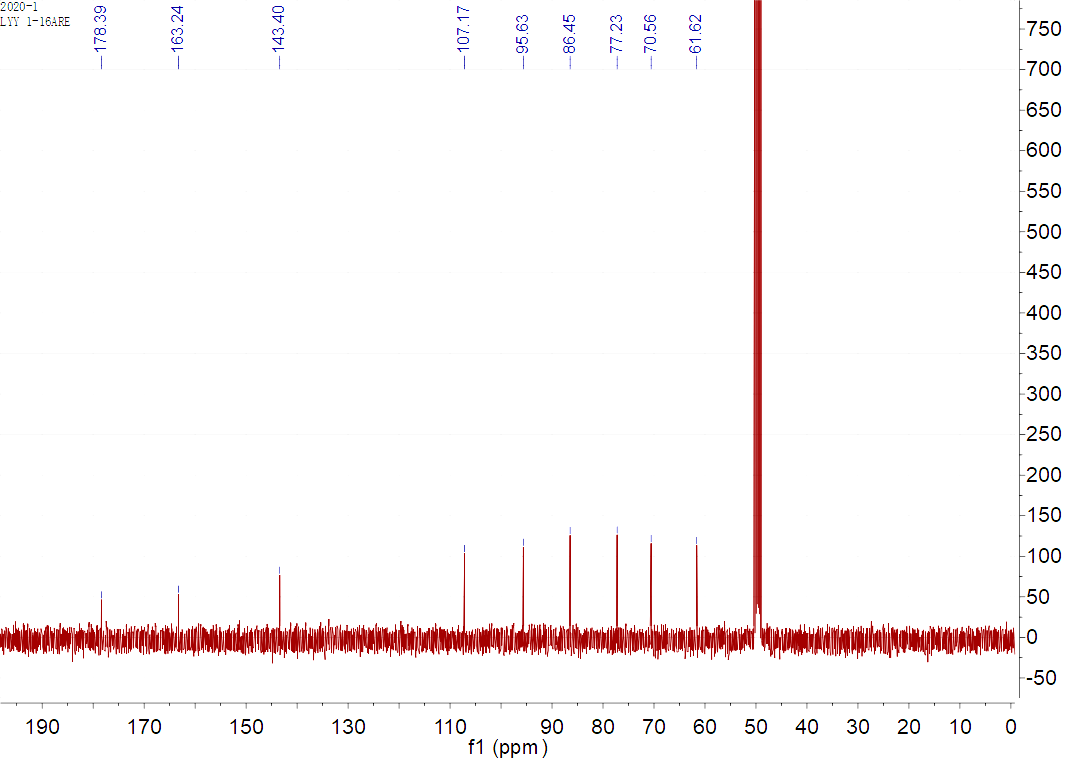

**Figure S37**. **^13^C NMR** spectra of nucleoside **22**.

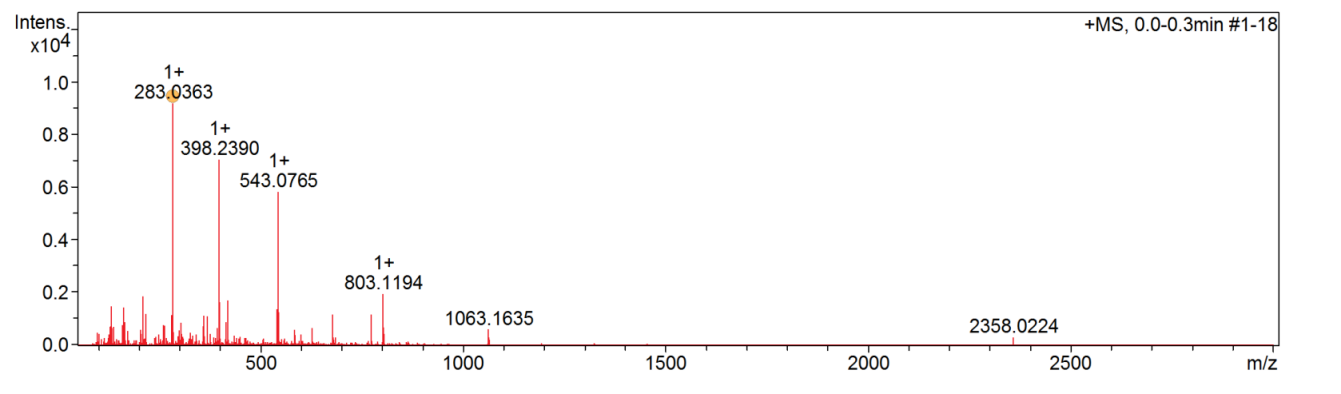

**Figure S38**. Mass spectra of nucleoside **22**.

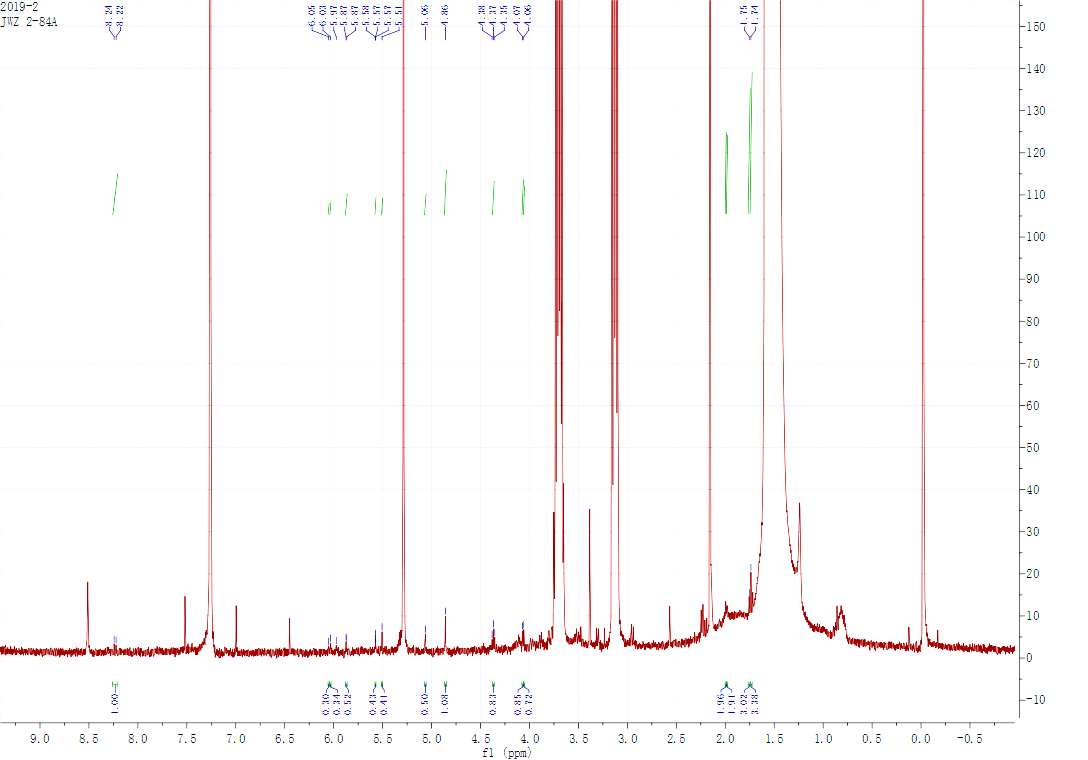

**Figure S39**. **^1^H NMR** spectra of nucleoside **23**.

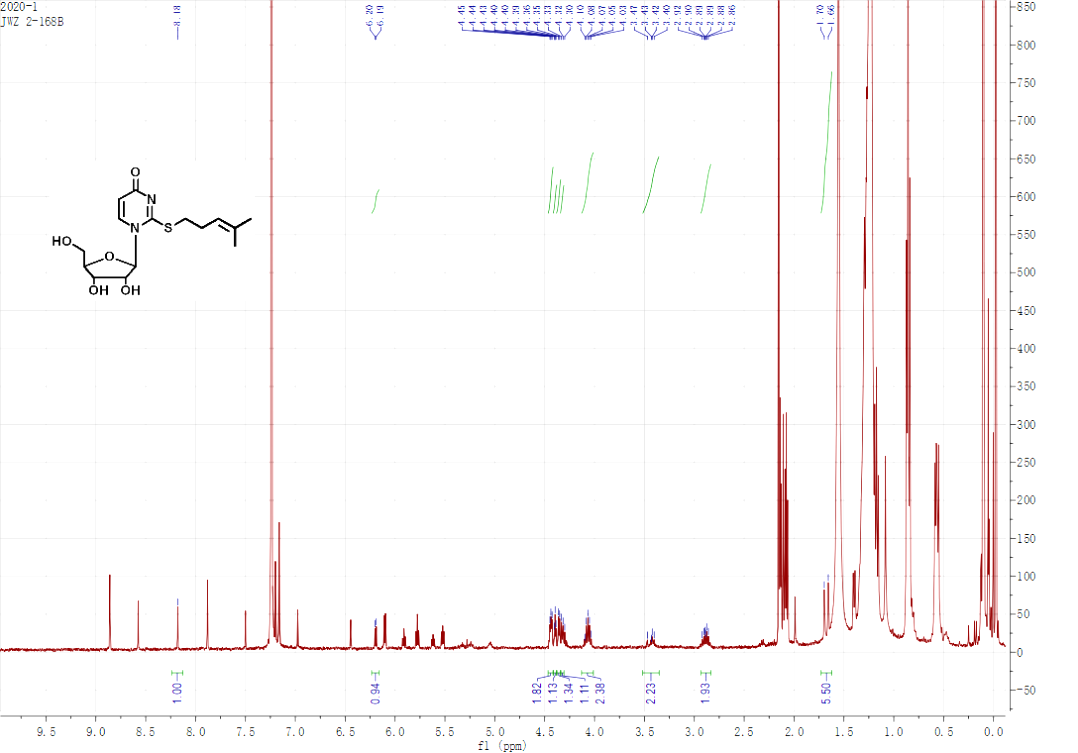

**Figure S40**. **^1^H NMR** spectra of nucleoside **24**.

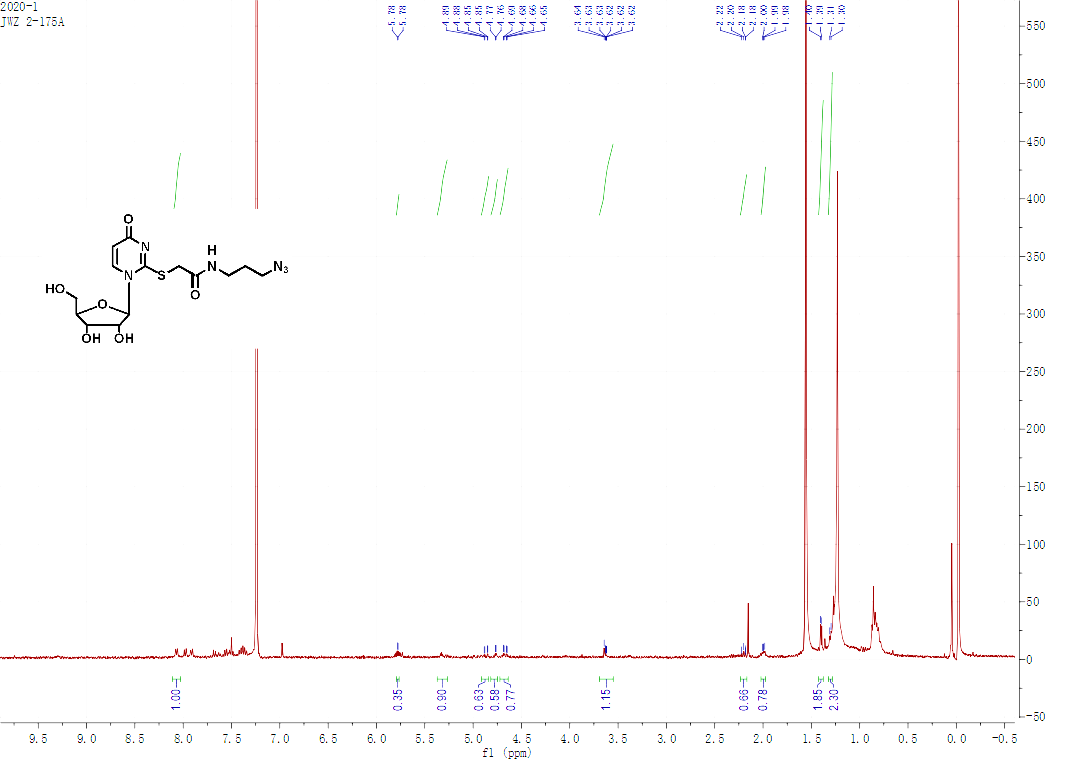

**Figure S41**. **^1^H NMR** spectra of nucleoside **25**.

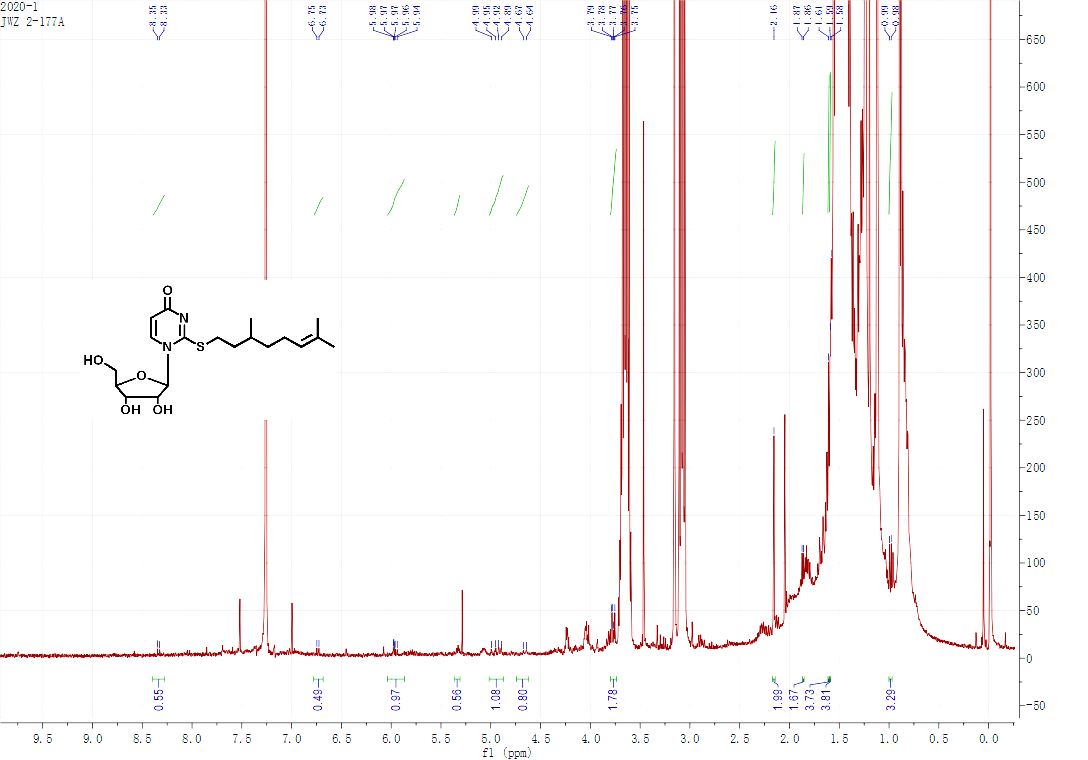

**Figure S42**. **^1^H NMR** spectra of nucleoside **26**.

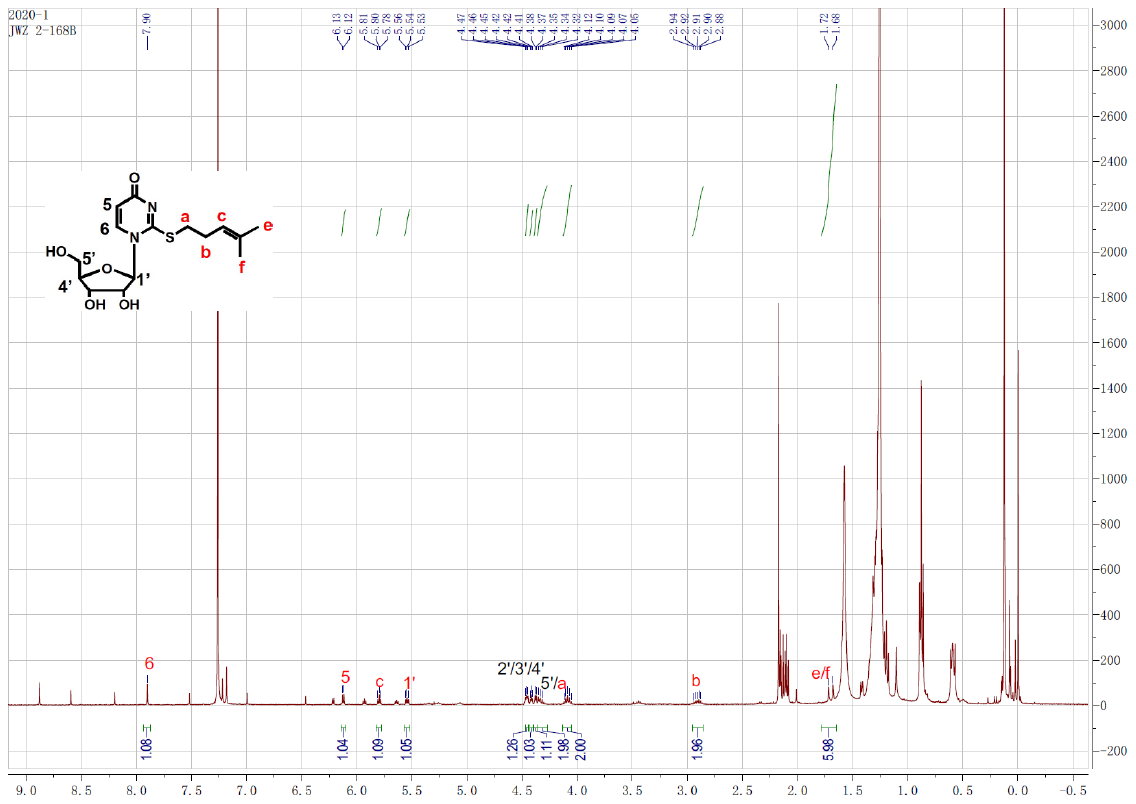

**Figure S43.** **^1^H NMR** spectrum of nucleoside **Ge3-s^2^U**.

**CONSTRUCTION OF *SELU* ENZYME**

(1) Amplification of gene *SelU****.***

A 1095 bp sequence of gene *SelU* was amplified using primers FselU and RselU (Table S2) with genome of *E. coli* DH5α as amplification template. Reaction mixture (Table 3) was amplified in PCR instrument (98 ºC 2 min, 98 ºC 1 s, 60 ºC 30 s, 72 ºC 1 min, 72 ºC 10 min, 12 ºC 10 min, 30 cycles). Product of amplified gene *SelU* was showed in Fig. S44a. and purified by *Takara* minibest agarose gel DNA extraction kit ver.4.0 (101 ng/ *μ*l).

**Table S2**. Primers FselU and RselU.

| Primers | Sequences (5'-3')^a^ | | | Restriction sites |
| --- | --- | --- | --- | --- |
| F*selU* | CGCGGATCCATGCAAGAGAGACACACGGAACAGGTG | *BamH*I | | |
| R*selU* | CCCAAGCTTTTACCGCGCCTTAACCCATTCCGCC | | *Hind*III | |

**Table S3.** PCR Mixtures.

|  | ddH_2_O | 144 *μ*l | |
| --- | --- | --- | --- |
|  | 5×buffer | 40 *μ*l | |
| HS primesta room temperature taq | | | 2 *μ*l |
|  | dNTP | 4 *μ*l | |
|  | F*selU* | 4 *μ*l | |
|  | R*selU* | 4 *μ*l | |
|  | template | 2 *μ*l | |
|  | total | 200 *μ*l | |

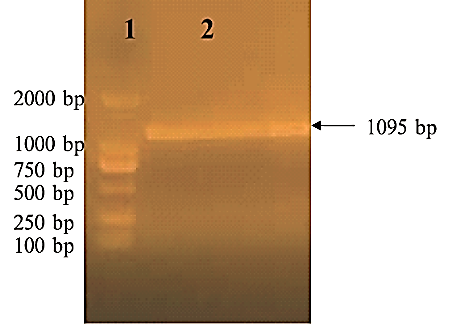

**Figure S44a.** Gene Production of *SelU*. Lane 1: DL 2000 maker. Lane 2: gene *SelU* amplified from genome of *E. coli* DH5α.

(2) Construction of plasmid pMD19T-*SelU*.

The purified product of *SelU*-A was ligated into pMD19-T vector with reaction mixture (Solution I, 5 *μ*l, *SelU*-A 4.5 *μ*l, pMD19-T 0.5 *μ*l) overnight at 16 ºC. Then transfer connection product of *SelU*-A and pMD19-T into DH5α and cultivated on Amp (ampicillin) resistance LB plate with 1.5% agar overnight at 37 ºC. Result of transformation was detected by PCR (98 ºC 2 min, 98 ºC 1 s, 60 ºC 15 s, 72ºC 1 min, 72 ºC 10 min, 12 ºC 10 min, 30 cycles) with reaction mixture (ddH_2_O 13.2 *μ*l, 5×buffer 4 *μ*l, HS primestart taq 0.3 *μ*l, dNTP 0.5 *μ*l, FselU 1 *μ*l, RselU 1 *μ*l, template 5 *μ*l) (**Fig. S44b**).

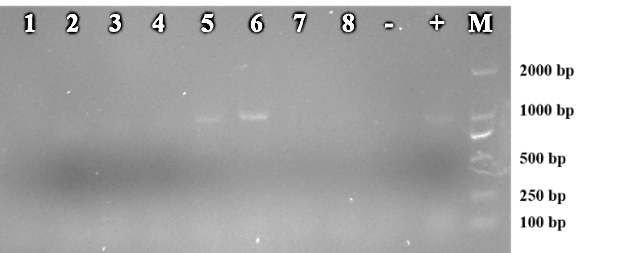

**Figure S44b.** *SelU* gene PCR product. Lane 1-8: PCR product of gene *SelU* with template of *SelU*-T/ DH5α. -: PCR product of gene *SelU* with template of ddH_2_O. +: PCR product of gene *SelU* with template of *E. coli* DH5α genome. M: DL 2000 maker.

**Figure S44b** showed that sequence of *SelU*-T/ DH5α (5) and *SelU*-T/ DH5α (6) were possible right. Result of sequencing detection suggested that sequence of *SelU*-T/ DH5α (5) was totally right (**Fig. S44c**).

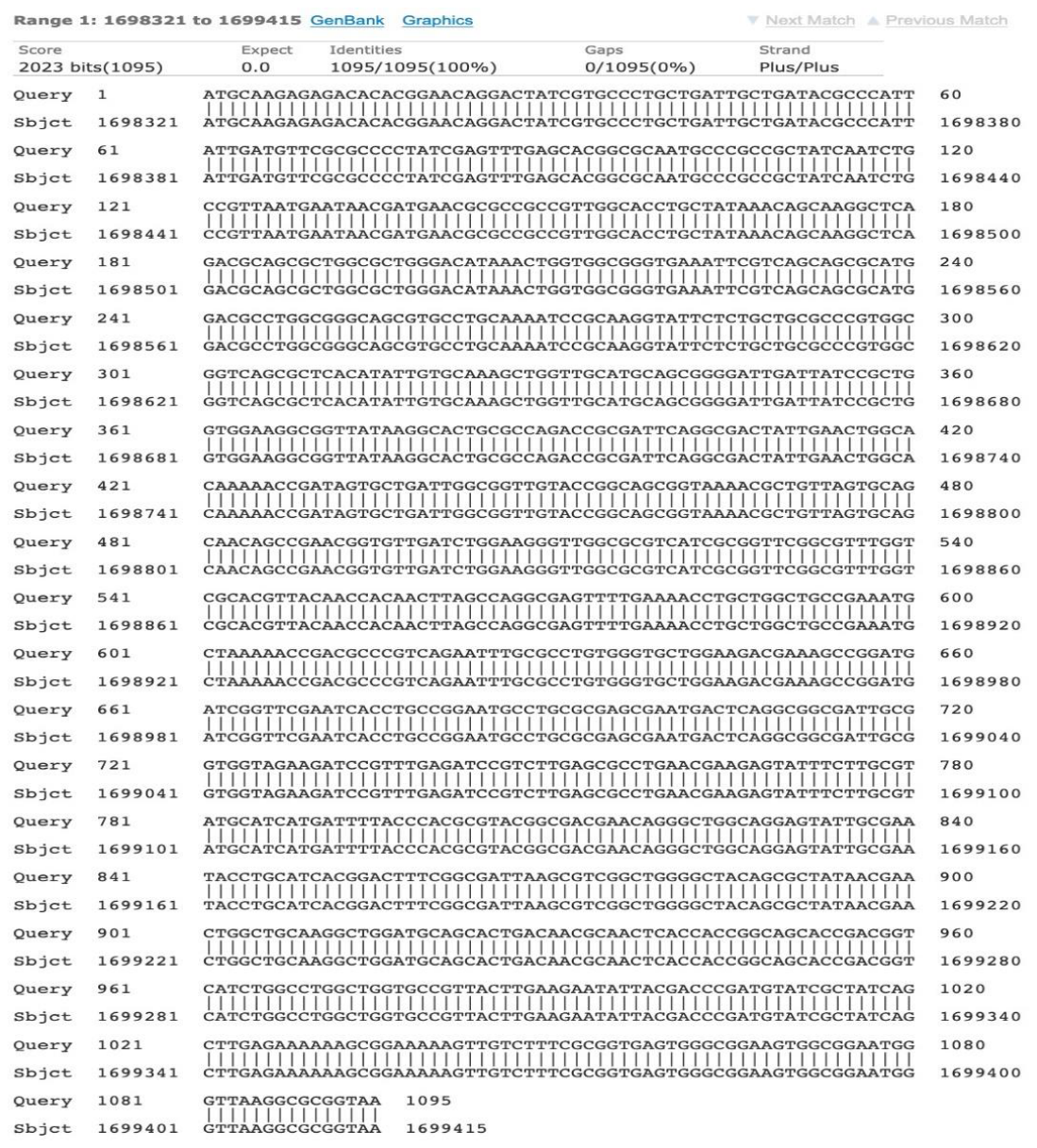

**Figure S44c.** Result of sequencing detection. A. Sequencing detection result of *SelU*-T/ DH5α (5). B. Sequencing detection result of *SelU*-T/ DH5α (6).

(3) Extraction of plasmid pMD19T-*SelU* and pET28b.

Bacterial strain of pMD19T-*SelU* (DH5α) and pET28b (DH5α) were incubated in 37 ºC overnight. 2.0 mL bacterial solution was collected and plasmid was extracted with *Takara* minibest plasmid purification kit ver. 4.0. And concentration of plasmid was detected by Nanodrop. Concentration of pMD19T-*SelU* and pET28b were 172 ng/ *μ*l and 96 ng/ *μ*l respectively.

(4) Digestion of plasmid pMD19T-*SelU* and pET28b.

The plasmid of *SelU*-pMD19T and pET28b were digested with *BamHI/ Hind**III.* The reaction mixture (10× buffer 10 *μ*l, plasmid 80 *μ*l, *BamHI* 5 *μ*l, *HindIII* 5 *μ*l) was incubated in 37ºC overnight and digested sequence of *SelU* and pET28b were collected with Takara minibest DNA fragment purification kit ver.4.0.

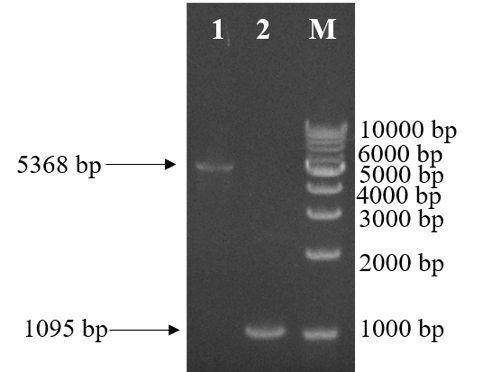

**Figure S44d.** Digestion of plasmid *SelU*-pMD19T and pET28b.

Lane 1: pET28b was digested with *BamHI/ HindⅢ.* Lane 2: *SelU* was digested with *BamHI/ HindⅢ*. M:1 kbp maker.

(5) Connection of gene *SelU* and pET28b.

The sequence of *SelU* was digested with *BamHI/ HindIII* and ligated into the same digested vector pET-28b with mixture (10 × buffer 1 *μ*l, ligase 0.5 *μ*l, *SelU* 7 *μ*l, pET-28b 1.5 *μ*l) overnight at 16 ºC. Then the product of mixture was transformed into DH5α, cultivated on Kan (kanamycin) resistance LB plate with 1.5% agar overnight at 37 ºC.

(6) PCR detection of pET28b-*SelU* (DH5α) transformant.

The transformants of pET28b-*SelU* (DH5α) were cultivated in Kan (kanamycin) resistance LB in 37 ºC overnight. Sequence of gene *SelU* was amplified with genome of pET28b-*SelU* (DH5α) transformants as amplification template. Reaction mixture (ddH_2_O 13.2 *μ*l, 5× buffer 4 *μ*l, HS primestart taq 0.3 *μ*l, dNTP 0.5 *μ*l, FselU 1 *μ*l, RselU 1 *μ*l, template 5 *μ*l) was amplified in PCR instrument (98 ºC 2 min, 98 ºC 1 s, 60 ºC 15 s, 72 ºC 1 min, 72 ºC 10 min, 12 ºC 10 min, 30 cycles) (**Fig. S44e**). The result of PCR detection showed below,

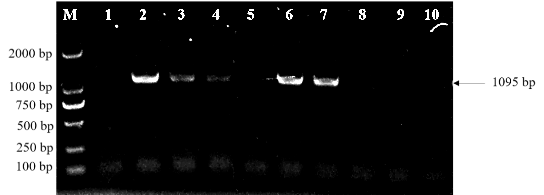

**Figure S44e.** Production of amplified gene *SelU*. M: DL 2000 maker. Lane 1-10: PCR product of gene *SelU* with template of pET28b-*SelU* DH5α.

The result of PCR detection showed number 2, 3, 4, 6, 7 of pET28b-*SelU* (DH5α) transformants were right. In order to confirm this conclusion, we will detect these five transformants by digestion.

(7) Digestion detection of pET28b-*SelU* (DH5α) transformant.

Plasmids of pET28b-*SelU* (DH5α) transformants (NO. 2, 3, 4, 6, 7) were extracted and digested with *BamHI/ HindIII*. The reaction mixture (10× buffer 1 *μ*l, plasmid 8 *μ*l, BamHI 0.5 *μ*l, *HindIII* 0.5 *μ*l) was incubated in 37 ºC overnight (**Fig. S44f**).

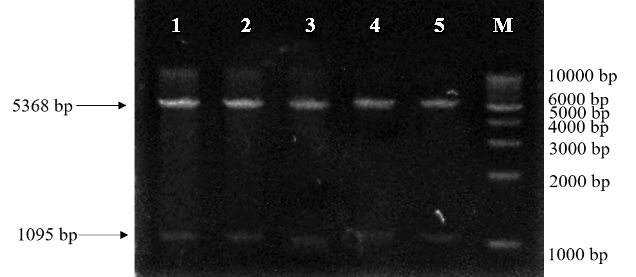

**Figure S44f.** Digestion of pET28b-*SelU* (DH5α) transformant plasmids.

Lane 1-5: Plasmids of pET28b-*SelU* (DH5α) transformants (2, 3, 4, 6, 7) were digested with *BamHI*/ *HindIII*. M: 1 kbp maker.

(8) Construction of *SelU* expression strain.

The plasmid of pET28b-*SelU* was transformed into BL21, cultivated on Kan (kanamycin) resistance LB plate with 1.5% agar overnight at 37 ºC. And the transformants of pET28b-*SelU* (BL21) were detected by PCR. Sequence of gene *SelU* was amplified with genome of pET28b-*SelU* (BL21) transformants as amplification template. Reaction mixture (ddH_2_O 13.2 *μ*l, 5× buffer 4 *μ*l, HS primestart taq 0.3 *μ*l, dNTP 0.5 *μ*l, FselU 1 *μ*l, RselU 1 *μ*l, template 5 *μ*l) was amplified in PCR instrument (98 ºC 2 min, 98 ºC 1 s, 60 ºC 15 s, 72 ºC 1 min, 72 ºC 10 min, 12 ºC 10 min, 30 cycles) (**Fig. S44g**). The result of PCR detection showed number 1-5 of pET28b-*SelU* (BL21) transformants were right.

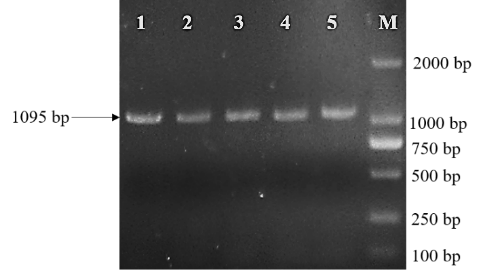

**Figure S44g.** Production of amplified gene *SelU*.

Lane 1-5: PCR product of gene *SelU* with template of pET28b -*SelU* / BL21. M: DL 2000 maker.

(9) Expression of *SelU* in small scale.

Strain pET28b-*SelU* (BL21) (1-5) were cultivated in Kan (kanamycin) resistance LB overnight at 37 ºC and transferred 40 *μ*l of strain liquid into fresh 4 mL Kan (kanamycin) resistance LB, cultivated at 220 rpm/ min, 37ºC for about 3 h to OD = 0.6. Then the 4.0 mL of strain liquid was separated to two equant parts, cultivated at 220 r.p.m./ min, 28 ºC for 5 h. One part was cultivated with 1.0 mmol/ L IPTG and another part was cultivated without IPTG. 2 mL of strain liquid was collected and suspended in 40 *μ*l of PBS buffer and 10 *μ*l of loading buffer, and boiled for 20 min. 10 *μ*l of samples were detected by 12% SDS-PAGE (**Fig. S44h**).

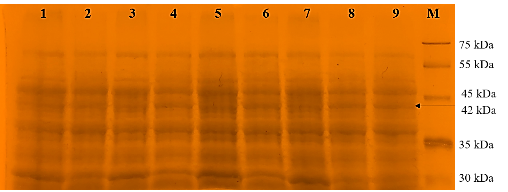

**Figure S44h.** Detected of pET28b-*SelU* (BL21) (1-5) by SDS-PAGE.

Lane 1: pET28b-*SelU* (BL21) (1) cultivated without IPTG. Lane 2: pET28b-*SelU* (BL21) (1) cultivated with 1 mmol/L IPTG. Lane 3: pET28b-*SelU* (BL21) (2) cultivated without IPTG. Lane 4: pET28b-*SelU* (BL21) (2) cultivated with 1 mmol/L IPTG. Lane 5: pET28b-*SelU* (BL21) (3) cultivated without IPTG. Lane 6: pET28b-*SelU* (BL21) (3) cultivated with 1 mmol/L IPTG. Lane 7: pET28b-*SelU* (BL21) (4) cultivated without IPTG. Lane 8: pET28b-*SelU* (BL21) (4) cultivated with 1 mmol/ L IPTG. Lane 9: pET28b-*SelU* (BL21) (5) cultivated with 1 mmol/ L IPTG. M: Clearly stained protein ladder.

(10) Expression of *SelU*.

Strain pET28b-*SelU* (BL21) was cultivated in Kan (kanamycin) resistance LB overnight at 37 ºC and transferred 2.0 mL of strain liquid into fresh 200 mL Kan (kanamycin) resistance LB, cultivated at 220 r.p.m./ min, 37 ºC for about 3 h to OD = 0.6. Then the 200 mL of strain liquid was cultivated with 1 mmol/ L IPTG at 220 r.p.m./ min, 28ºC for 5 h.

(11) Lysis of bacteria solution.

Strain liquid of pET28b-*SelU* (BL21) was collected and suspended in 10 mL PBS buffer. Then the suspended liquid was lysed with ultrasonication (work 5 s and pause 5 s) for 1 h.

(12) Purification of *SelU* protein.

*Step 1*. Equilibrate the His_6_ Ni Gravity Column (1.0 ml) and all buffers to the working temperature. (Perform purifications at r.t. or at 4 °C.)

*Step 2*. Wash the column with 5-10 column volumes of His_6_ Ni Equilibration Buffer (50 mM sodium phosphate, 6 M guanidine-HCl, 300 mM NaCl, 20 mM imidazole; pH = 7.4).

*Step 3*. Add the clarified sample to the column and carefully connect the top stopper to the top of the column. Allow target protein to bind by slowly inverting the column for 1 h.

*Step 4*. Install the column in a vertical position and let the resin settle at the bottom of the column.

*Step 5*. Wash the column with 10 column volumes of His_6_ Ni Equilibration Buffer followed by 10 column volumes of His_6_ Ni Wash Buffer (50 mM sodium phosphate, 6.0 M guanidine-HCl, 300 mM NaCl, 40 mM imidazole; pH = 7.4).

*Step 6*. Elute the target protein with approximately 10 column volumes of Elution Buffer (50 mM sodium phosphate, 6 M guanidine-HCl, 300 mM NaCl, 300 mM imidazole; pH = 7.4) and collect 1 ml once in a tube.

(13) Detection of *SelU* after purification.

Collect 40 *μ*l of purified *SelU* protein, add 10 *μ*l of loading buffer, boil for 20 minutes, and take 5 *μ*l of sample for 10% SDS-PAGE detection (**Fig. 44i**).

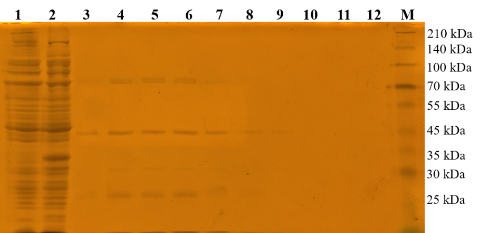

**Figure S44i.** Detection of *SelU* purification by SDS-PAGE. Lane 1: Liquid supernatant after ultrasonication. Lane 2: Liquid flowed from Ni-Sepharose columns. Lane 3-12: *SelU* eluted from Ni-Sepharose columns (1-10). M: Clearly stained protein ladder.

(14) Detection of *SelU* purification.

40 *μ*l of purified *SelU* protein were collected and added 10 *μ*l of loading buffer, boiled for 20 min. 5 *μ*l of samples were detected by 10% SDS-PAGE.

(15) Confirmation of K-12 MG1655.

Sequence of gene *SelU* was amplified with genome of *E. coli* K-12 MG1655, DH5α and ddH_2_O as amplification template. Reaction mixtures (ddH_2_O 23 *μ*l, 5🞨 Tsingke master mix 25 *μ*l, FselU 1 *μ*l, RselU 1 *μ*l, template 5 *μ*l) were amplified in PCR instrument (94 ºC 5 min, 94 ºC 30 s, 60ºC 30 s, 72 ºC 1 min, 72 ºC 10 min, 12 ºC 10 min, 30 cycles) (**Fig. S44j**).

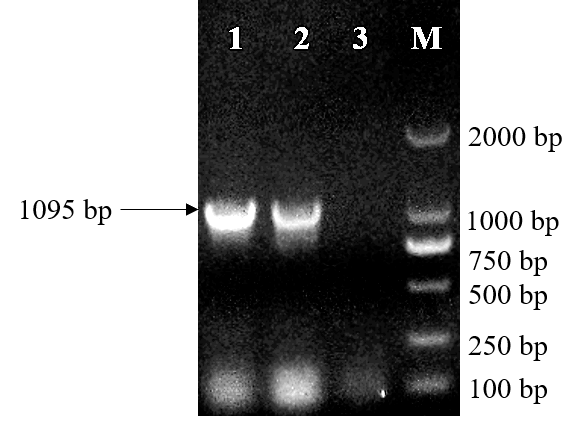

**Figure S44j.** Production of Amplified Gene *SelU*.

Lane 1: PCR product of gene *SelU* with template of K-12 MG1655 genome. Lane 2: PCR product of gene *SelU* with template DH5α genome. Lane 3: PCR product of gene *SelU* with template of ddH_2_O. M: DL 2000 maker.

**STRUCTURE ELUCIDATION OF *SELU* ENZYME AND ITS MOLECULAR DOCKING ANALYSIS WITH PYROPHOSPHATE VARIANTS**

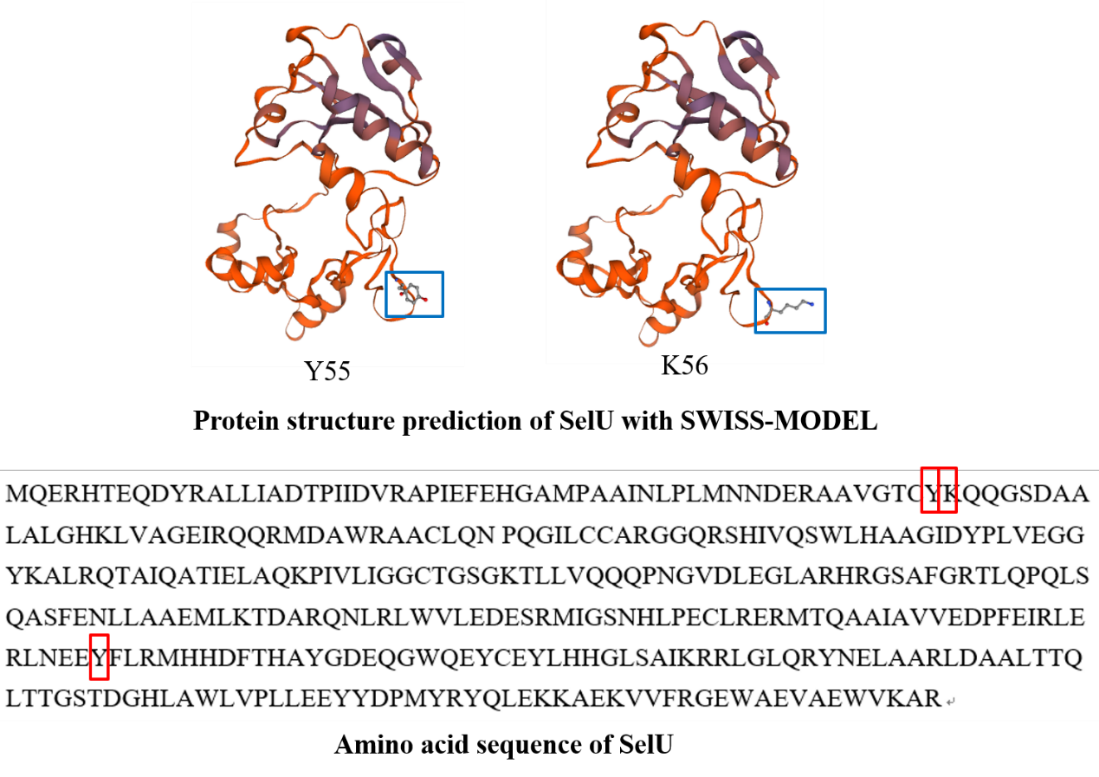

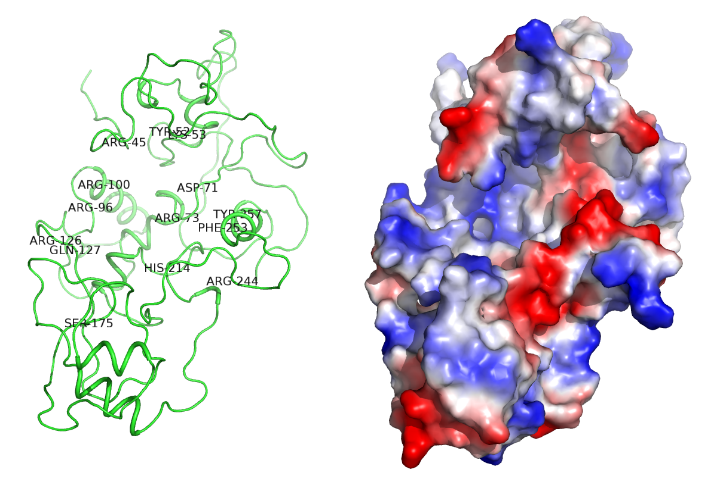

**Figure S45.** Results of *SelU* structure prediction using SWISS-MODEL.

**Prediction of *SelU* structure using *ALPHAFOLD*.**

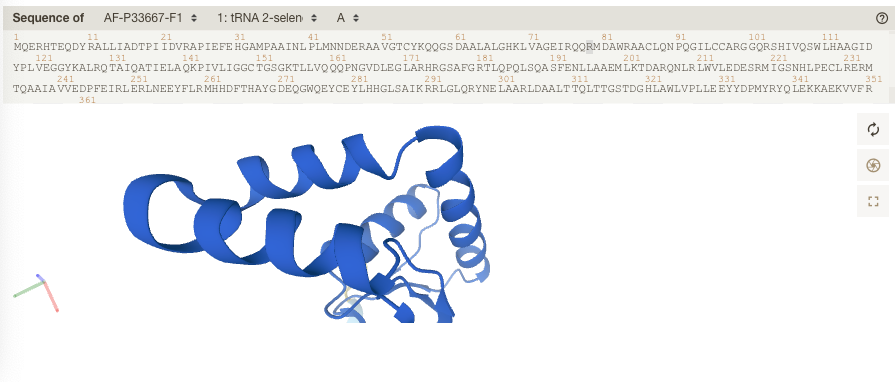

**Figure S46.** Results of *SelU* structure prediction using *AlphaFold*.

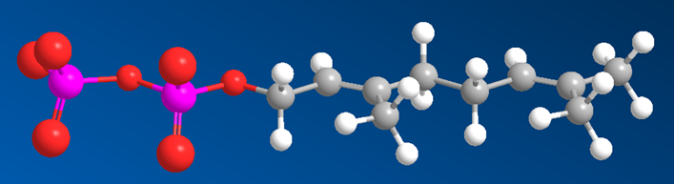
***Ge1***

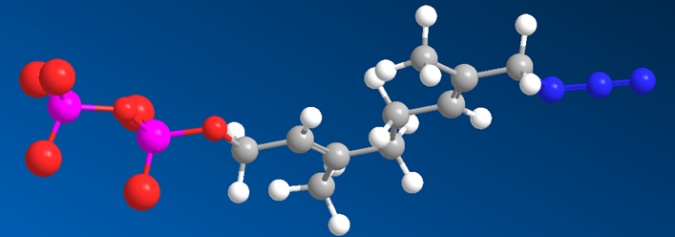
***Ge5***

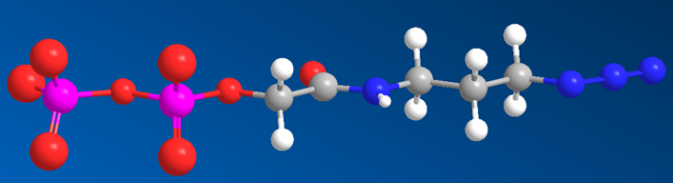
***Ge6***

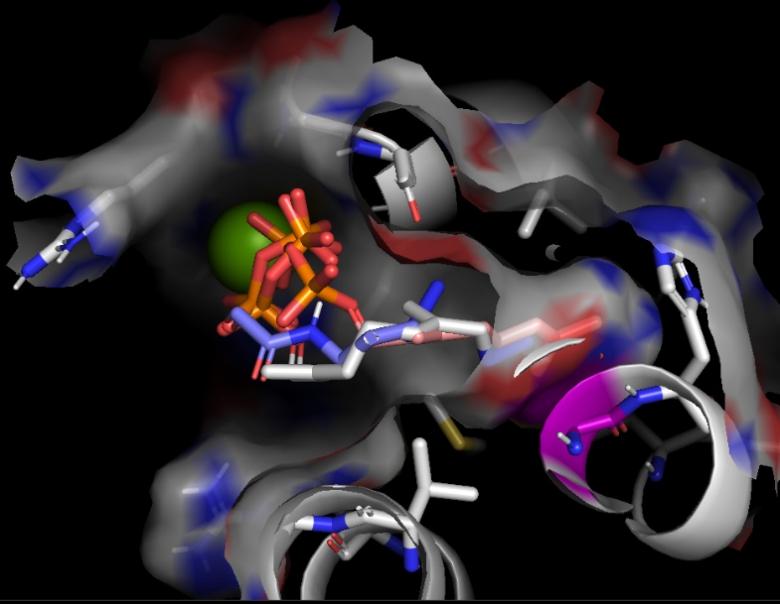

**Figure S47**. Molecular docking analysis of *SelU* with pyrophosphate variants **Ge5-6** with **Ge1** using *Autodock Vina.*

***SELU*-MEDIATED GERANYLATION OF tRNA IN THE FLUORESCENT LABELING INVESTIGATIONS**

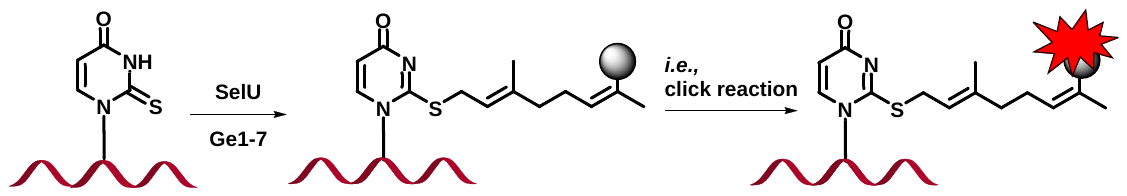

**Scheme S7.** Overall illustration of the labeling by metabolic methods.

***General procedure***: Partially purified *SelU*-His_6_ (15 *µ*g, ca. 0.35 nmol), 20 *µ*g (ca. 3.74 nmol) tRNA and geranyl pyrophosphate ammonium salt (Sigma, 5 eq.) was dissolved in 100 *µ*l of buffer containing 10 mM Tricine-KOH, pH = 7.2, 0.2 mM dithiothreitol (DTT), and 100 mM MgCl_2_, and the sample was incubated at 25 ^o^C for 24 h. The Ge-tRNA product was monitored by RP-HPLC using a Kinetex C_18_ column (5 *µ*, 100A; in the gradient of buffer B: 0-10 min 0% B; 10-40 min 0-35% B; 40-45 min 35-100% B; 45-50 min, 100% B; 50-55 min, 100-0% B; 55-60 min, 0% B. Buffer A: 0.1 M CH_3_CO_2_NH_4_; pH = 6.8. Buffer B: 0.1 M CH_3_CO_2_NH_4_ with 40% CH_3_CN; the collected fraction was desalted and the geranylated product was tested by fluorescent labeling or MALDI-TOF MS.

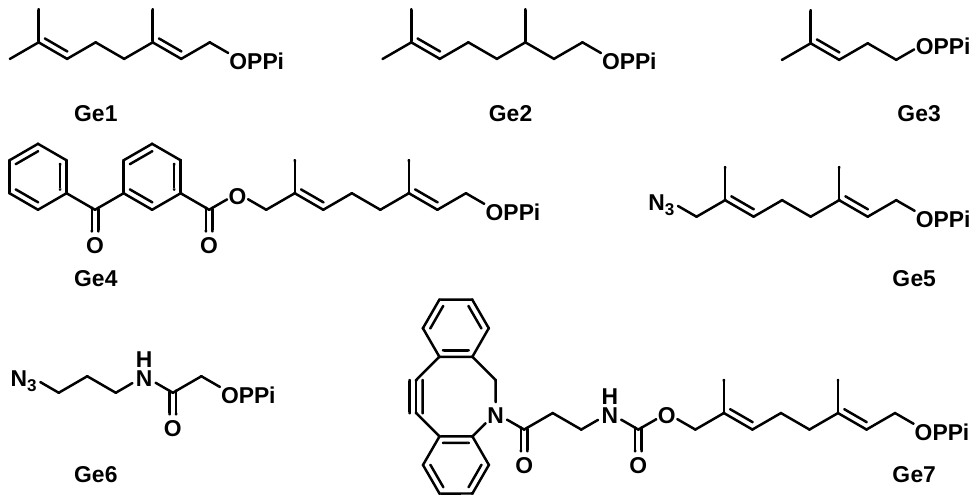

**Scheme S8.** Overall illustration of the fluorescently tagging medicated by *SelU*. **Ge1** for positive control. **Ge2-3** for nucleosides analysis experiment. **Ge5-7** for fluorescent labeling studies. **Ge4** for photo-mediated cross-linking experiment.

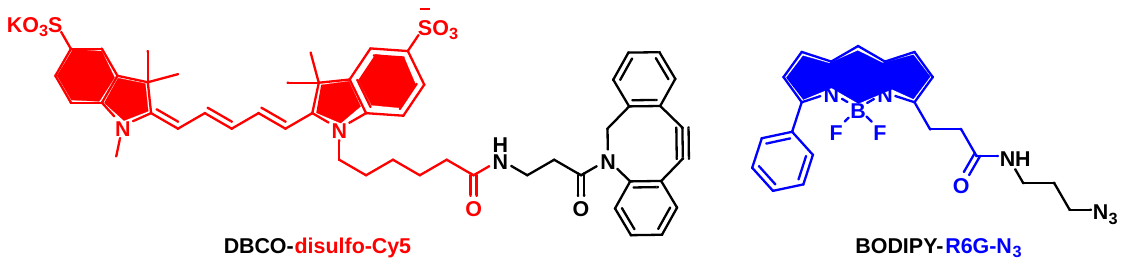

**Figure S48.** Fluorescent dyes used herein for tagging of geranylated RNA.

(1) Fluorescent labeling investigations (**Ge5-7** substrates).

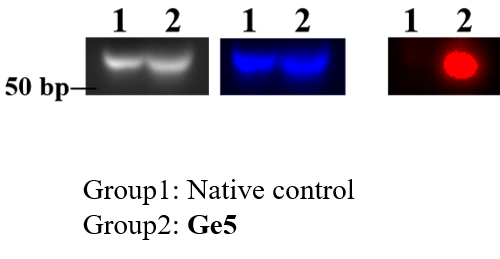

**Figure S49.** *SelU*-catalyzed tRNA geranylation with pyrophosphate **Ge5** as substrate, the in-gel fluorescent image results.

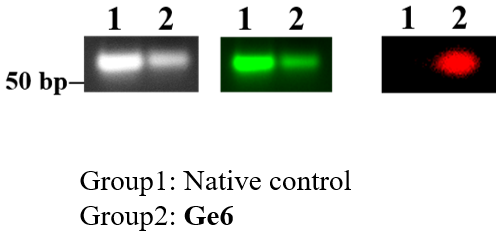

**Figure S50.** *SelU*-catalyzed tRNA geranylation with pyrophosphate **Ge6** as substrate, the in-gel fluorescent image results.

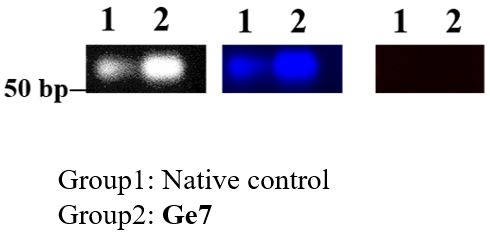

**Figure S51**. *SelU*-catalyzed tRNA geranylation with pyrophosphate **Ge5-7** as substrate, the in-gel fluorescent image results. **Ge5-6** and **Ge7** substrates in the fluorescent labeling study. PTAD-DBCO-Cy5 was used for **Ge5** or **Ge6**. BODIPY was used for **Ge7**. Blue channel (E.X. 492 nm. E.M. 507 nm), red channel (E.X. 649 nm. E.M. 660 nm) and green channel (nm).

(2) Hydrolysis analysis by *RNase T1*.

*RNase T1* is an endoribonuclease that specifically degrades single-stranded RNA at G residues. It cleaves the phosphodiester bond between 3’-guanylic residue and the 5’-OH residues of adjacent nucleotide with the formation of corresponding intermediate 2’, 3’-cyclic phosphate. The reaction products are 3’-GMP and oligonucleotides with a terminal 3’-GMP. *RNase T1* does not require metal ions for activity.

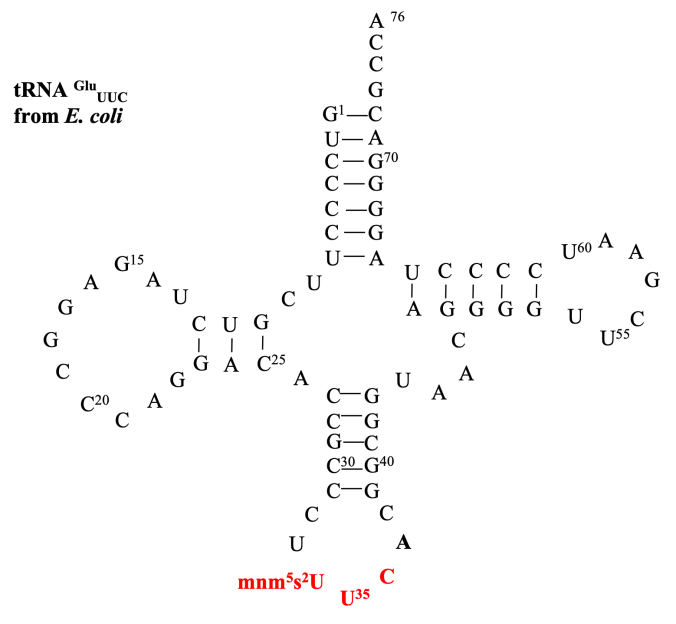

**Figure S52**. Structure details of tRNA^Glu^_UUC_ from *E. coli* and its cleavable G site by use of *RNase T1*. (5’-GUCCC CUUCG UCUAG AGGCC CAGGA CACCG CCCUU UCACG GCGGU AACAG GGGUU CGAAU CCCCU AGGGG ACGCC A-3’, 24.441 kDa).

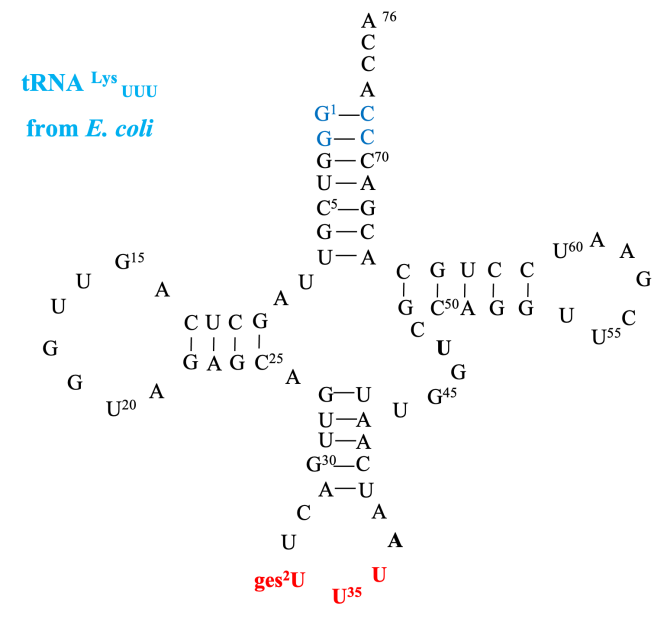

**Figure S53**. Structure details of tRNA^Lys^_UUU_ from *E. coli* and its cleavable G site by use of *RNase T1*. (5’-GGGUC GUUAG CUCAG UUGGU AGAGC AGUUG ACUUU UAAUC AAUUG GUCGC AGGUU CGAAU CCUGC ACGAC CCACC A-3’, 24,441 kDa).

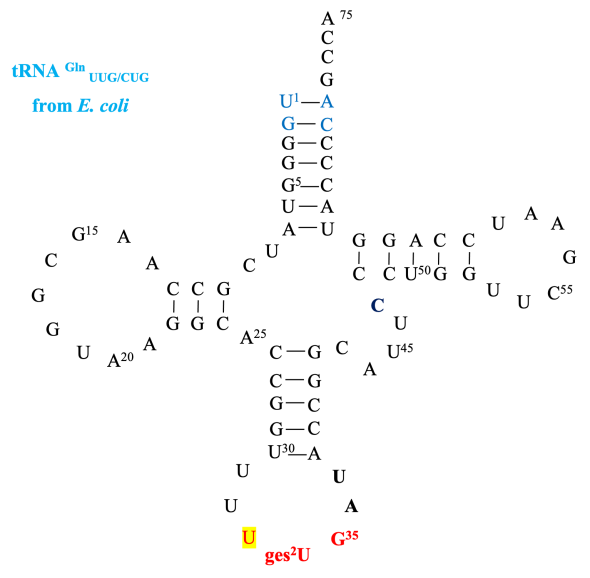

**Figure S54**. Structure details of tRNA^Gln^_UUU/CUG_ from *E. coli* and its cleavable G site by use of *RNase T1*. (5’-UGGGG UAUCG CCAAG CGGUA AGGCA CCGGU UUUUG AUACC GGCAU UCCCU GGUUC GAAUC CAGGU ACCCC AGCCA-3’, 24,124 kDa).

**Table S4**. Mass results analysis of tagged tRNA.

| Entries | **tRNA^Glu^_UUC_-A** | | **tRNA^Lys^_UUU_-B** | | **tRNA^Gln^_UUU/CUG_-C** | |
| --- | --- | --- | --- | --- | --- | --- |
|  | Sequence | M.W. | 5’-UUGACU**U**UUAAUCAAUUp-3’ | 5375 | Sequence | M.W. |
|  | 5’-CCCU**U**UCACp-3’ | 2792 | 5’-UUGACU**U**UUAAUCAAUUGp-3’ | 5720 | 5’-CCCU**U**UCACp-3’ | 2792 |
|  | 5’-CCCUUUCACGp-3’ | 3137 | 5’-UUGACUUUUAAUCAAUUGGUCp-3’ | 6677 | 5’-CCCUUUCACGp-3’ | 3137 |
|  | 5’-CCCU**U**UCACGGCp-3’ | 3787 | 5’-ACU**U**UUAAUCAAUUGp-3’ | 4763 | 5’-U**U**UUUGAUACCp-3’ | 3469 |
|  | 5’-CCCU**U**UCACGGCGp-3’ | 4132 | 5’-ACU**U**UUAAUCAAUUGGUCp-3’ | 5719 | 5’-U**U**UUUGAUACCGp-3’ | 3814 |
| Modification | **Ge5** vs U, Δm = 193 **I**  **Ge6** vs U, Δm = 127 **II**  **Ge7** vs U, Δm = 438 **III** | | | | | |
|  | **Found MS** | | **Calculated MS** | | Δm | |
| **Ge5** | 3398.8 | | 3330.0 (**A2 + I** or **C2 + I**) | | 68.8 Da | |
| **Ge6** | 3268.8 | | 3264.0 (**A2 + II** or **C2 + II**) | | 4.8 Da | |
| **N.C.** | 3216.0, 3293.0 | | **3137 (A2** or **C2)** | | 79.0, 156 Da | |
| Note: Several modifications have been reported on the codon loop of varied tRNA species. The targeted mass could be varied due to those modification occurred. | | | | | | |

(3) Hydrolysis nucleosides identifications (**Ge2-3** substrates).

**Table S5**. HPLC condition: **ges^2^U** (21 min).

| **Entry** | **t/min** | **Flow (mL/min)** | **A%** | **B%** |
| --- | --- | --- | --- | --- |
| 1 | 0 | 6 | 100 | 0 |
| 2 | 20 | 6 | 0 | 100 |
| 3 | 28 | 6 | 0 | 100 |
| 4 | 32 | 6 | 100 | 0 |
| 5 | 35 | 6 | 100 | 0 |
| 6 | 37 | 6 | 100 | 0 |
| **A** | H_2_O+0.1%HCCOH | | | |
| **B** | CH_3_CN+0.1%HCOOH | | | |

***Results analysis.***

*(a). +ESI, blank solvent first-order MS (+ESI, background).*

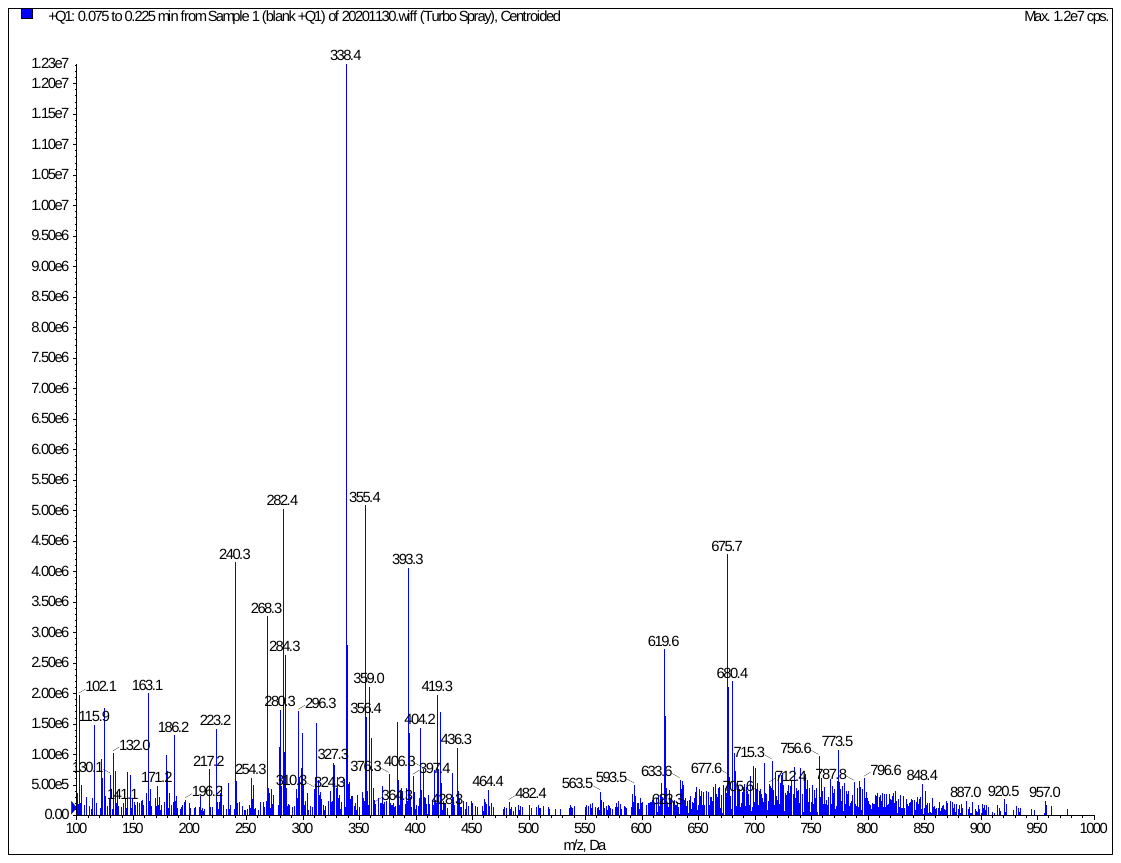

*(b). -ESI, blank solvent first-order MS (-ESI, background).*

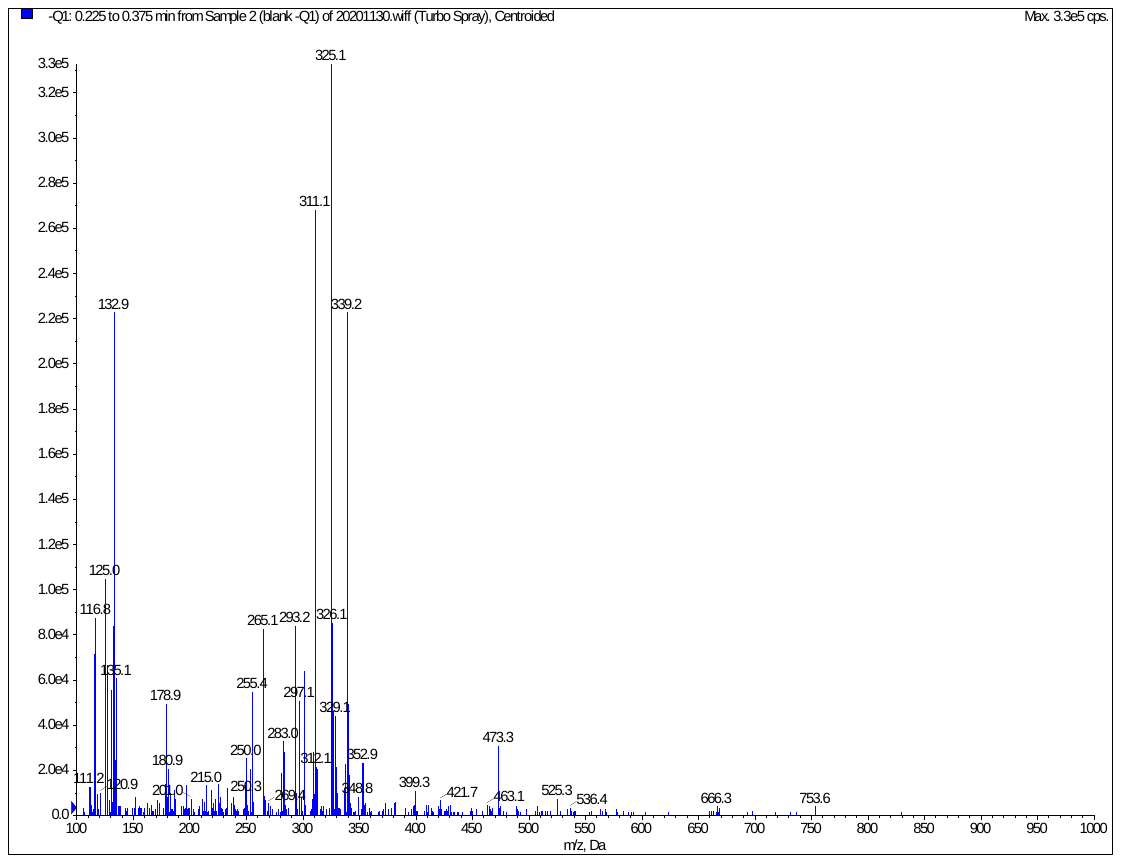

*(c).* *Ge-s^2^U sample (+ESI). Model: first-order MS.*

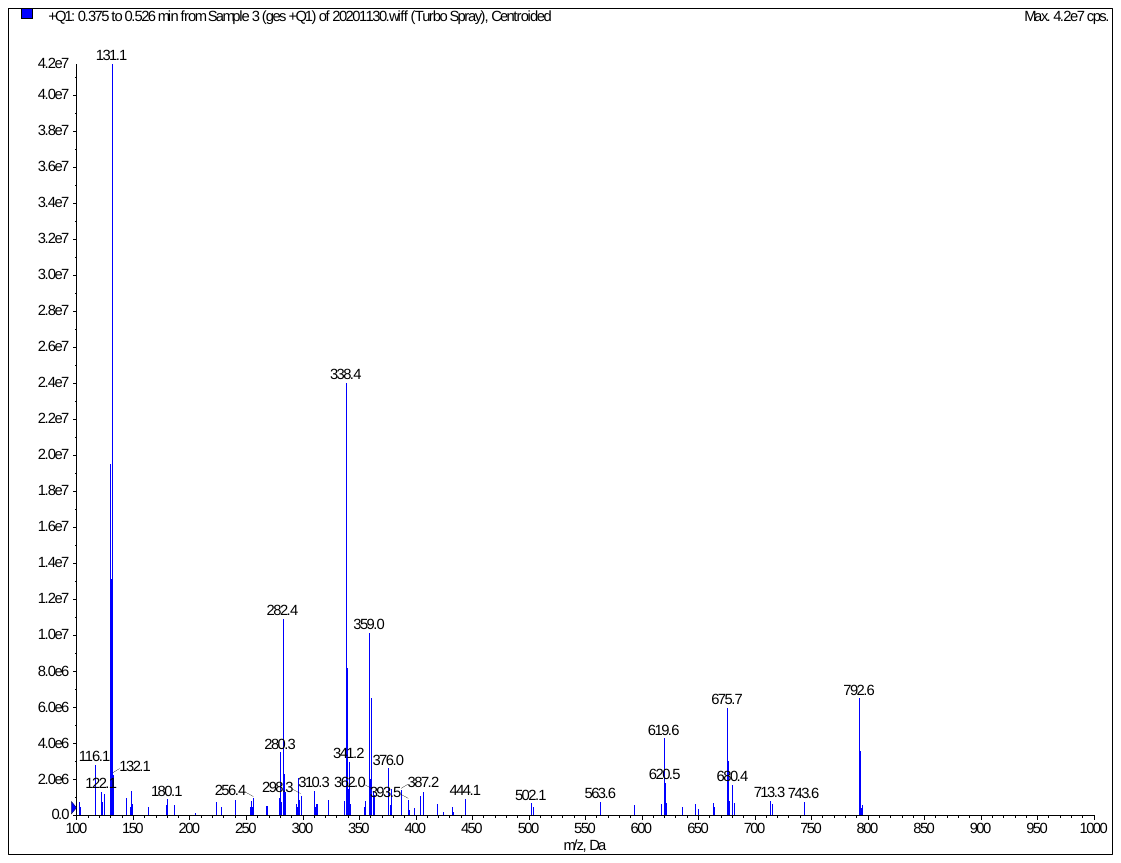

*(d). Ge-s^2^U sample* *(+ESI**). Model: second-order MS2* *(mother MS 792.6, child MS 130.1, 432.2).*

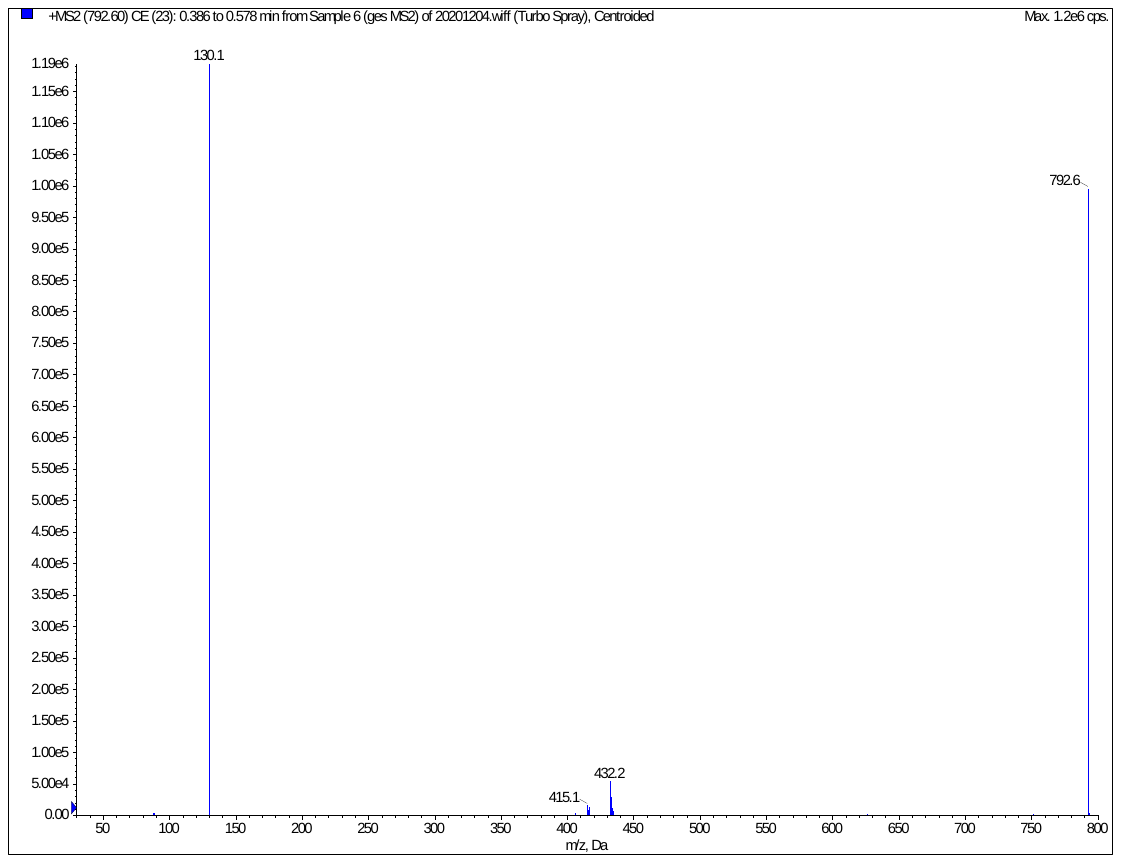

**A+H_2_O+H_2_O**

**B+18**

**A+H_2_O+1**

**2A+1**

*(e). Ge-s^2^U sample (-ESI). Model: first-order MS.*

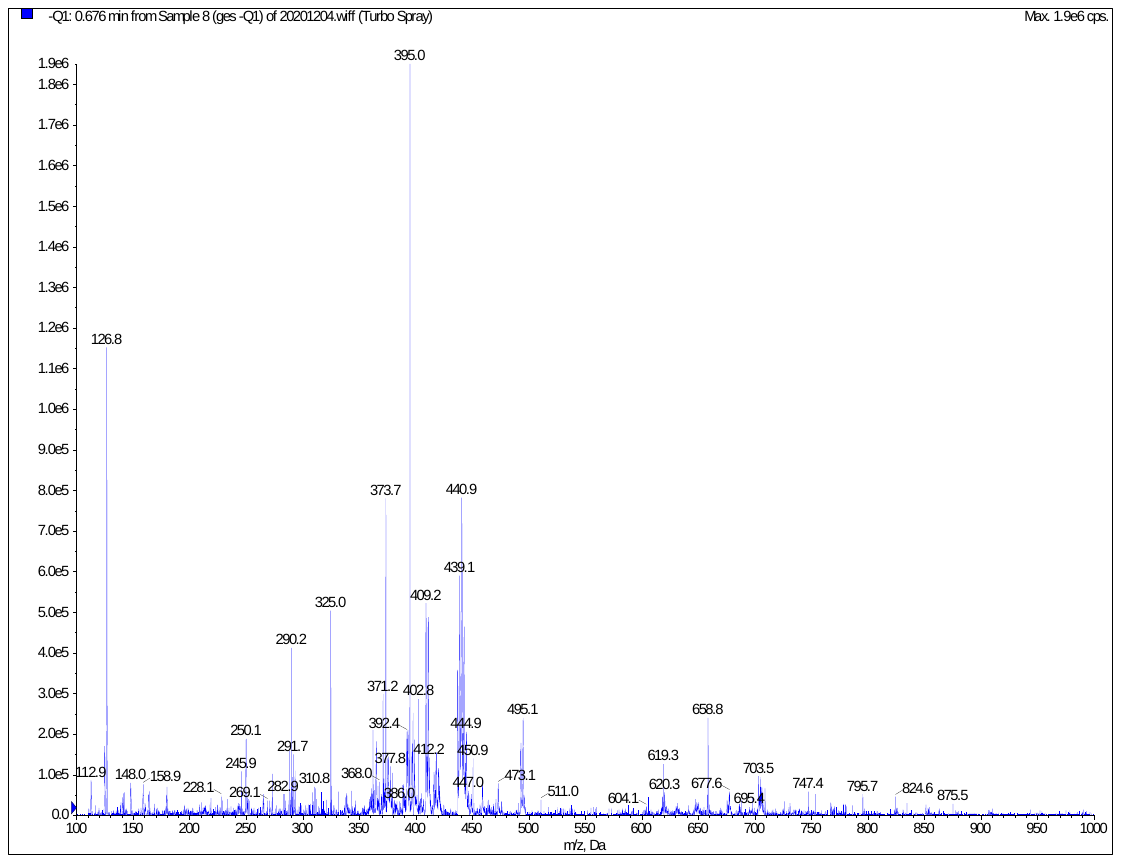

**A-1**

*(f). Ge-s^2^U sample (-ESI). Model: second-order MS2 (mother MS 395.0, child MS 148.0, 80.8).*

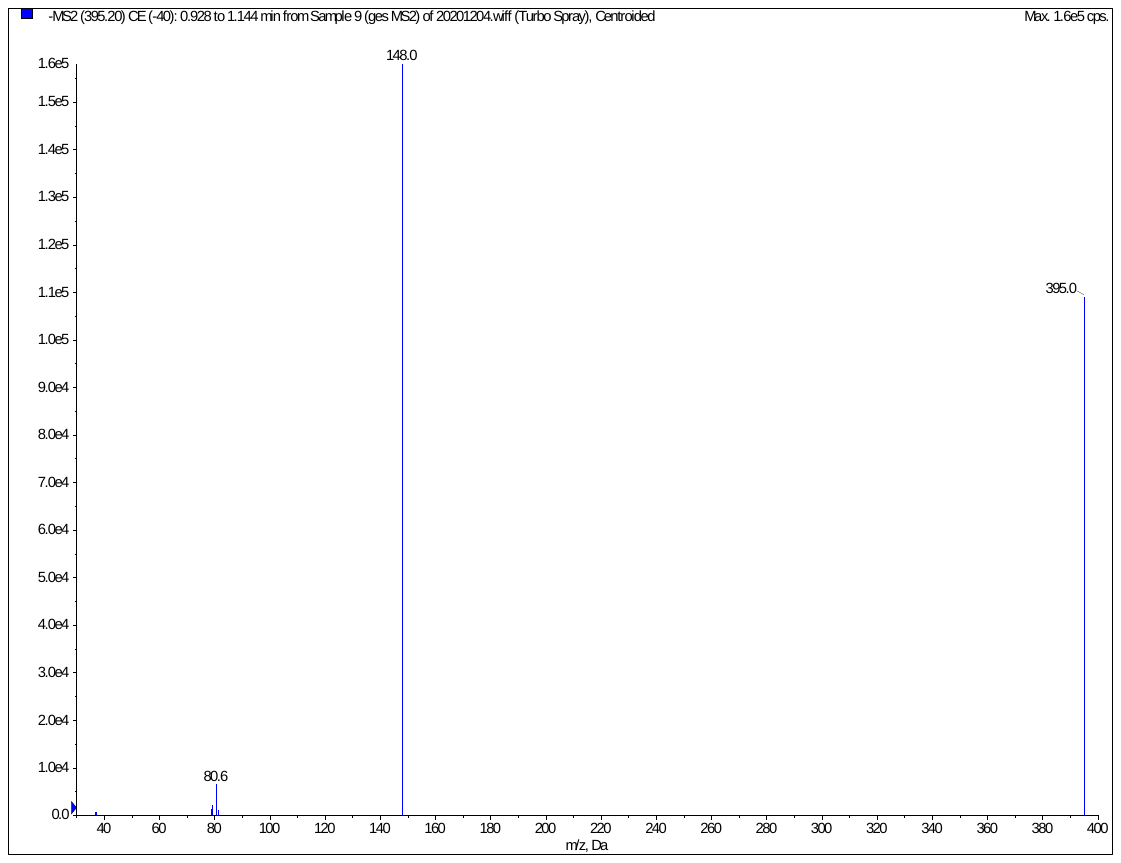

**A-1**

*(g).* *Ge-s^2^U sample LC-MS/MS chromatography (-MRM model, un-detectable).*

**Table S6**. Parameters 1.

| **Q1** | **Q3** | **DP** | **CE** |
| --- | --- | --- | --- |
| 792.6 | 130.1 | 51 | 59 |
| 792.6 | 432.1 | 12 | 11 |

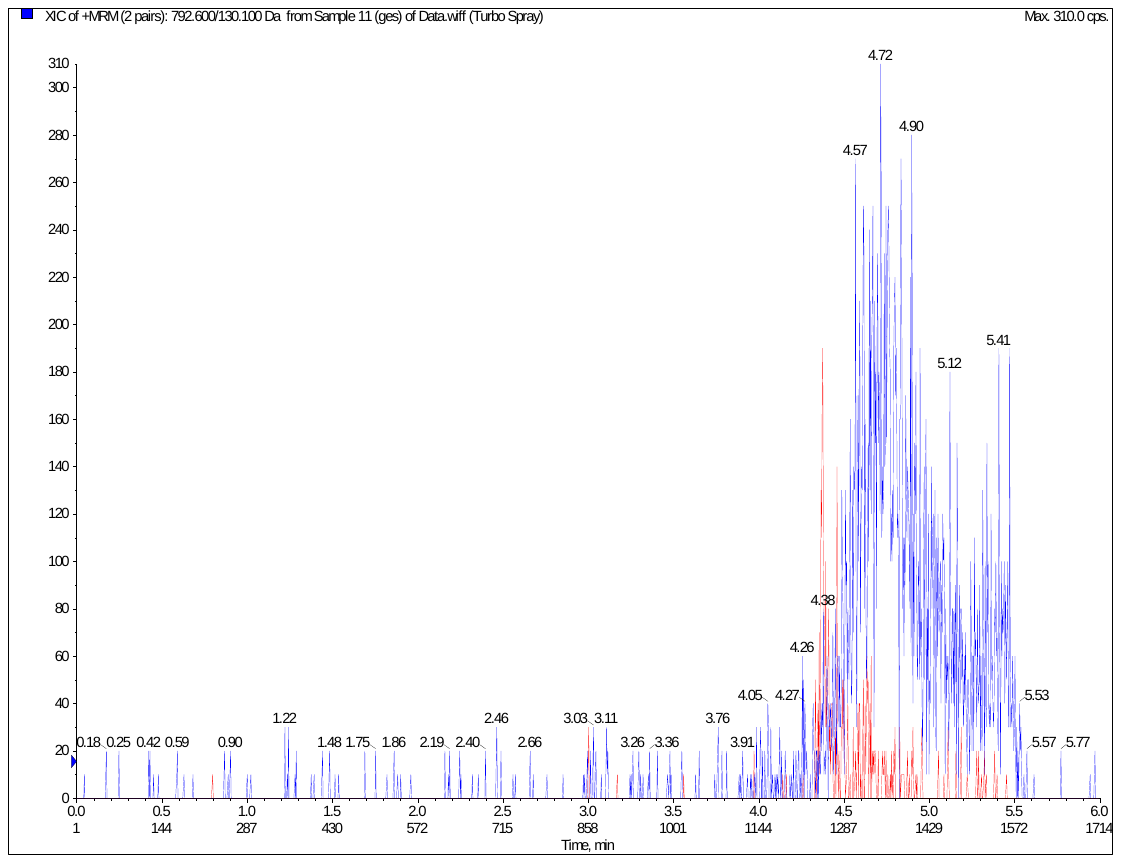

*(h)*. *Ge-s^2^U sample LC-MS/MS chromatography (-MRM model, un-detectable).*

**Table S7**. Parameters 2.

| **Q1** | **Q3** | **DP** | **CE** |
| --- | --- | --- | --- |
| 395.0 | 147.9 | -21 | -39 |
| 395.0 | 80.7 | -18 | -36 |

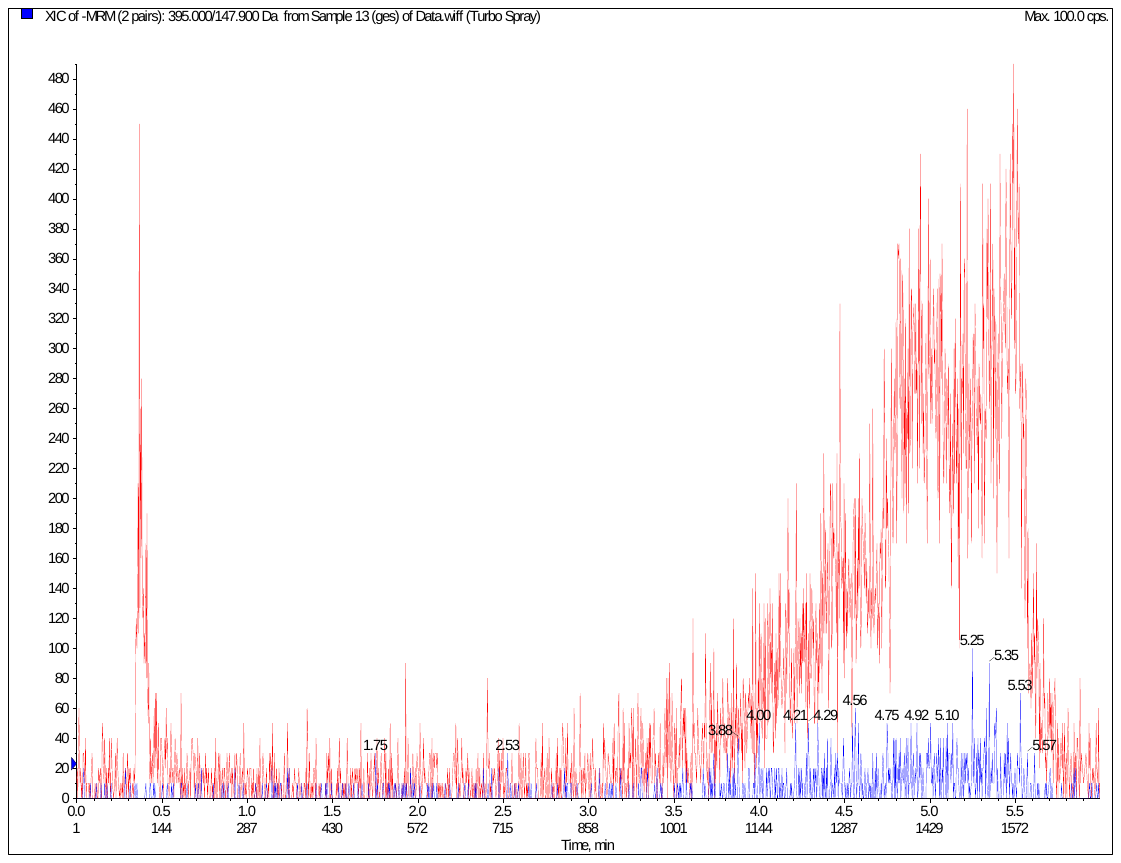

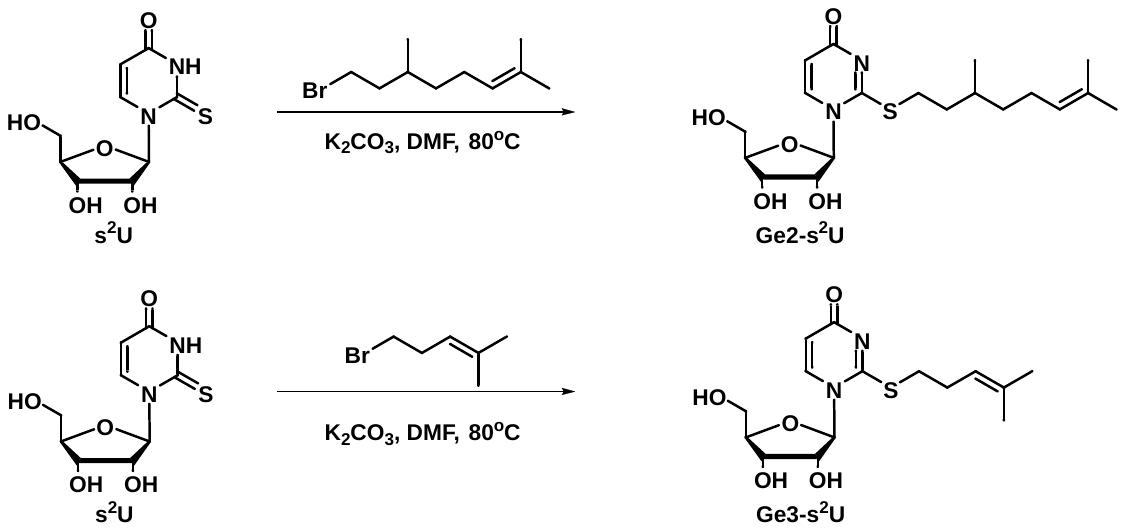

**Scheme S9**. Detailed synthetic approaches for standard nucleosides **Ge2-s^2^U** and **Ge3-s^2^U**.

LC/MS analysis. Q1: parent positive model: 397.1 MRM second: 119.0; 210.2; 323.0 (ion pair). Conditions for LC/MS: 80% CH_3_CN - 0.1% FA and 20% H_2_O - 0.1% FA, LC-4.5 min.

For the comparison of abundance of different nucleosides in various *E. coli* strains, RNA was digested with *nuclease P1* and *alkaline phosphatase* (*Lee biosciences*) to mono-nucleosides. Samples were dissolved in 50 *µ*l of 0.1% NH_4_COOH and LC was performed using a linear gradient from 0.1% aqueous formic acid (A2) to acetonitrile (B2) on an Aquity UPLC BEH C_18_ column (1.7 *µ*m, 2.1 mm Í 100 mm, Waters) at a flow rate of 0.3 mL/ min. The mobile phase composition was as follows: 100% A2 for 5 min; linear increase over 17 min to 100% B2; maintain at 100% B2 for 3 min; return to 100% A2 for 5 min.

Electrospray ionization conditions were as described above with detector was operating in positive mode. LC/MS/MS experiments for confirmation of the individual mono-nucleosides were performed at collision energies of 10, 20 and 30 eV.

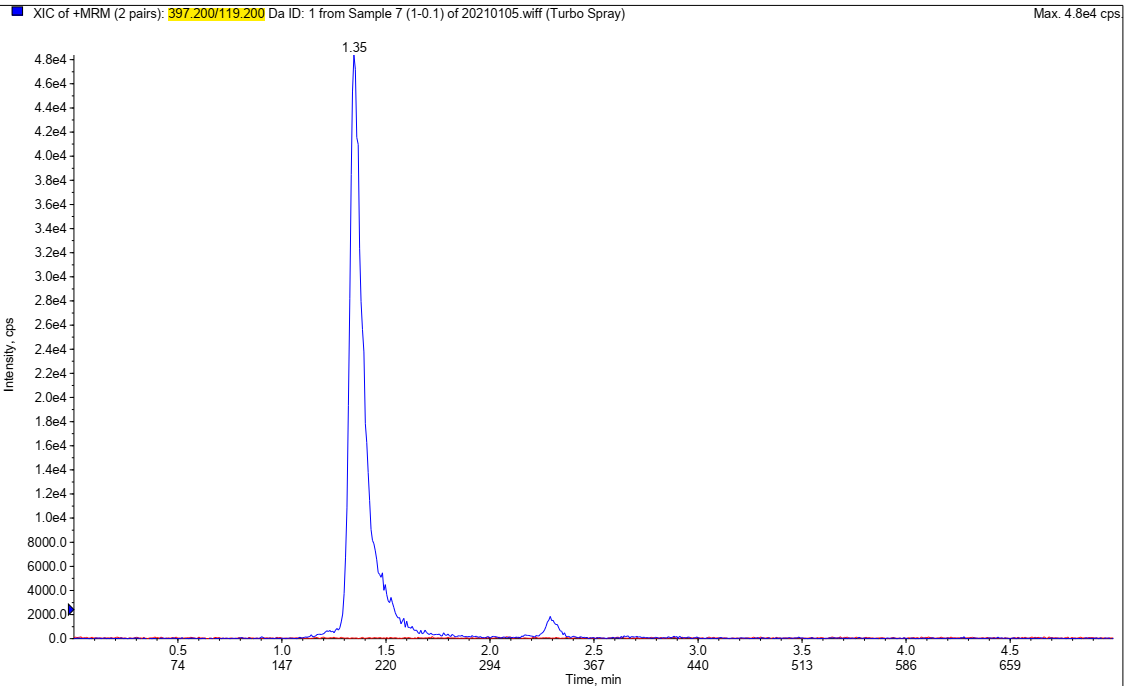

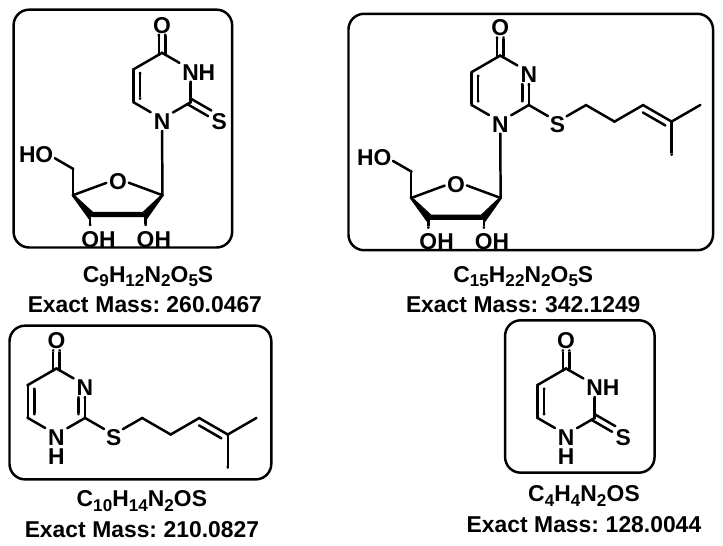

LC/MS analysis. Q1: parent positive model: 343.1 MRM second: 151.1; 211.2 (ion pair). Conditions for LC/MS: 80% CH_3_CN - 0.1% FA and 20% H_2_O - 0.1% FA, LC-4.5 min.

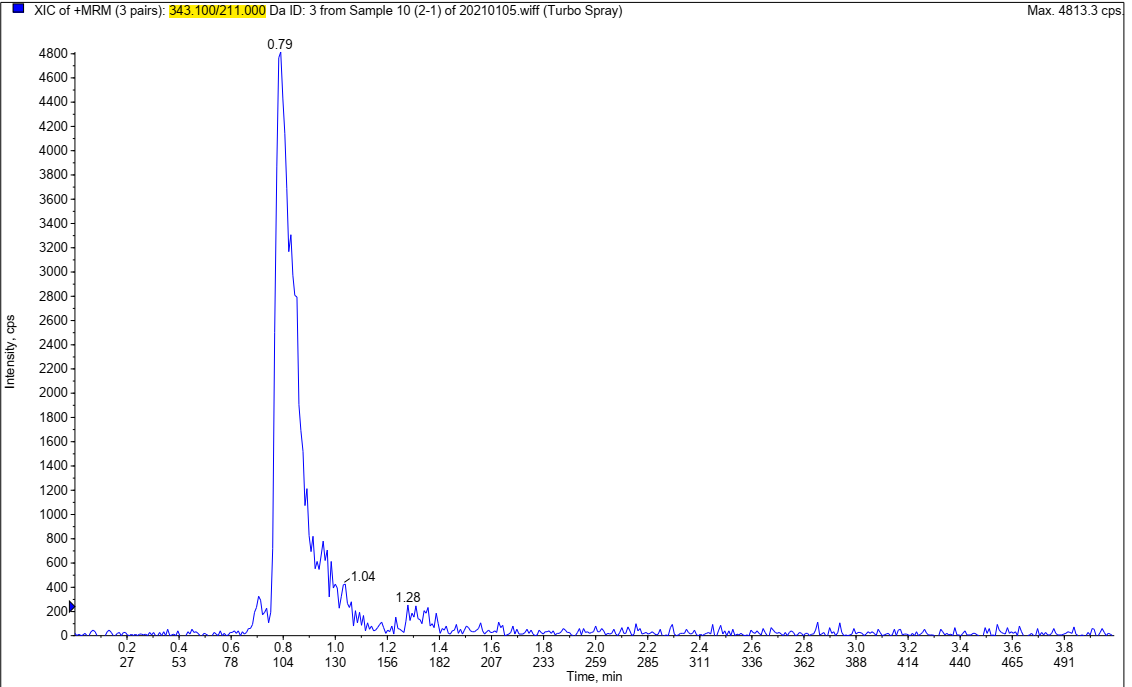

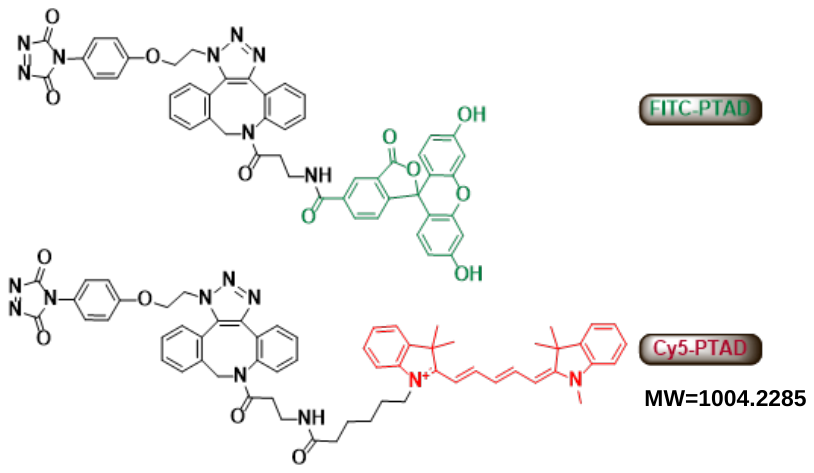

(4) Photo linking substrate (**Ge4**) for Reader Evaluation.

Co-incubation experiment of pET28b-*SelU* lysate and **Ge4** (benzophenone pyrophosphate).

Protocol:

1. Add the *E.coli* bacterial cells to the lysis solution (Western and IP lysis buffer, Beyotime), ultrasonically sonicate lysed for 5 min（80W, work 10 seconds, stop 10 seconds), lysed until the sterile body precipitates, centrifuge at 10,000*×g* for 15 min, collect the supernatant, and determine the protein concentration by BCA kit.
2. Add appropriate amount of **Ge4** (benzophenone pyrophosphate) to the protein lysate and incubate overnight at 4 °C with UV (365 nm) irradiation on ice for 15 min.
3. Take 50 *μ*l of probe from each tube of sample and add excess NBS to oxidize (the reaction solution turns pink when added), and incubate the **PTAD-DBCO-biotin** probe with the above-mentioned lysis solution for 4 h (4°C). Take 50 *μ*l beads and add them to the EP tube, place them on the magnetic stand, discard the supernatant, add buffer 2 to wash the beads twice, and add the lysate to the washed beads. Incubate overnight with rotation at 4 °C, place on a magnetic stand, washed the complex with eluent for 3 times, add 20 *μ*l protein loading buffer, centrifuge at 14,000*×g* for 30s, denaturized at 95 °C for 5 min, centrifuge at 14000*×g* 1 min, then the supernatant for SDS-PAGE electrophoresis.

SDS-PAGE electrophoresis:

**Table S8**. Preparation of SDS-PAGE gel (10%).

| **Reagents** | **Volume (mL)** |
| --- | --- |
| ddH_2_O | 1.3 |
| 30% Acr/ Bis | 1.7 |
| 1M Tris•HCl (pH 8.8) | 1.9 |
| 10% SDS | 0.05 |
| 10% AP | 0.5 |
| TEMED | 0.002 |

After mixing the above reagents thoroughly, pour the gel, then add isopropanol to press the gel, and leave it at r. t. for 30 min until gelling, then discard the isopropanol, and blot the remaining reagents with filter paper.

**Table S9**. Component of SDS-PAGE gel (5%).

| **Reagents** | **Volume (mL)** |
| --- | --- |
| ddH_2_O | 1.4 |
| 30% Acr/Bis | 0.33 |
| 1M Tris•HCl (pH 6.8) | 0.25 |
| 10% SDS | 0.02 |
| 10% AP | 0.02 |
| TEMED | 0.002 |

Mix the reagents of the gel thoroughly, pour the gel, insert a comb, and let it stand at r.t. for 30 min until the gel is completely set.

The samples of each group were loaded and electrophoresed. The stacking gel was electrophoresed at a constant voltage of 80 V for 30 min, and the separation gel was electrophoresed at 120 V for 1 h.

Stain with Coomassie Brilliant Blue staining solution on a decolorizing shaker for 30 min, and decolorize with the decolorizing solution until the protein bands can be clearly seen. The result is shown in the Figure S54 below:

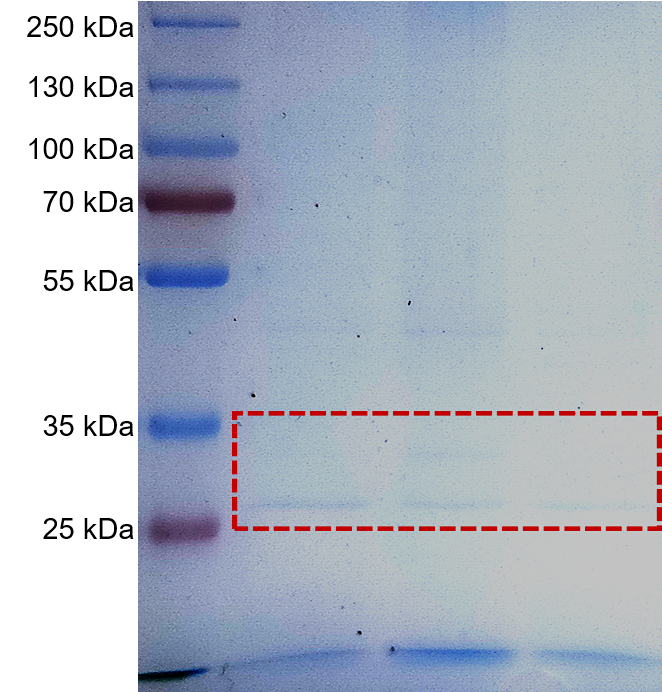

**Figure S55**. Gel image. Group1, **Ge4** was added to lysate; Group2, in the absence of **Ge4**; Group3, **Ge4** was added to lysate in the presence of *SelU*.

**DIRECT LABELING OF PRENYLATED tRNA USING THIS ENE-LIGATION**

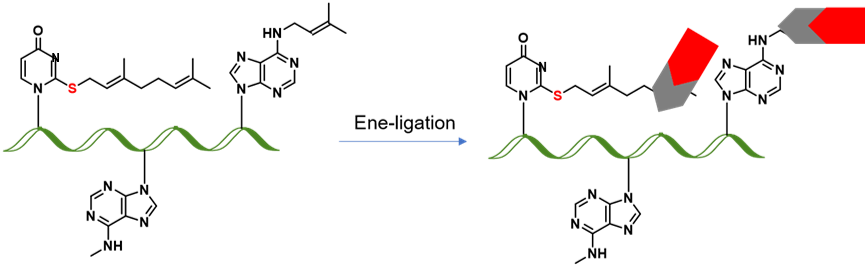

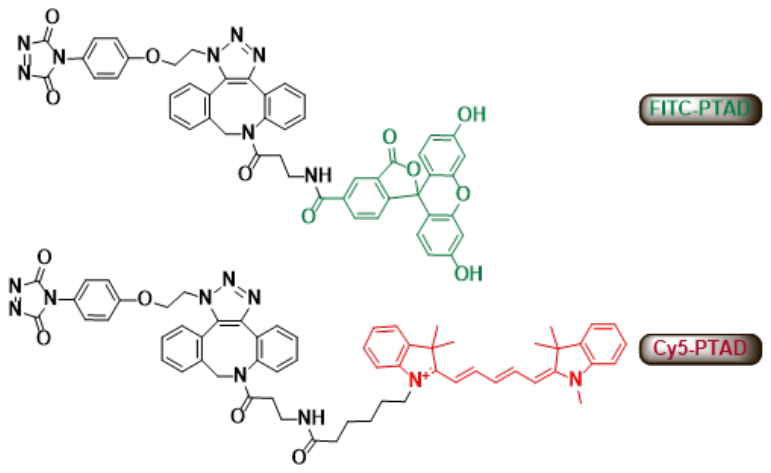

**Scheme S10**. Mass analysis of tRNA from *E. coli* with PTAD-DBCO-Cy5 *via* our well-established Ene-ligation.

tRNA ^Glu^ from *E. coli.* mass analysis: **Calculated MS 24441 (or 24124) + 77.952 + 15.023 + 181.129 (or 225.119) - 108.042 - 108.042 + 1004 = 24715.105/24579.233/24759.094/24442.094 (+1004), Found 25417.4251, 25724.55**. It is reported that geranylated RNA in *Escherichia coli*, *Enterobacter aerogenes*, *Pseudomonas aeruginosa* and *Salmonella enterica var. Typhimurium*. Ana these geranylated nucleotides occur in the first anticodon position of tRNA^Glu^_UUC_, tRNA^Lys^_UUU_, tRNA^Gln^_UUG_ at a frequency of up to 6.7% (~ 400 geranylated nucleotides per cell).

**CELL-SPECIFIC LABELING OF *SELU*-MEDIATED tRNA GERANYLATION**

**HEK293T cell culture.**

1. HEK293T cell culture and plating. washed the HEK293T cells twice with PBS, add trypsin to digest for 1 min. add 2.0 mL of DMEM medium to mix well, centrifuged at 1,000 r.p.m for 5 mins, and discarded the supernatant in the ultra-clean table. Adding fresh medium to resuspend the cell pellet and count, seeding the cells in confocal dish at the density of 0.5×10^5^, gently pipette the cells to mix, and place them in CO_2_ incubator.

2. Transfection of HEK293T cells. Prepare 150 mM sterile NaCl solution as a diluent for DNA and *Vigofect*. , replace the culture medium in the confocal dish with fresh complete culture medium before transfection, and incubate at 37 °C, 5% CO_2_. The groups were *pCMV-Myc-SelU-GFP* group, *pCMV-Myc-SelU-5s-GFP* and **Ge1** or **Ge5** or **Ge6** group. Configure the transfection working solution:

a Take 2.5 *μ*g pCMV-Myc-SelU-GFP or pCMV-Myc-SelU-5s-GFP, add the diluent to 100 *μ*l and store at room temperature 5 min.

b Add 2.5 *μ*l VigoFect to the diluent to 100 *μ*l, mix gently and place at room temperature for 5 min.

c Add the diluted *VigoFect* to the diluted DNA solution gently, and place the resulting transfection working solution at room temperature for 15 minutes.

d Add working solution to the culture solution, mix gently and incubated for 24 hours.

3. After 24 hours of transfection, the **Ge1、Ge5、Ge6** (1 mM) were added to the medium for 4 hours. Cells were washed with PBS for 3 times, fixed with 4% paraformaldehyde for 15 min, and subsequently permeabilized with 0.5% Triton X-100 for 15 min. The TMSN_3_ and Selectfluor (50 *μ*M) were added to **Ge1** group for 30 min, Cells were then incubated with DBCO-Cy5 for 30 min before stained with 4′, 6-diamidino-2-phenylindole (DAPI) for 15 min in darkness at room temperature (RT). Samples were washed with PBS for three times and examined under a confocal microscope (LSM780，Carl Zeiss).

**Amplification, expression, and extraction of *pCMV-Myc-SelU-GFP*, *pCMV-Myc-SelU-5s-GFP* plasmid.**

1. *pCMV-Myc-SelU-GFP* or *pCMV-Myc-SelU-5s-GFP* transformation. Take out the competent cell DH5α from -80 °C and place it on ice. Add 1 *μ*l plasmid to competent cells, mix gently and bath in ice for 30 min. Heat shock at 42 °C for 90s, and quickly place on ice for 5 min. Add 400 *μ*l of LB solution, shake at 37 °C, 220 r.pm./min for 45 min, spread the bacterial solution onto the Kana-containing LB solid culture plate. Place it in 37 ℃ incubator, 12-16 h later until a single colony appears.

2. After 16 hours，Pick a single clone and amplify it. In the ultra-clean table, pick a single colony from the plate into 5 mL of LB solution (kana), incubate overnight at 37 °C, 220 r.p.m./min, until the culture solution becomes turbid. Perform plasmid extraction.

3. Plasmid extraction adopts Dynake Biological Plasmid Extraction Kit

① Add 250 *μ*l of Buffer BL to the adsorption column AC and centrifuge at 12,000🞨*g* for 1 min to activate the silica gel membrane.

② Take 4 ml of the bacterial solution cultured overnight, centrifuge at 12,000🞨*g* for 1 min, and collect the bacterial cells.

③ Add 200 *μ*l Buffer S1 to resuspend the bacterial pellet, vortex and shake until a sterile block is reached.

④ Add 200 *μ*l Buffer S2, turn up and down and mix 7 times to fully lyse the bacteria.

⑤ Add 200 *μ*l Buffer S3, and mix up and down 7 times. Currently, the solution appears white flocculent precipitate. Centrifuge at 12,000🞨*g* for 15 min.

⑥ Aspirate the supernatant carefully, transfer the supernatant to the adsorption column AC, centrifuge at 12000🞨*g* for 1 min, discard the waste liquid, and put the adsorption column AC back into the empty collection tube

⑦ Add 700 *μ*l Buffer W2 to the adsorption column AC, centrifuge at 12,000🞨*g* for 1 min, and discard the waste liquid. Repeat this step.

⑧ Put the adsorption column AC back into the empty collection tube and centrifuge at 120,00 🞨*g* for 2 min.

⑨ Take out the adsorption column AC, put it in a clean 1.5 mL centrifuge tube, let it stand at 25 ℃ for 2 min, add 30 *μ*l of Eluent to the middle of the adsorption membrane, let it stand at 25 ℃ for 2 min, and centrifuge at 12,000🞨*g* for 2 min. Obtain the plasmid, and determine the concentration.

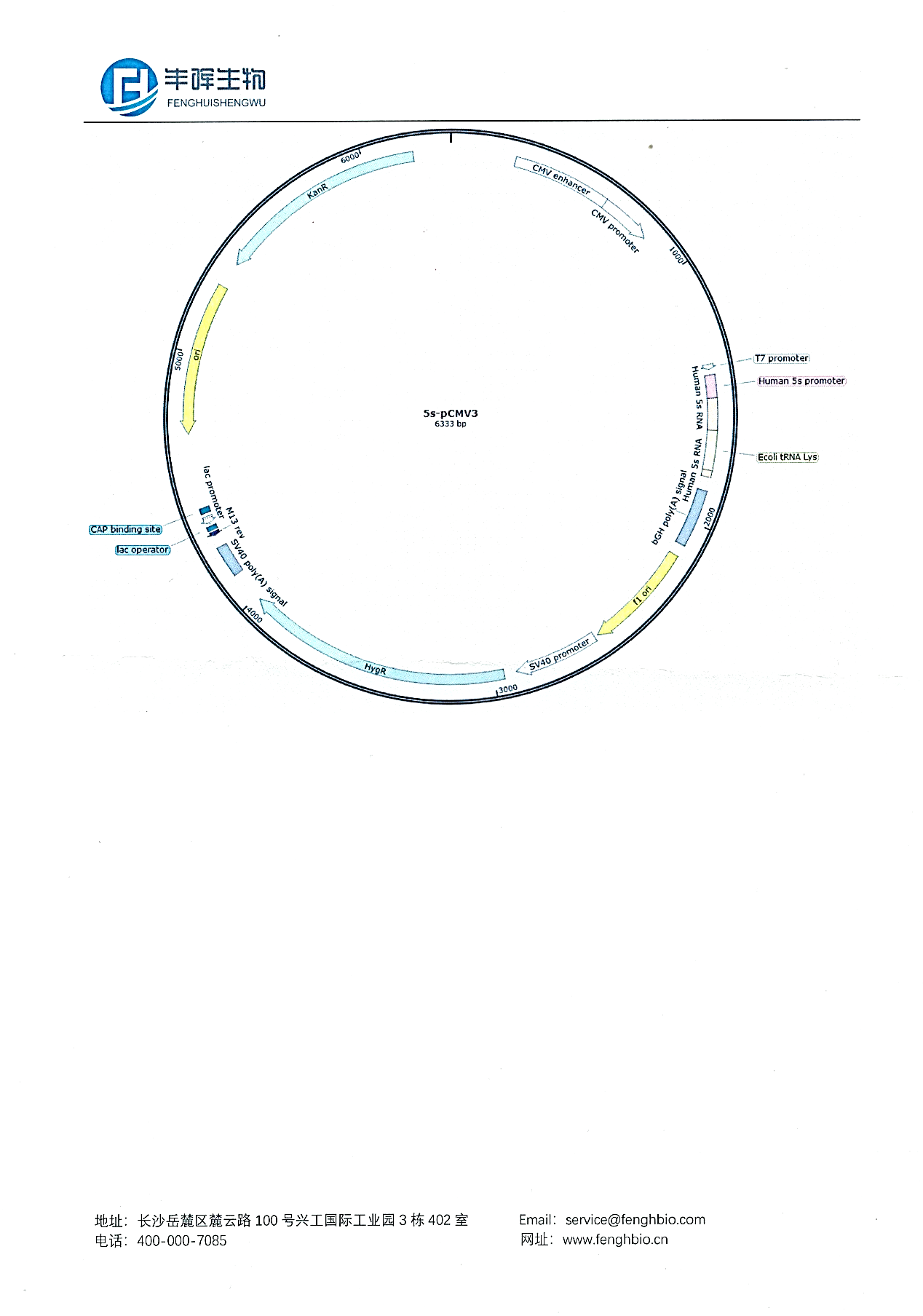

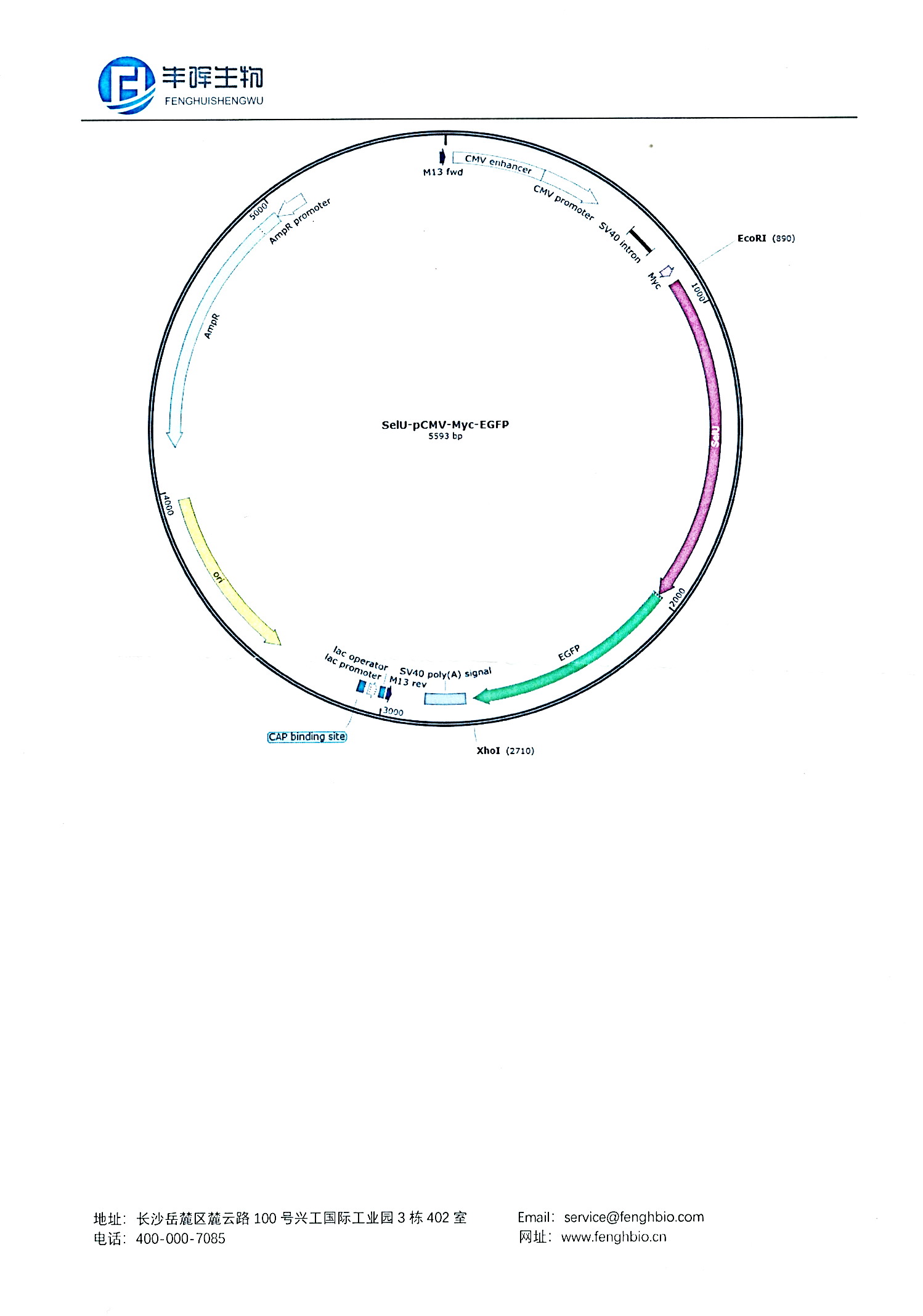

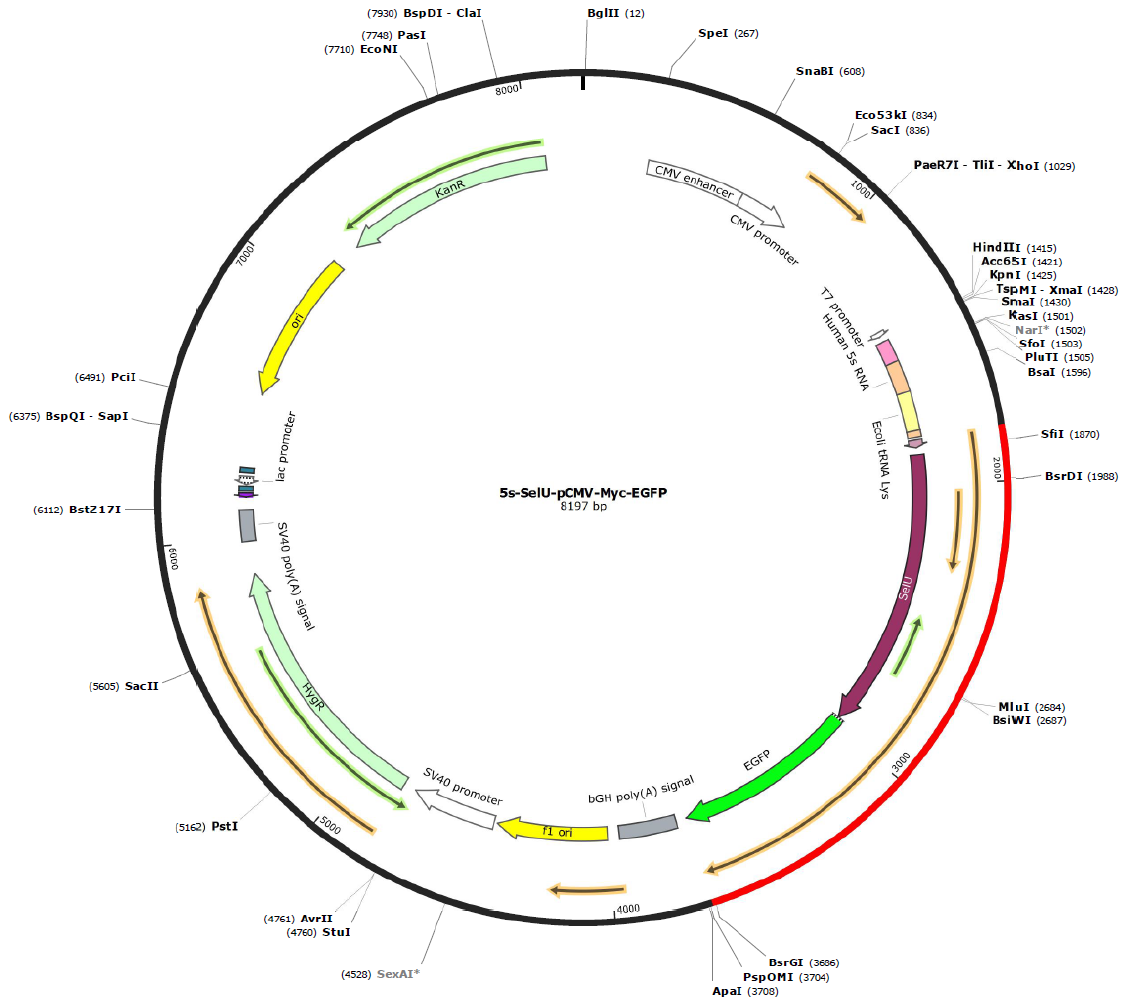

**Figure S56.** Construction map for *5s-pCMV3* and *SelU-pCMV-Myc-EGFP*. These plasmids were construction for the studies of expressed SelU enzyme as well as tRNA^Lys^ from *E.coli.*

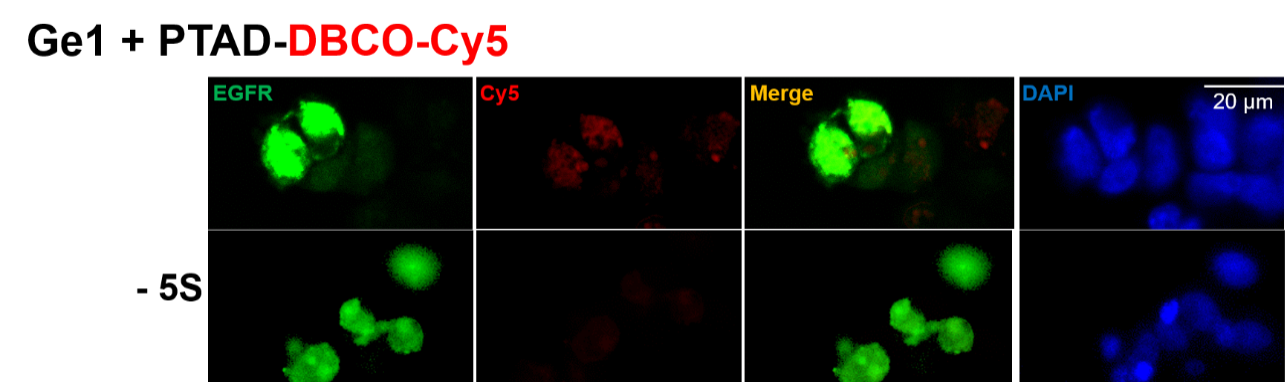

**Figure S57.** Confocal imaging investigations of fluorescent labeling of tRNA harboring *5s-pCMV3* and *SelU-pCMV-Myc-EGFP* in HEK293T cell lines using probe PTAD-DBCO-Cy5 in the presence of pyrophosphate **Ge1**, or probe DBCO-Cy5 in the presence of **Ge5** or **Ge6**. Control assays are: in the absence of *SelU-pCMV-Myc-EGFP,* in the absence of *5s-pCMV3.* Scale bar: 20 µm.

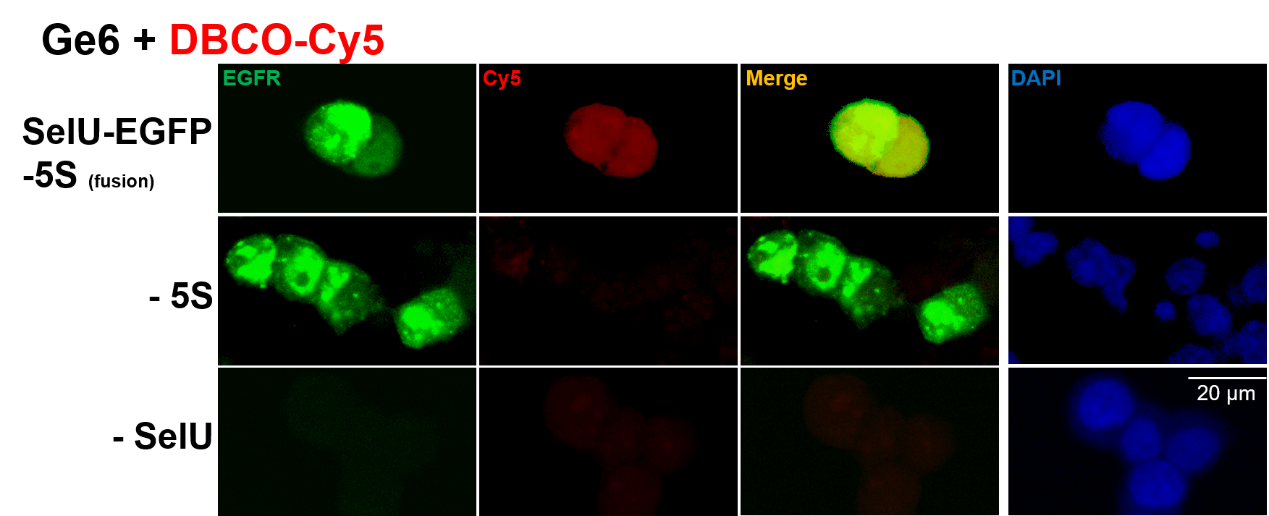

**Figure S58.** Confocal imaging investigations of fluorescent labeling of tRNA harboring *SelU-pCMV-Myc-EGFP* and *SelU-pCMV-Myc-EGFP* in HEK293T cell lines using probe DBCO-Cy5 in the presence of **Ge6**. Scale bar: 20 µm.

**INSTALLATION OF PHOTO LINKING PROBE IN THE tRNA GERANYLATION FOR THE SRUDIES OF POSSIBLE READER PROTEINS**

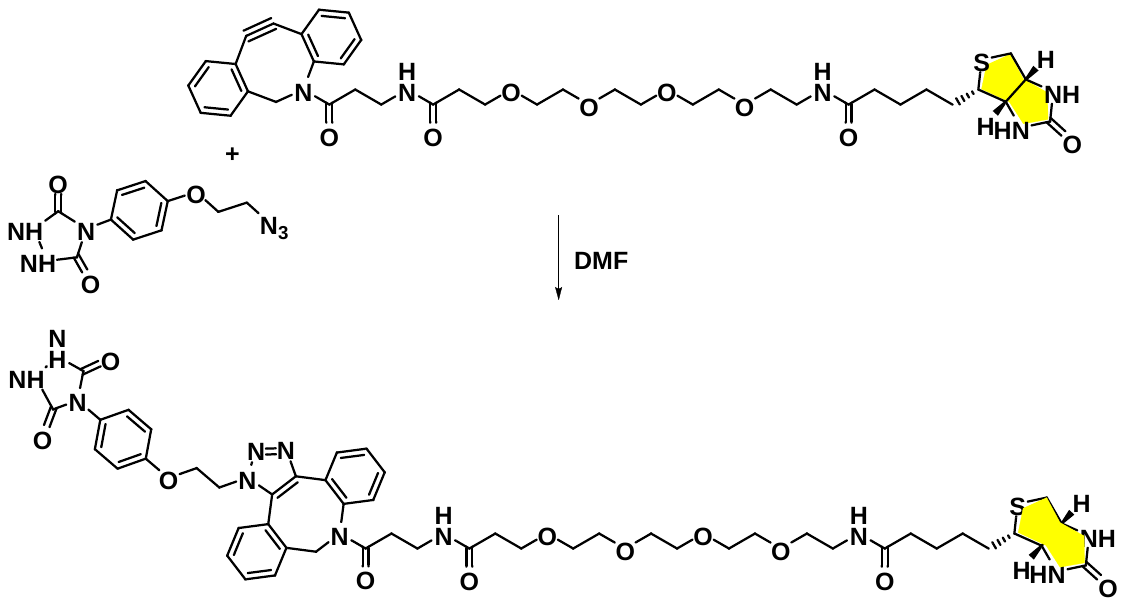

**Scheme S11.** Overall illustration of two-head probe (**5c**).

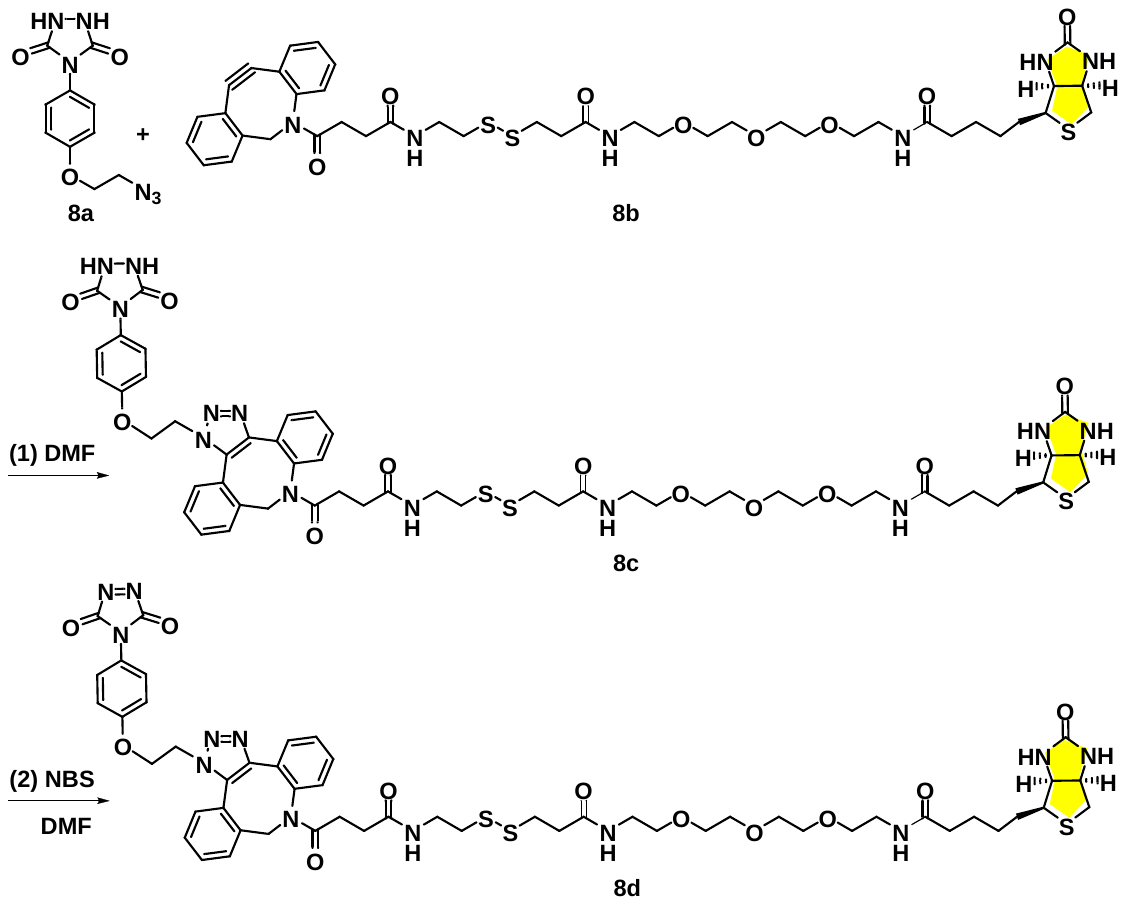

**Scheme S12.** Overall illustration of two-head probe (**8d**).

Procedure: A mixture of **8b** (5.1 mg, 0.006 mmol), **8a** (3.2 mg, 0.012 mmol) in anhydrous DMF (0.2 mL) under N_2_ atmosphere was stirred overnight at r. t. The reaction mixture was purified by HPLC.

**Three-head probe design and synthesis and its use for trapping tRNA geranylation reader proteins.**

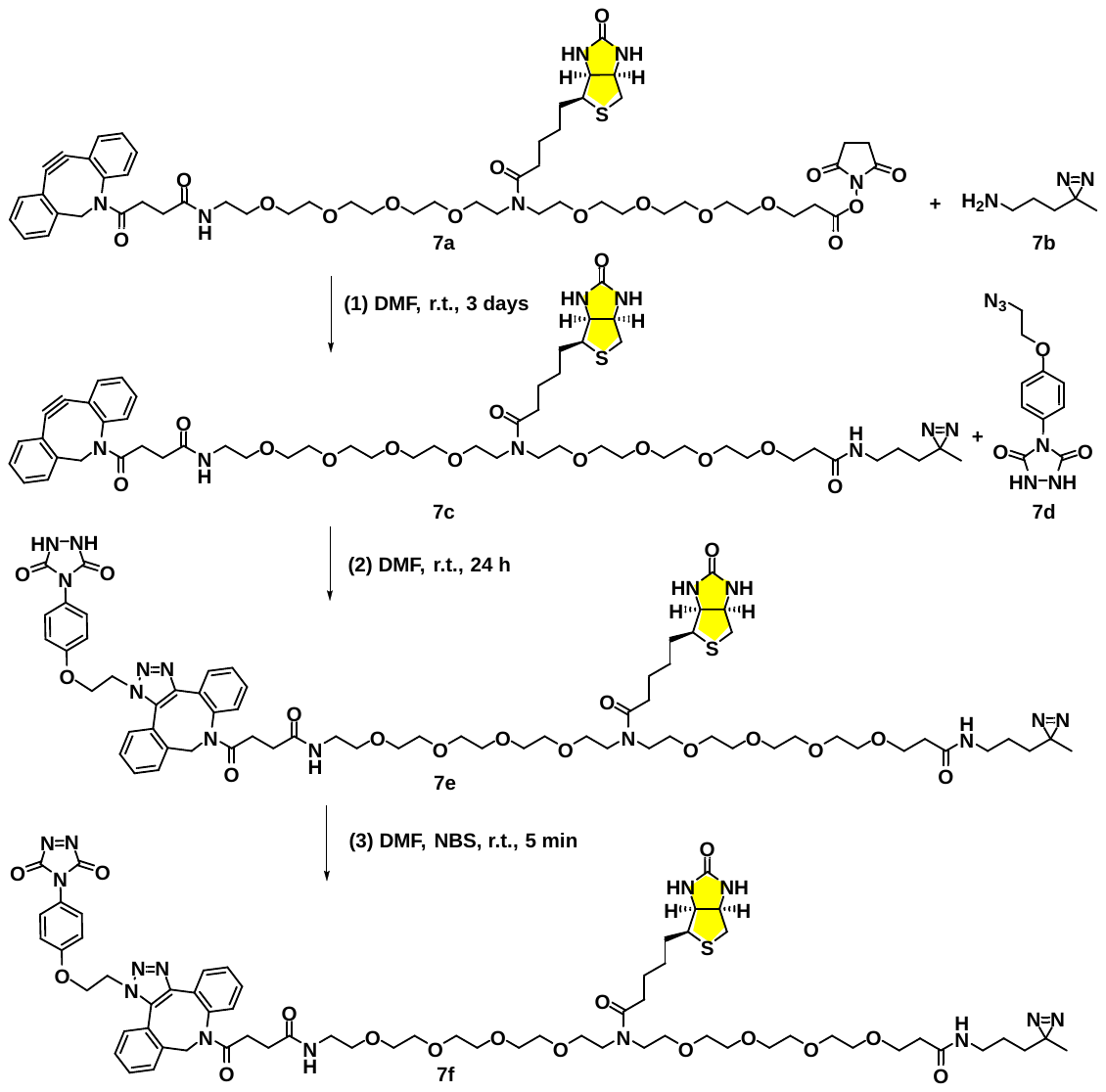

**Scheme S13**. Overall illustration of synthetic route for three-head probe (**7f**).

General procedure: To a stirred solution of **7a** (1 mg, 0.001 mmol) in dry DMF (0.2 mL) was added **7b** (1 mg, 0.001 mmol) under N_2._ This mixture was stirred at r.t. overnight. 3 days later PTAD-N_3_ (**7d**, 2 mg, 0.007 mmol) was added (in DMF solution). The mixture was stirred at r.t. overnight. Compound **7e** was obtained through HPLC purification. [M+H]^+^=1355.6458. Probe **7f** was generated from **7e** which was formed by in-situ oxidation with NBS in DMF for 5 min.

**NMR and Mass Spectra III of the Probe**

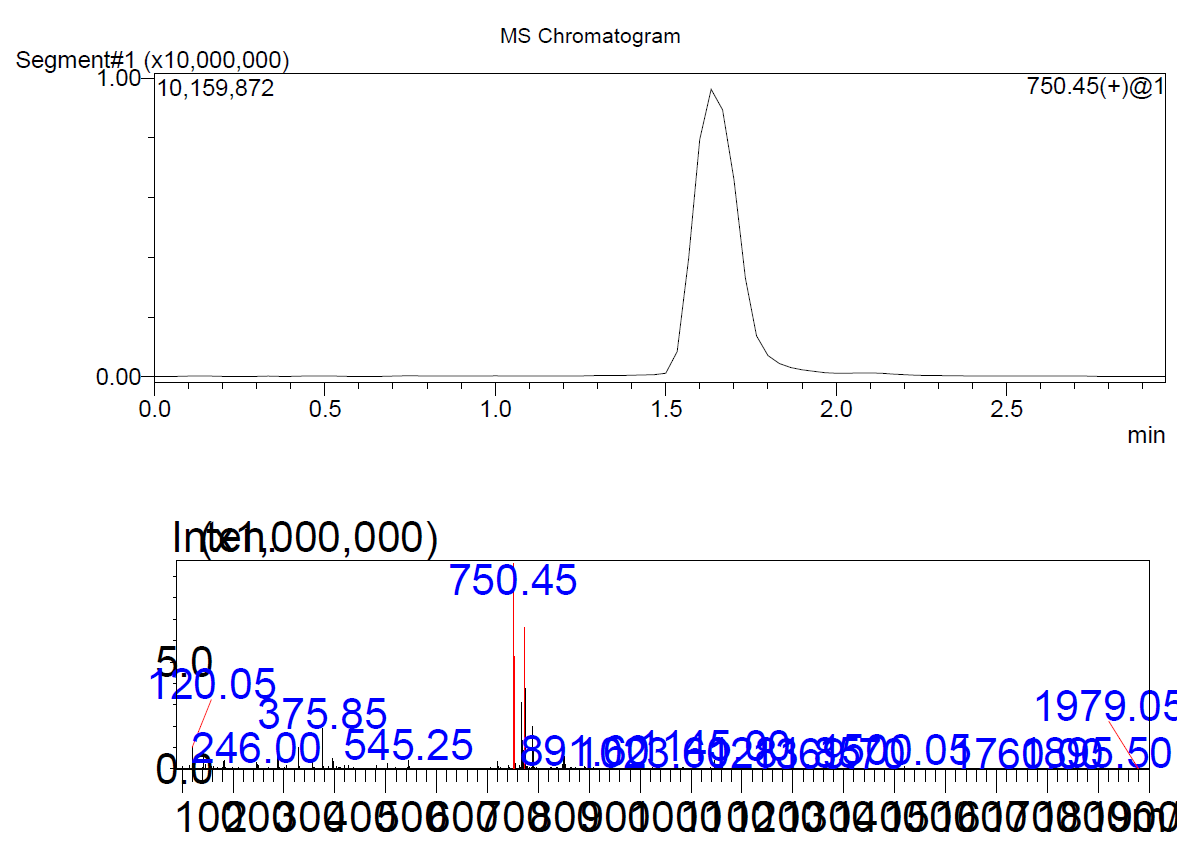

**Figure S59. HRMS** of compound (**6c**).

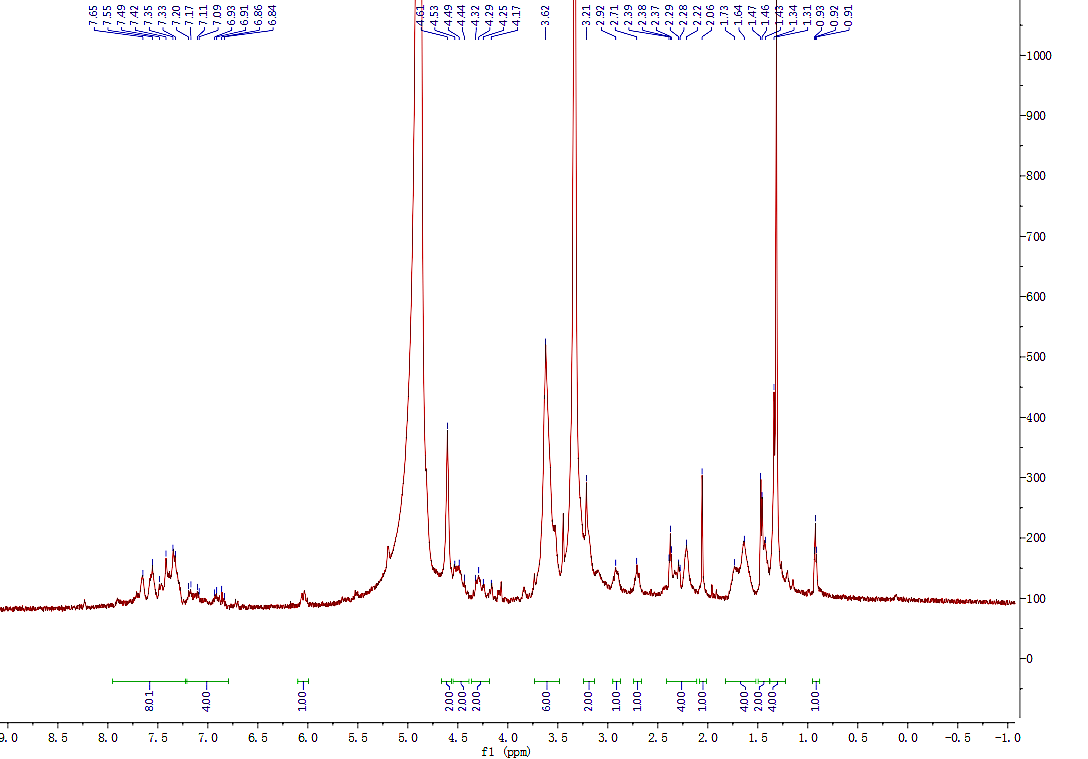

**Figure S60. ^1^H NMR** of compound (**6c**).

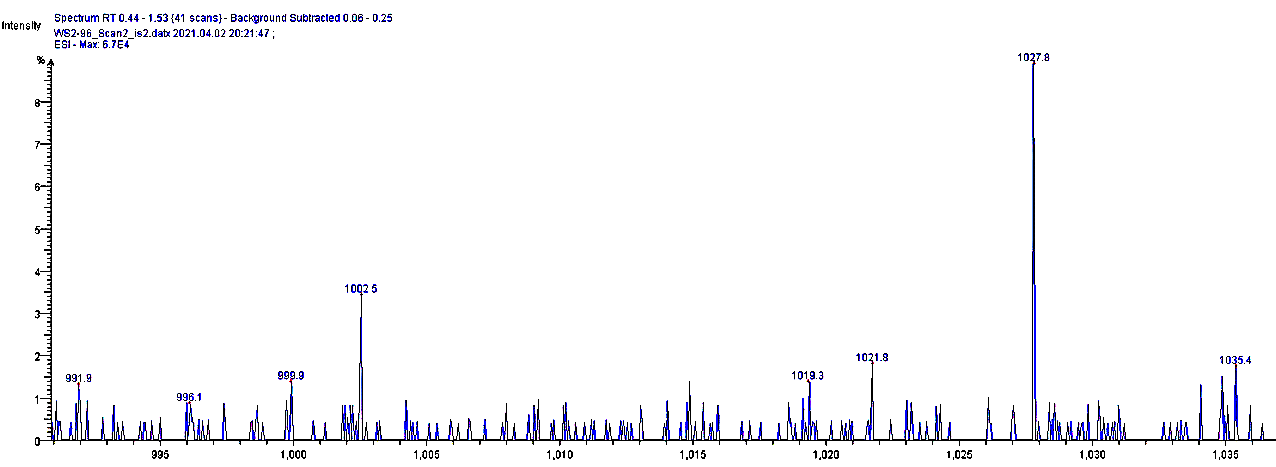

**Figure S61**. **MS** spectrum of compound (**6c**). MS(ESI): Calculated [M+H] ^+^ = 1011.4273 (C_49_H_62_N_11_O_11_S), Found [M+H] ^+^ = 1027.8.

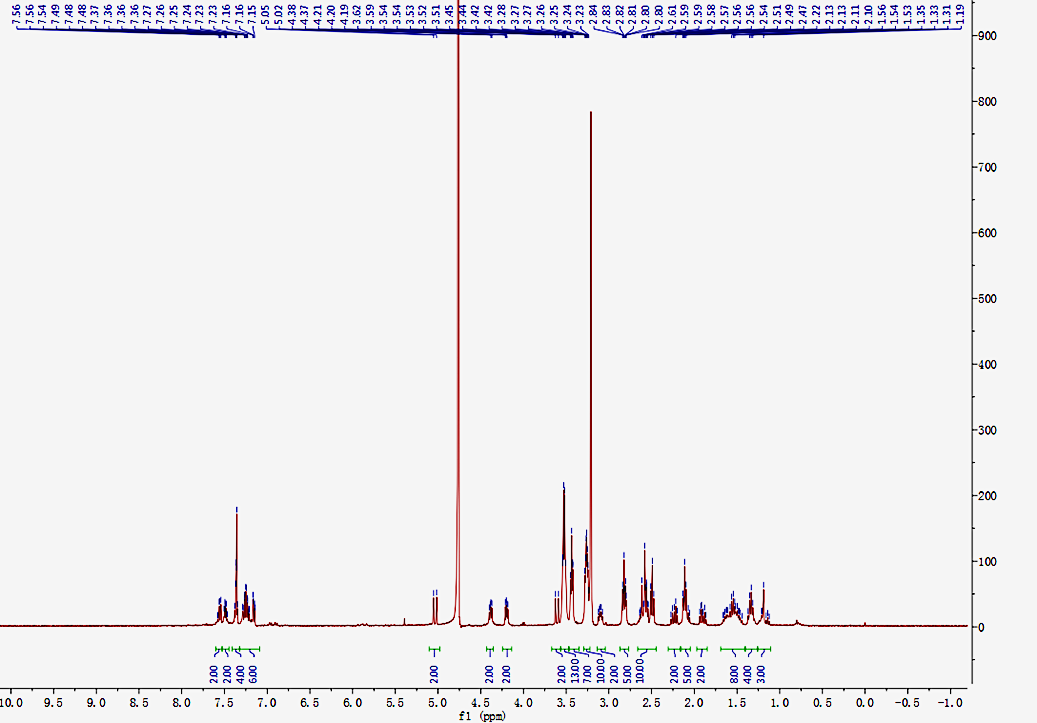

**Figure S62. ^1^H NMR** of biotin (**8d**).

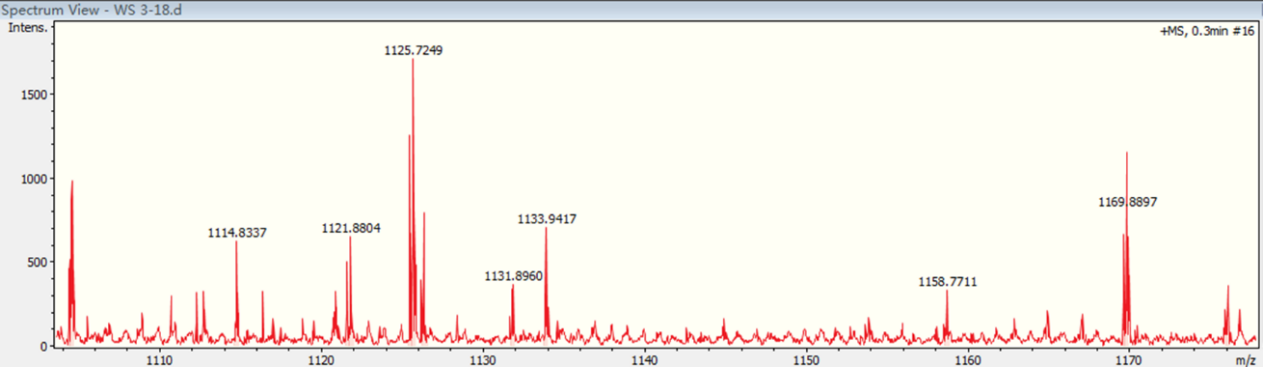

**Figure S63**. **HRMS** of biotin (**8d**). MS(ESI): Calculated [M+H] ^+^ = 1131.4144 (C_52_H_67_N_12_O_11_S_3_), Found [M+H] ^+^ = 1131.8960.

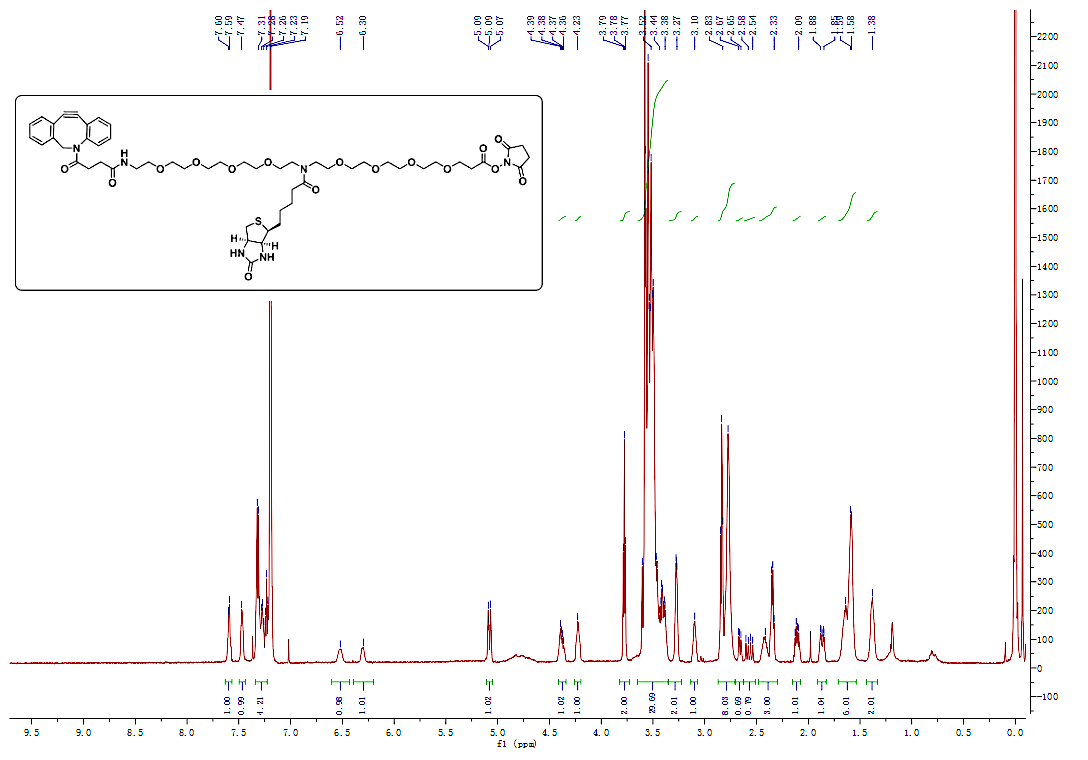

**Figure S64**. **^1^H NMR** of compound (**7f**).

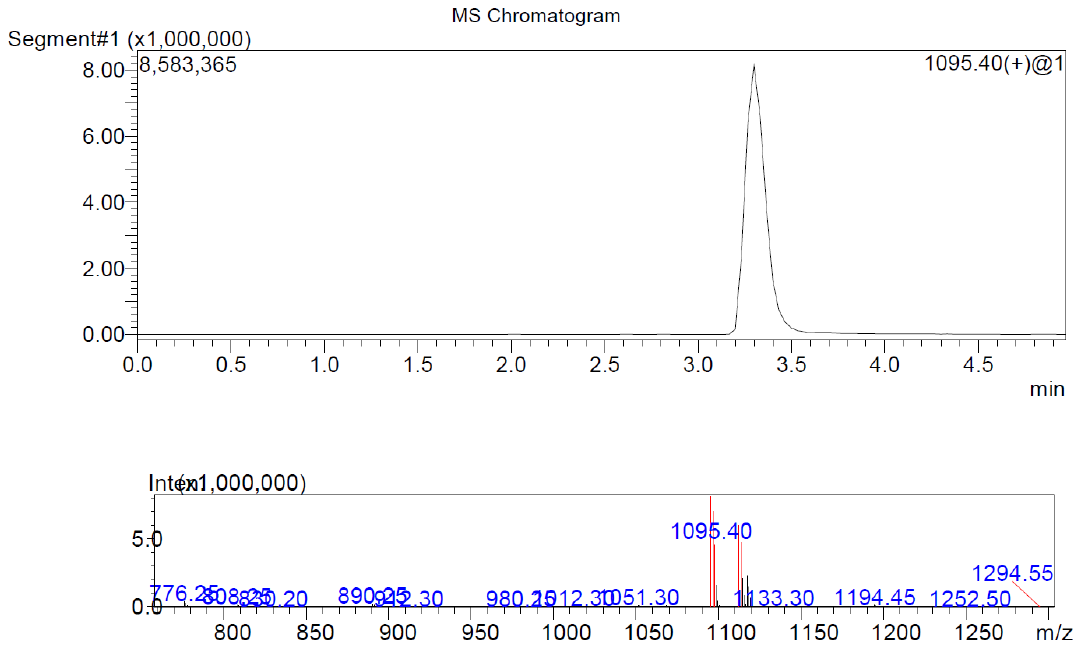

**Figure S65. MS** result of compound (**7f)**. Target MS [M+H] ^+^ = 1095.40.

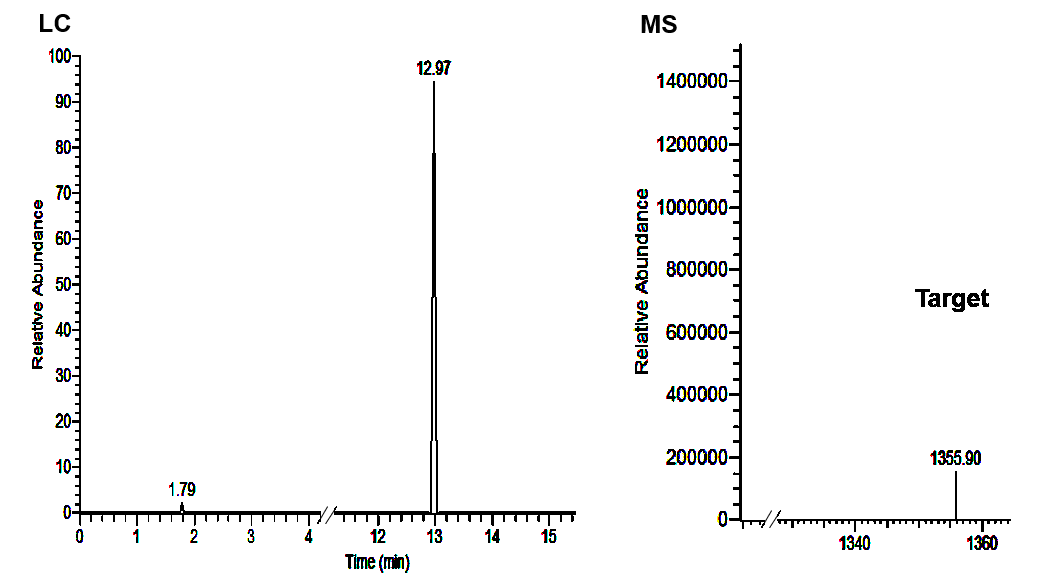

**Figure S66.** MS result analysis of target probe (**7f**) using LC-MS chromatography. Target MS [M+H] ^+^ = 1355.90.

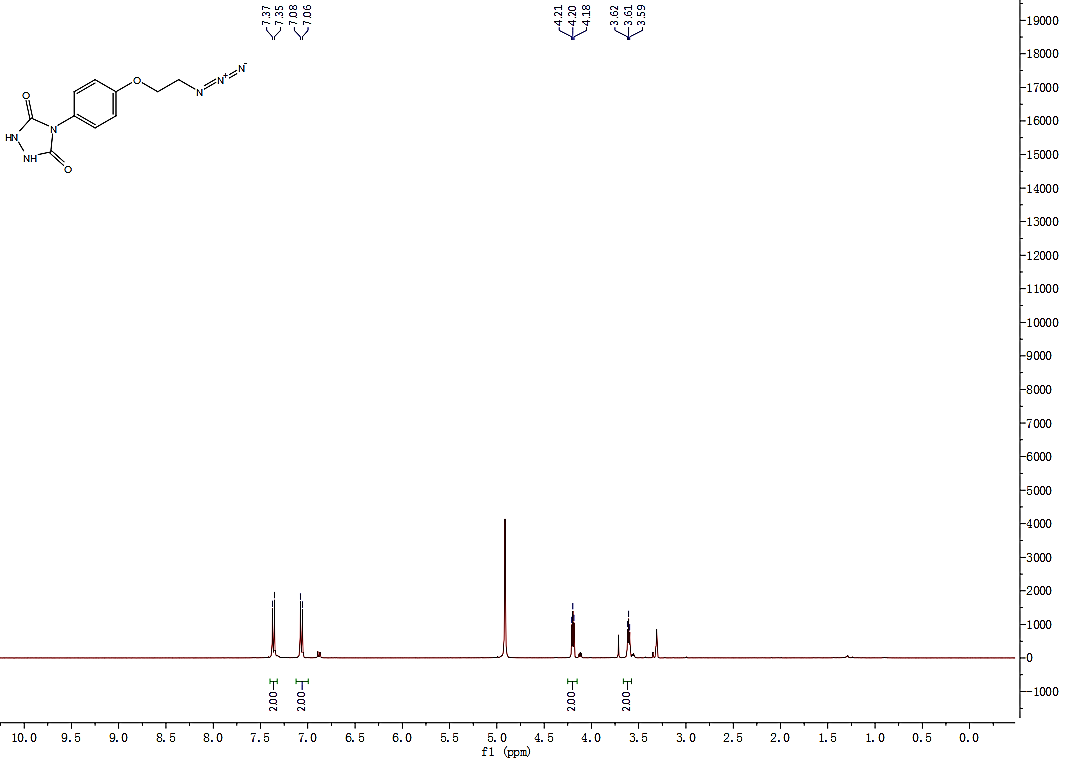

**Figure S67. ^1^H NMR** of azido intermediate.

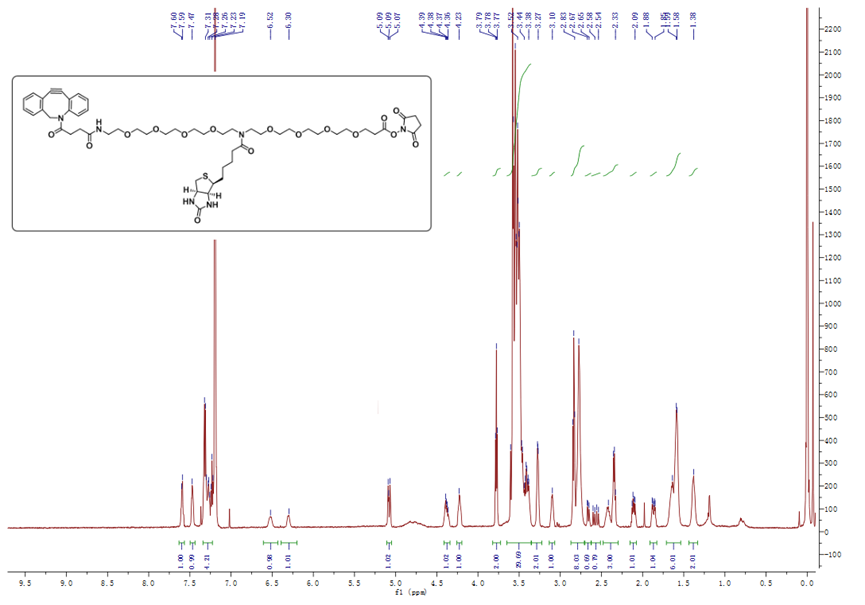

**Figure S68.** **^1^H NMR** of DBCO intermediate.

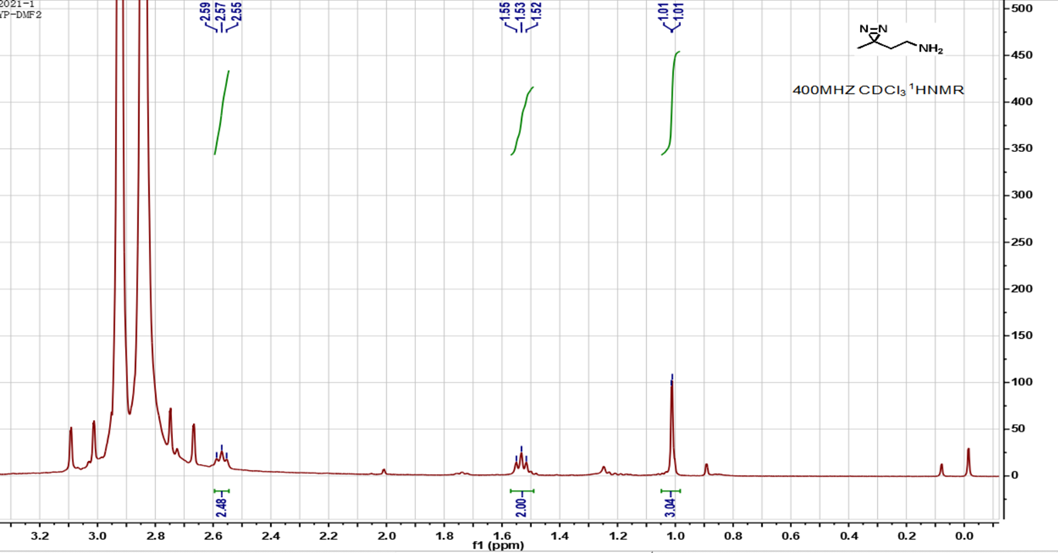

**Figure S69. ^1^H NMR** of amine intermediate.

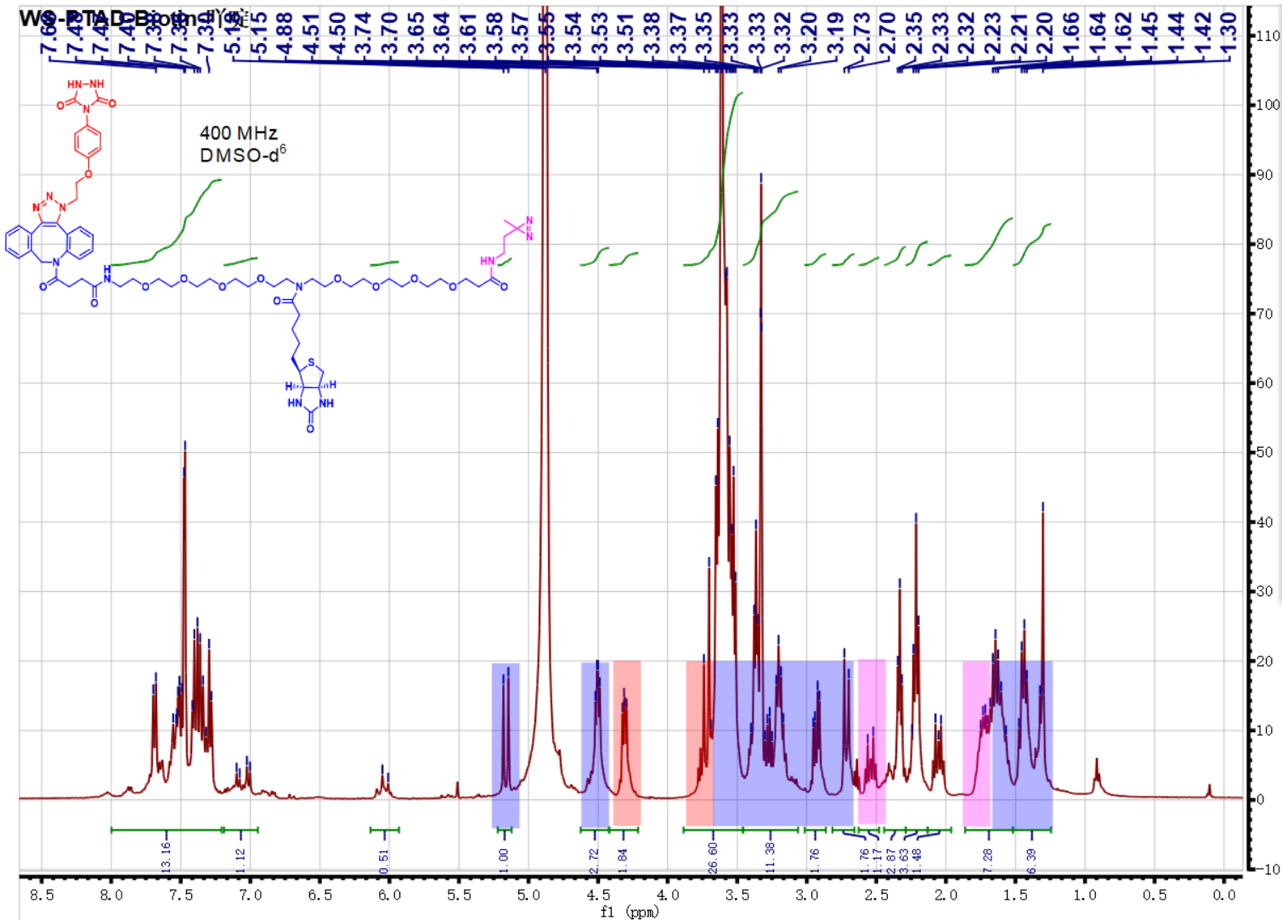

**Figure S70. ^1^H NMR** of three-head probe (**7g**).

**General procedure of geranylated tRNA and interacted proteins evaluations.**

(1) The target probe **7g** precursor (2.0 mM) is dissolved in DMF and NBS (200.0 mM) is added for 5 min at r.t. Then, probe **7g** (100 *μ*L) was added into 1.0 mL of protein lysate (200 *μ*M) and was incubated for further 5 min. Then, the mixture was irradiated with UV for 15 min, and was incubated at 4 °C overnight.
(2) The binding beads was washed twice with PBST, then was placed on a magnetic stand to remove the supernatant.
(3) The probe-containing protein lysate was added into Beads and was incubated at 4°C overnight with slow rotation.
(4) The EP tube was placed on a magnetic stand to elute unbound proteins, subsequently protein loading buffer was added, and was centrifuged at 14,000 ×*g* for 30 s. Then the mixture was heated at 95 °C for 5 min, and was centrifuged again. Eventually, the supernatant was analyzed and detected by SDS-PAGE electrophoresis.

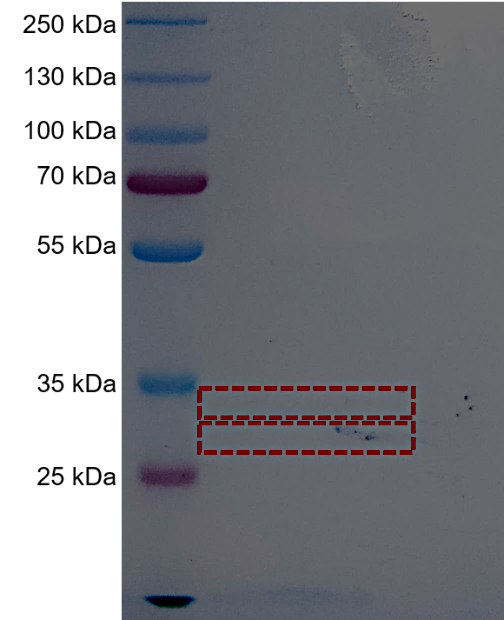

**Figure S71**. First experiment result. Group 1, probe **7g** generated *in-situ* was added in to *E. coli* protein lysis solution.

Optimized method:

In more details,

(1) First, 500 *μ*g *E. coli* tRNA (Zymo purification kit or from Roche) was incubated with 20 *μ*l purified *SelU* protein (1 mM), 100 *μ*l geranyl pyrophosphate (i.e., **Ge1,** 1 mM), and the probe (i.e., **7g**, 1 mM) in 100 *μ*l buffer (10 mM Tricine-NaOH, pH 8.0, 0.2 mM DTT, and 100 mM MgCl_2_, 2% glycerol at 25 ^o^C) for 24 h. Then, the mixture was irradiated with UV light (365 nm) for 15 min at 4 ^o^C.

Alternatively, the geranylated tRNA incubated with NBS-oxidated probe (i.e., **7g**) for 10 min on icy-water bath**.** Then the *E. coli* lysate was co-incubated with the mixture at 4 ^o^C for 30 min, and was irradiated with UV light (365 nm) for 15 min at 4 ^o^C.

(2) The mixture was incubated at 4 ^o^C while kept rotating overnight, and the washed Beads was added in the sample, the beads-protein mixture was incubated and rotated for 4 h. Then the unbounded protein was washed out on magnetic stand. Eventually, the sample was detected by SDS-PAGE.

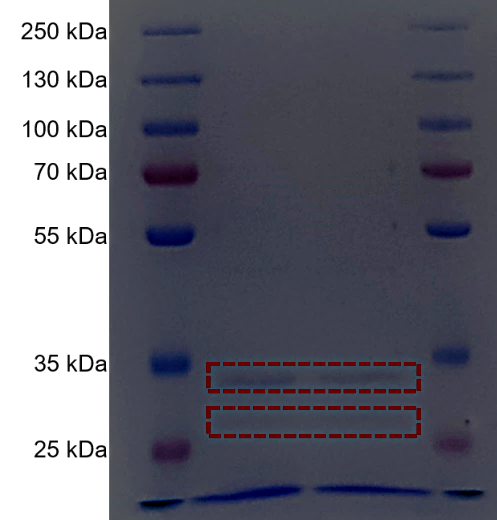

**Figure S72.** Second experiment result.

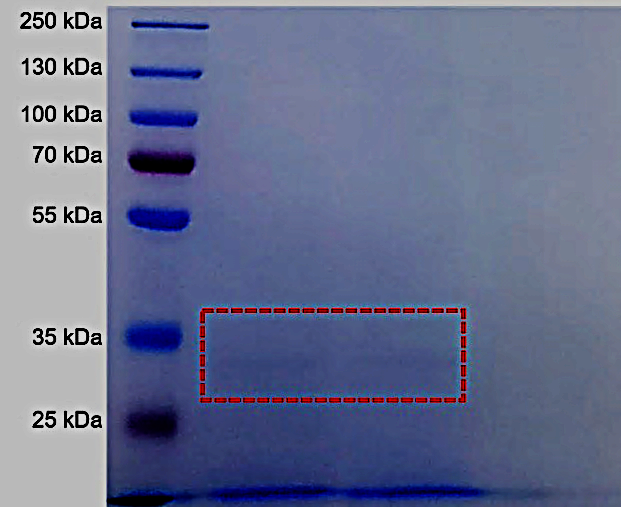

**Figure S73.** Repeated experiment result.

**Procedure of proteomics analysis of tRNA geranylation reader proteins.**

**Technique route**

In this project, a series of cutting-edge technologies, such as high-performance liquid chromatography (HPLC) classification technology and mass-spectrometry-based proteomics technology, were combined to detect and analyze the protein modification of a sample. The technical route is as follows:

**Trypsin hydrolysis**

Cut the film into 1 mm^3^ rubber blocks with surgical blade and pack them in clean EP tube; wash the rubber blocks with water for 2-3 times, add 200 *μ*l decolorizing solution (35% acetonitrile ACN and 50 mM NH_4_HCO_3_), decolorize with slight vibration, add 70% ACN (200 *μ*l) for dehydration, add DTT (200 *μ*l, 25 mM, dithiothreitol), place at r.t. for 1 h, discard the supernatant; add NH_4_HCO_3_ (200 *μ*l, 50 mM), IAM (iodoacetamide), standing at r.t. in dark for 10 min, discarding the supernatant, adding 200 *μ*l 70% ACN for dehydration, and finally adding trypsin at the mass ratio of 1: 50 (trypsin: protein), enzymolysis at 37°C overnight. Then the trypsin was added in the mass ratio of 1: 100 (trypsin: protein), and the enzymatic hydrolysis was continued for 4 h.

**HPLC classification**

The peptides were graded by high pH reverse HPLC on an Agilent 300 extend C_18_ column (5.0 *μ*m particle size, 4.6 mm inner diameter, 250 mm length). The operation is as follows: peptide grading gradient is 8% - 32% acetonitrile, pH = 9, and the combined components are vacuum freeze-dried for subsequent operation.

**LC-MS analysis**

1. Each pre-separated component was dissolved in the liquid phase, centrifuged at 20,000×*g* for 5 min, and the supernatant was transferred to the upper sample bottle;
2. Each component was analyzed by LC-MS for 1 h, and the loading amount was about 2.0 *μ*g.

**Table S10**. Parameter setting of nasol ultra high-performance liquid in liquid chromatography mass spectrometry.

| Chromatograph model | Thermo EASY-nLC 1200 |
| --- | --- |
| Model of capture column | Thermo Acclaim PepMap 100 C_18_ column, 2 *μ*m, 75 *μ*m × 20 mm |
| Model of analytical column | Thermo Acclaim PepMap RSLC C_18_ column, 2 *μ*m, 75 *μ*m × 500 mm |
| A / B solution | Thermo Acclaim PepMap 100 C_18_ column, 2 *μ*m, 75 *μ*m × 20 mm |
| Loading solution | Thermo Acclaim PepMap RSLC C_18_ column, 2 *μ*m, 75 *μ*m × 500 mm |
| Column oven temperature | 100% H_2_O, 0.1% FA/80% ACN, 0.1% FA |
| Current Speed | 250 nl/min |
| Gradient program | 0-6 min, 2-10% B; 6-51 min, 10-20% B; 51-58 min, 20-80% B; |
|  | 58-62 min, 80% B; 62-63 min, 80-2% B; 63-70 min, 2% B. |

**Table S11.** Parameter setting of quadrupole orbital trap mass spectrometry in liquid chromatography mass spectrometry system.

| Mass spectrometer model | Thermo Scientific Q Exactive HFX |
| --- | --- |
| Spray voltage | 2.2 kV |
| Capillary temperature | 280℃ |
| MS1 scan range | 350-1800 m/z |
| MS1 resolution | 60000 |
| MS1 AGC | 3E6 |
| MS1 maxIT | 20 ms |
| MS2 scan range | from 100 m/z |
| MS2 resolution | 30000 |
| MS2 AGC | 5E4 |
| MS2 maxIT | 200 ms |
| Parent ion selection | Top15 |
| Selection threshold of parent ion | 1E4 |
| Split window | 1.4 m/z |
| Fragmentation energy | HCD, 25% NCE |
| Dynamic exclusion | On, 30s |

**Database Search**

The secondary mass spectrum data were retrieved by pFind (v3.1). Parameter setting: the target protein database, added the reverse library to calculate the false positive rate (FDR) caused by random matching, and added the common pollution Library in the database to eliminate the influence of contaminated protein in the identification results; the enzyme digestion method was set to trypsin/p; the number of missing sites was set to 2; the minimum length of peptide was set to 7 amino acid residues; the maximum modification number of peptide In the first search and main search, the mass error tolerance of primary parent ion is 20 ppm and 5 ppm respectively, and the mass error tolerance of secondary fragment ion is 0.02 Da. The FDR of protein identification, and PSM identification were set to 1%.

**(1) Protein extraction**

Serum samples:

Firstly, the cellular debris of serum sample was removed by centrifugation at 12,000×*g* at 4°C for 10 min. Then, the supernatant was transferred to a new centrifuge tube. The top 12 high abundance proteins were removed by Pierce™ Top 12 Abundant Protein Depletion Spin Columns Kit (Thermo Fisher). Finally, the protein concentration was determined with BCA kit according to the manufacturer’s instructions.

**(2) Trypsin digestion**

For digestion, the protein solution was reduced with dithiothreitol (5 mM) for 30 min at 56°C and alkylated with iodoacetamide (11 mM) for 15 min at r.t. in darkness. The protein sample was then diluted by adding TEAB (100 mM) to urea concentration less than 2.0 M. Finally, trypsin was added at 1:50 trypsin-to-protein mass ratio for the first digestion overnight and 1: 100 trypsin-to-protein mass ratios for a second 4 h-digestion.

**(3) TMT/ iTRAQ labeling**

After trypsin digestion, peptide was desalted by Strata X C_18_ SPE column (Phenomenex) and vacuum-dried. Peptide was reconstituted in TEAB (0.5 M) and processed according to the manufacturer’s protocol for TMT kit/ iTRAQ kit. Briefly, one unit of TMT/ iTRAQ reagent were thawed and reconstituted in acetonitrile. The peptide mixtures were then incubated at r.t. for 2 h and pooled, desalted and dried by vacuum centrifugation.

**(4) HPLC fractionation**

The tryptic peptides were fractionated into fractions by high pH reverse-phase HPLC using Thermo Betasil C_18_ column (5 *μ*m particles, 10 mm i.d., 250 mm length). Briefly, peptides were first separated with a gradient of 8% to 32% acetonitrile (pH 9.0) over 60 min into 60 fractions. Then, the peptides were combined into fractions and dried by vacuum centrifuging.

**(5) Affinity enrichment (optional)**

Bio-material-based PTM enrichment (for phosphorylation): Peptide mixtures were first incubated with IMAC microspheres suspension with vibration in loading buffer (50% acetonitrile/ 6% trifluoroacetic acid). The IMAC microspheres with enriched phosphopeptides were collected by centrifugation, and the supernatant was removed. To remove nonspecifically adsorbed peptides, the IMAC microspheres were washed with 50% acetonitrile/ 6% trifluoroacetic acid and 30% acetonitrile/ 0.1% trifluoroacetic acid, sequentially. To elute the enriched phosphopeptides from the IMAC microspheres, elution buffer containing 10% NH_4_OH was added and the enriched phosphopeptides were eluted with vibration. The supernatant containing phosphopeptides was collected and lyophilized for LC-MS/ MS analysis.

**(6) LC-MS/MS analysis**

The tryptic peptides were dissolved in 0.1% formic acid (solvent A), directly loaded onto a home-made reversed-phase analytical column (15-cm length, 75 *μ*m i.d.). The gradient was comprised of an increase from 6% to 23% solvent B (0.1% formic acid in 98% acetonitrile) over 26 min, 23% to 35% in 8 min and climbing to 80% in 3 min then holding at 80% for the last 3 min, all at a constant flow rate of 400 nL/ min on an EASY-nLC 1000-UPLC system.

The peptides were subjected to NSI source followed by tandem mass spectrometry (MS/MS) in Q Exactive TM Plus (Thermo) coupled online to the UPLC. The electrospray voltage applied was 2.0 kV. The m/z scan range was 350 to 1800 for full scan, and intact peptides were detected in the Orbitrap at a resolution of 70,000. Peptides were then selected for MS/ MS using NCE setting as 28 and the fragments were detected in the Orbitrap at a resolution of 17,500. A data-dependent procedure that alternated between one MS scan followed by 20 MS/ MS scans with 15.0s dynamic exclusion. Automatic gain control (AGC) was set at 5E4. Fixed first mass was set as 100 m/z.

**(7) Database search**

The resulting MS/ MS data were processed using Maxquant search engine (v.1.5.2.8). Tandem mass spectra were searched against the defined database concatenated with reverse decoy database. Trypsin/ P was specified as cleavage enzyme allowing up to 4 missing cleavages. The mass tolerance for precursor ions was set as 20 ppm in First search and 5 ppm in Main search, and the mass tolerance for fragment ions was set as 0.02 Da. Carbamidomethyl on Cysteine was specified as fixed modification and the defined modification and oxidation on Met were specified as variable modifications. FDR was adjusted to < 1% and minimum score for modified peptides was set > 40.

**Table S12.** Summary of reagents and solvents.

| **Reagents** | **Vendors** |
| --- | --- |
| Trypsin | Promega |
| Acetonitrile | Fisher Chemical |
| Trifluoroacetic acid | Sigma-Aldrich |
| Formic acid | Fluka |
| Iodoacetamide | Sigma |
| Dithiothreitol | Sigma |
| Urea | Sigma |
| dd H_2_O | Fisher Chemical |

**Table S13.** Summary of the proteomic analysis.

| **Protein identifications** | **Numbers of PSMs** | **Note** |
| --- | --- | --- |
| *infC* | 233 |  |
| *ECBD_0490* | 133 |  |
| *rpsC* | 41 | ******* |
| *rplF* | 23 | ***** |
| *rplD* | 16 |  |
| *rpsD* | 11 | ****** |
| *ECBD_4279* | 10 |  |
| *rpsL* | 4 |  |
| *ECBD_1987* | 4 |  |
| *rplC* | 3 | ****** |
| *ECBD_4099* | 2 |  |
| *ECBD_3170/ ECBD_2242/ flhA/ECBD_1207 /ECBD_3065 /ECBD_2143/ ECBD_1083/ ECBD_0273/ mltG* | 2 | ***** |
| *ECBD_1207* | 1 |  |
| *serS* | 1 |  |
| *ECBD_3170* | 1 |  |

**Table S14.** Summary of the proteomic analysis results.

| **Entries** | **Gene name** | **Note** |
| --- | --- | --- |
| Translation initiation factor IF-3 | *infC* |  |
| 50S Ribosomal protein | *rplC, rplD, rplF* |  |
| 30S Ribosomal protein | *rpsC, rpsD, rpsL* |  |
| Endolytic transglycosylase | *mltG* |  |
| Serine--tRNA ligase | *serS* |  |
| Hypothetical protein | *ECBD* series |  |

**Table S15.** Three-headed probe (**7g**).

| **Protein Identifications** | **Numbers of PSMs** | **Note** |
| --- | --- | --- |
| *rplB* | 51 |  |
| *rbsB* | 28 |  |
| *rpsC* | 13 | ******* |
| *ompA* | 7 |  |
| *rpmA* | 6 |  |
| *rpsD* | 6 | ****** |
| *ompF* | 5 |  |
| *mdh* | 5 |  |
| *gpmA* | 4 |  |
| *rplC* | **4** | ****** |
| *slyD* | **4** |  |
| *tsf* | **3** |  |
| *clpX* | **2** |  |
| *nuoB* | **2** |  |
| *rplR* | **2** |  |
| *phoE / ompF / ompN* | **1** |  |
| *ybaT / febD / mltG / ydbH / yddA / flhA / hyfB / yfiR / yhhN / ydgU / yfjD* | **1** | ***** |
| *mltG* | **1** | ***** |
| *mltG / yddA* | **1** | ***** |
| *fabl* | **1** |  |
| *ansP* | **1** |  |
| *yebC* | **1** |  |
| *motB/ gnsA* | **1** |  |
| *yhdH* | **1** |  |
| *rplF* | **1** | ***** |
| *hupA* | **1** |  |
| *ynaA* | **1** |  |
